# Identification of spacer and protospacer sequence requirements in the *Vibrio cholerae* Type I-E CRISPR/Cas System

**DOI:** 10.1101/2020.08.09.243105

**Authors:** Jacob Bourgeois, David W. Lazinski, Andrew Camilli

## Abstract

The prokaryotic adaptive immune system CRISPR/Cas serves as defense against bacteriophage and invasive nucleic acid. A Type I-E CRISPR/Cas system has been detected in classical biotype isolates of *Vibrio cholerae*, the causative agent of the disease cholera. Experimental characterization of this system revealed a functional immune system that operates using a 5’-TT-3’ protospacer-adjacent motif (PAM) for interference. However, several designed spacers against the 5’-TT-3’ PAM do not interfere as expected, indicating further investigation of this system is necessary. In this study, we identified additional sequence requirements of a pyrimidine in the 5’ position of the spacer and purine in the complementary position of the protospacer using 873 unique spacers and 2267 protospacers mined from CRISPR arrays in deposited sequences of *V. cholerae*. We present bioinformatic evidence that during acquisition the protospacer purine is captured in the prespacer and that a 5’-RTT-3’ PAM is necessary for spacer acquisition. Finally, we demonstrate experimentally that a 5’-RTT-3’ PAM is necessary for CRISPR interference by designing and manipulating spacer and cognate PAMs in a plasmid conjugation assay and discover functional consequences of base pairing with the 5’ spacer pyrimidine in spacer efficacy.

**Importance:** Bacterial CRISPR/Cas systems provide immunity by defending against phage and other invading elements. A thorough comprehension of the molecular mechanisms employed by these diverse systems will improve our understanding of bacteriophage-bacterial interactions and bacterial adaptation to foreign DNA. The *Vibrio cholerae* Type I-E system was previously identified in an extinct classical biotype and was partially characterized for its function. Here, using both bioinformatic and functional assays, we extend that initial study. We have found that the Type I-E system still exists in modern strains of *V. cholerae*. Furthermore, we defined additional sequence elements in both the CRISPR array and in target DNA that are required for immunity. CRISPR/Cas systems are now commonly used as precise and powerful genetic engineering tools. Knowledge of the sequences required for CRISPR/Cas immunity is a prerequisite for the effective design and experimental use of these systems. Our results greatly facilitate the effective use of one such system. Furthermore, we provide a publicly available script that assists in the detection and validation of CRISPR/Cas immunity requirements when such a system exists in any bacterial species.

## Introduction

Clustered regularly interspaced short palindromic repeats (CRIPSR) and CRISPR-associated genes (*cas*) comprise an adaptive immune system found in many bacteria and archaea that protect cells from invasion by foreign nucleic acid (1). These CRISPR/Cas systems consist of several *cas* genes that encode proteins for adaptation and immunity alongside an array of short repeat sequences alternating with sequences derived from foreign invader, termed spacers. CRISPR targets and degrades invading nucleic acid in a sequence-specific manner. Most systems protect against DNA, though rarer types also target RNA (2, 3). In adaptation, the CRISPR/Cas system acquires and stores novel invading DNA sequences as spacers in its array. Novel spacers are integrated at the 5’ end of the array immediately downstream from the leader-proximal repeat (1, 4). In immunity, the CRISPR locus, which contains recorded spacers, is first transcribed and processed to yield CRISPR RNAs (crRNAs). On recognition by a crRNA of a complementary DNA sequence called the protospacer, the target DNA is cleaved, thereby providing immunity.

There is substantial diversity among CRISPR/Cas systems in architecture, protein composition, target affinity, and mechanisms of interference and adaptation. Systems are broadly classified into multi-subunit effector complexes in Class I, and single-unit effectors in Class II (3). These classes are further organized into several types and subtypes by virtue of genetic organization and Cas protein composition (3, 5, 6). Aside from complementarity between spacer and protospacer, CRISPR/Cas systems possess an additional sequence requirement adjacent to the protospacer called a protospacer-adjacent motif, or PAM (7, 8). The PAM size and location varies by CRISPR system and is typically a 2-5 bp sequence that may be located upstream or downstream of the target (6, 9, 10). An intact PAM is essential for acquisition and interference; mutations in this motif provide escape from CRISPR-mediated immunity (11). In the CRISPR array, the repeat sequence that is adjacent to the spacer lacks a PAM. This absence prevents autoimmunity, as the crRNA is unable to cleave the DNA template from which it is transcribed.

*V. cholerae* is a Gram-negative facultative bacterium that is the causative agent of cholera (12, 13). The seventh, most recent pandemic began in 1961, caused by *V. cholerae* serotype O1 biotype El Tor that replaced the classical biotype of previous pandemics (12, 13). A Type I-E CRISPR/Cas system has recently been described in the now extinct classical biotype of *V. cholerae*. The genomic island GI-24, found in the classical strain O395, was predicted to encode Cas proteins (14). Box et al. introduced GI-24 into the El Tor biotype, and demonstrated experimentally that the putative CRISPR/Cas system is functional, and artificial induction of the system provides resistance against virulent bacteriophage under laboratory conditions (15). Based on sequence alignment of 33 cognate protospacers from spacers mined from five sequenced classical isolates and analysis of bacteriophage escape mutants, the PAM for the system was determined to be 5’-TT-3’, located immediately downstream of the protospacer.

A useful application of the CRISPR/Cas system is the creation of insertions, deletions, and point mutations in virulent bacteriophages, manipulations that can be quite difficult using conventional genetic engineering techniques (16, 17). By introducing spacers against wild-type sequences into a host strain containing *cas* genes, mutant bacteriophage are enriched as they escape CRISPR/Cas immunity. Such escape mutants can be precisely edited by the provision of homologous recombination templates that contain the desired alteration (16, 17). This principle has been used to edit bacteriophages in several model systems, such as *Escherichia coli* and *Streptococcus thermophilus*, with editing efficiencies as high as 100% (18, 19). In the characterization of the Type I-E CRISPR/Cas system in *V. cholerae*, Box et al. placed a minimal CRISPR array onto a plasmid such that engineered spacers may be introduced easily for editing of bacteriophage (15). However, the degree of interference varied widely between engineered spacers. Half of spacers targeting open reading frames in the virulent vibriophage ICP1 only offered ten-fold protection, even with induction of both *cas* genes and spacer crRNA, whereas other spacers produced efficiencies of plaquing as low as 10^-5^ (15). This unpredictability in the efficacy of engineered spacers suggests there exist further unknown requirements for predictable interference in the *V. cholerae* CRISPR/Cas system.

Knowledge of the parameters for interference is crucial in the design of effective spacers for CRISPR/Cas. Furthermore, dissection of CRISPR/Cas systems expands our knowledge of the broader molecular mechanisms of CRISPR adaptation and interference. In this study, we greatly expand the database of spacers and cognate protospacers by mining deposited sequences of *V. cholerae* with identical Type I-E CRISPR arrays and identify a conserved pyrimidine on the 5’ end of naturally acquired spacers that is derived from the incoming prespacer. Using the established plasmid-based CRISPR system, we experimentally determined that this pyrimidine is not required for efficient interference but its complementary purine in target DNA is required and therefore propose a 5’-RTT-3’ PAM. Finally, we investigate the role of base pairing at this position in relation to the efficiency of interference.

## Results

### Analysis of CRISPR repeats in publicly available sequences of *V. cholerae* identifies a conserved 5’-pyrimidine in spacers

The original characterization of the Type I-E CRISPR/Cas system in *V. cholerae* identified 78 unique spacers and 22 corresponding protospacers across five sequenced isolates (15). To obtain more spacers and protospacers, we identified additional CRISPR-containing *V. cholerae* isolates by querying NCBI databases for strains containing the 28 bp *V. cholerae* O395 CRISPR repeat (5’-GTCTTCCCCACGCAGGTGGGGGTGTTTC-3’). In total, 1671 repeats were detected over 66 sequence accessions containing 45 unique strains isolated over a wide geographical area (Figure 1A, Supplementary Table 1). Of note, several isolates were obtained as recently as 2018, indicating that the CRISPR/Cas system first identified in the extinct classical biotype itself is not extinct and continues to influence modern *V. cholerae* genomes. From these samples, spacers and protospacers were extracted using a custom script written using the programming language Python (see Materials and Methods). To ensure validity, only spacers that were no longer than 50 nucleotides, were flanked by perfect repeat sequences, and did not contain ambiguous nucleotides were considered. Only protospacers that were at least 93% identical over 96% of the corresponding spacer are considered. Overall, our method obtained 873 unique spacers (Figure 1A, Supplementary Table 2). The median length of spacer sequence was 33 bp, consistent with the previous characterization (15). Several spacers are shared among analyzed isolates – a spacer targeting vibriophages Rostov 7 and X29 is seen in eight separate isolates. However, nearly 75% of spacers are unique to a single strain, suggesting separate encounters and acquisition events (Supplementary Table 3).

**Figure 1.**
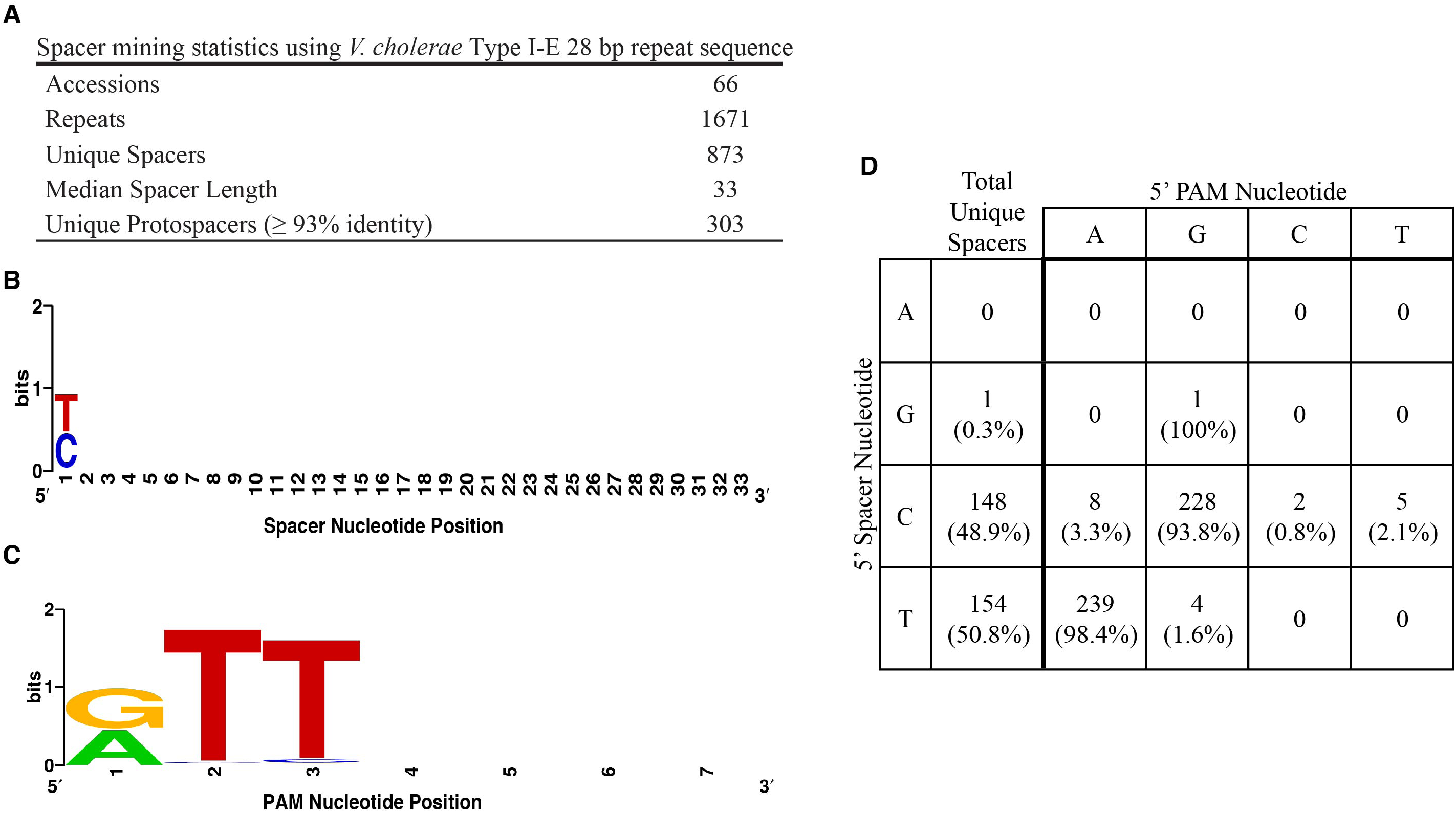
Analysis of spacer and protospacer-adjacent motifs mined from deposited *V cholerae* sequences. **A.** Overview of spacer mining statistics obtained from detection of repeats. **B.** Sequence alignment of unique spacers derived from between perfect repeats. Shown in the 5’ to 3’ direction. For spacers longer than 33 bp, only the first 33 bp were used in the creation of the Weblogo. **C.** Sequence alignment of the PAM retrieved immediately downstream from protospacers. The first position of the PAM is aligned with the 5’ position of the spacer. Shown in the 5’ to 3’ direction. **D.** Frequency table of 5’ spacer and 5’ PAM nucleotide identity for 487 unique spacer-protospacer pairs. Total unique spacers refers to the number of unique spacers with detected cognate protospacers that begins with the specified nucleotide.

Of the 873 unique spacers, 303 (34%) matched to at least one valid protospacer target (Figure 1A, Figure 1D, Supplementary Table 3). In accordance with the role of CRISPR as a bacterial immune system, several protospacers from many diverse bacteriophages were identified, including vibriophages fs1 and fs2, kappa, CP-T1, among others, for a total of 2267 protospacers. In several cases, for individual unique spacers we found nucleotide polymorphisms between cognate protospacer sequences obtained from different organisms. Overall, we obtained 487 unique spacer-protospacer pairs (Table 1D, Supplementary Table 4). Additional protospacers mapped to prophages found in other *Vibrio* isolates, environmental broad-host range replicons such as repSD172 (20), or identical spacers in sequenced CRIPSR arrays. In total, our analysis expanded the list of unique spacers acquired naturally by more than ten-fold and identified hundreds of novel target protospacers and genetic elements. Furthermore, the recent isolation dates of many isolates suggest that CRISPR/Cas is still an active force in *V. cholerae* evolution. However, no isolates of the *V. cholerae* El Tor biotype, the causative agent of the current seventh cholera pandemic (12), were detected.

**Table 1.**
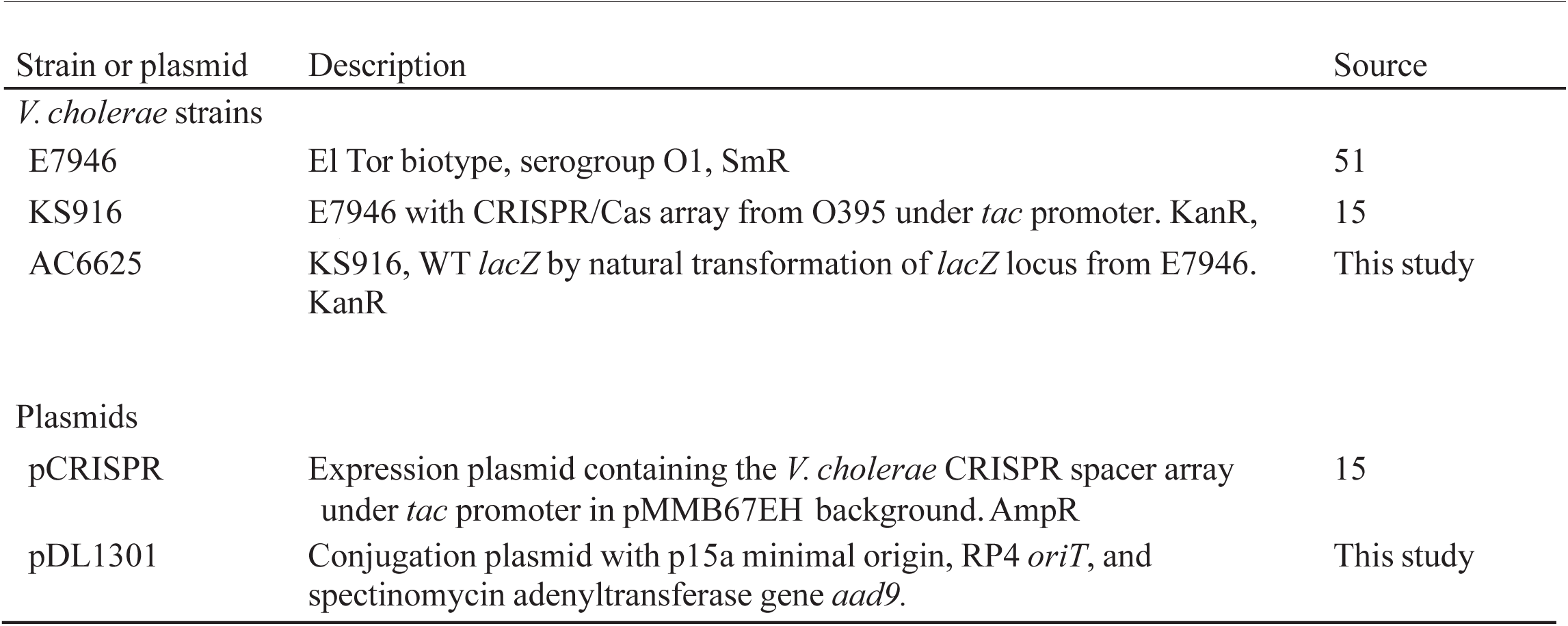
Bacterial strains and plasmids used in this study spectinomycin adenyltransferase gene *aad9*.

Upon further inspection of mined data, nearly every extracted spacer (99.7% of 303 unique spacers) began with a pyrimidine in the 5’ position (Figure 1D, Supplementary Table 2). Alignment of unique spacer sequences confirmed the 5’ position of the spacer is a conserved pyrimidine (Figure 1B). To detect the presence of the protospacer-adjacent motif (PAM), we aligned mined protospacers, starting from the 3’-most nucleotide of the protospacer that aligns with the 5’ spacer nucleotide. Alignment revealed a strong 5’-RTT-3’ motif, where the conserved purine is complementary to the 5’ pyrimidine of the spacer (Figure 1C). Additionally, examination of 483 unique spacer-protospacer pairs reveals that spacers with a 5’ cytosine have a cognate 5’ PAM guanine 93.8% of the time, and 98.4% of those matching 5’ thymine spacers possess a 5’ adenine PAM. This near-perfect complementarity observed between the purine of the PAM and the pyrimidine of the first position of the spacer provides evidence that the purine of the PAM is captured during spacer acquisition. The small number of cases where complementarity is lacking presumably results from mutational escape from CRISPR targeting. Therefore, our bioinformatic results suggest a role for the purine in capture and adaptation, and thus the functional PAM may be 5’-RTT-3’ instead of 5’-TT-3’.

### The conserved spacer pyrimidine and complementary protospacer purine is necessary for CRISPR interference

PAM sequences are required for both acquisition and interference (9, 10, 21). Thus far we have provided bioinformatic evidence consistent with a 5’-RTT-3’ PAM in the target DNA being required for acquisition. To experimentally determine the importance of the 5’ spacer nucleotide and 5’-RTT-3’ PAM in CRISPR interference, we designed eight independent spacers targeted against the *aad9* spectinomycin adenyltransferase gene, four of which have a 5’-pyrimidine against a 5’-RTT-3’ PAM, and four with a 5’-purine against a 5’-YTT-3’ PAM (Table 2). We introduced these spacers into a functional plasmid-based inducible CRISPR/Cas targeting system, pCRISPR (15). These eight targeting plasmids were then introduced into AC6625, a *V. cholerae* El Tor strain containing inducible *cas* genes from classical isolate O395. We then tested the efficiency of conjugation of pDL1301, a plasmid containing the *aad9* gene, into each targeting strain. The *trp*-*lac* fusion promoter (*tac*) is used to drive expression of both crRNA and Cas proteins in the targeting strains (15). The *tac* promoter is known to function at modest levels in the absence of IPTG and at very high levels upon induction (22). Initial experiments were therefore performed in the absence of IPTG. The conjugation efficiency of pDL1301 into a strain with a non-targeting spacer (5’-TGAGACCAGTTCTCTCGGAAGCTCAAAGGTCTC-3’) was obtained as a control. All four 5’-pyrimidine spacers provided appreciable interference against their 5’-RTT-3’ PAMs, three of which had conjugation efficiencies 10^3^-10^4^-fold lower than non-targeting control (Figure 2). In contrast, all four 5’-purine spacers had efficiencies no better than control at targeting 5’-YTT-3’ PAMs. These results demonstrate that either a 5’-pyrimidine in the spacer or a 5’-purine in the PAM, or both, are needed to elicit interference.

**Figure 2.**
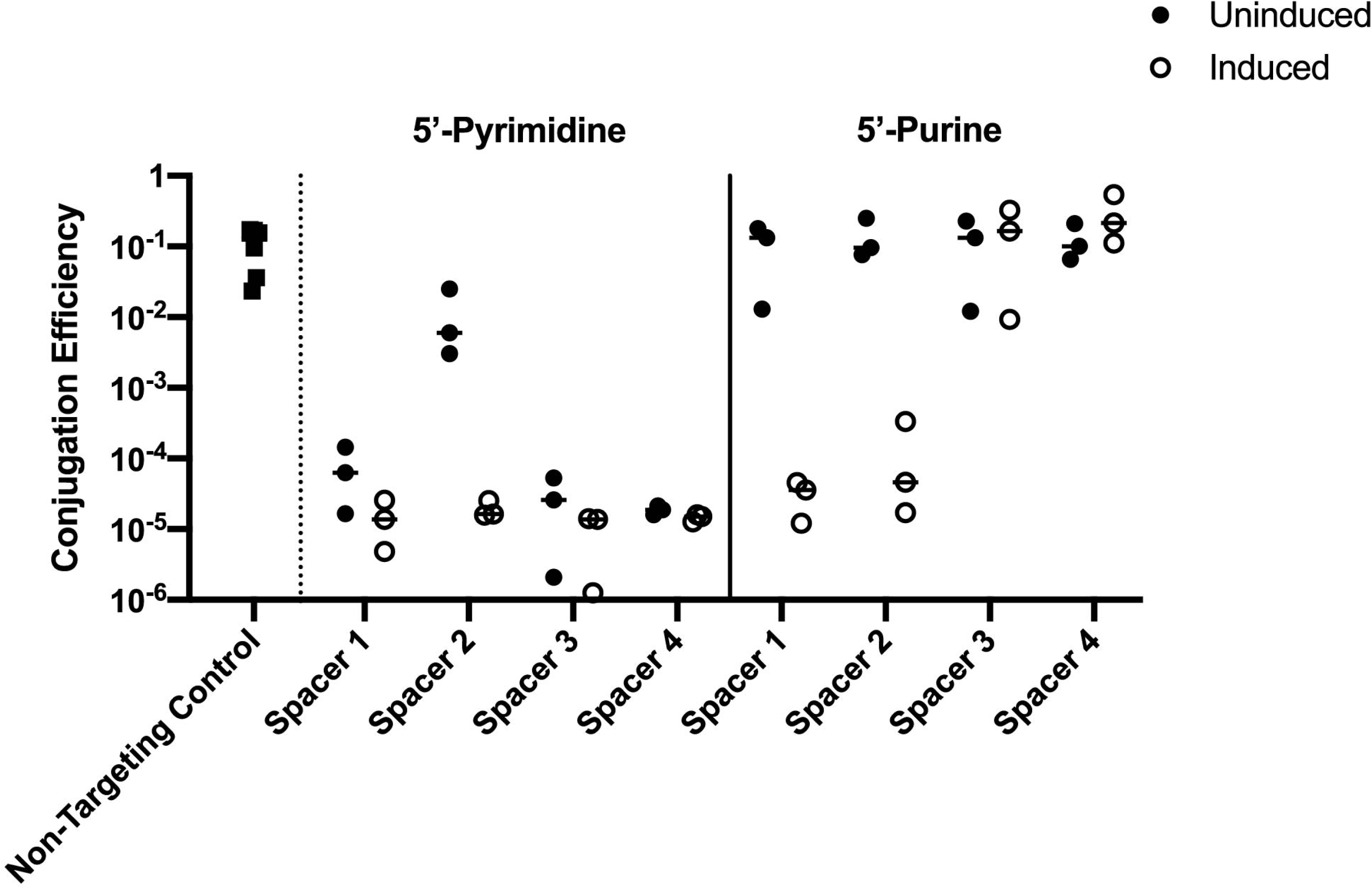
Targeting activity of 5’-pyrimidine and 5’-purine spacers against 5’-RTI-3’ and 5’-YTI-3’ PAMs in *aad9,* respectively. Conjugation efficiency of mating pDL1301, a conjugatable plasmid containing the *aad9* gene, into El Tor *V cholerae* possessing Cascade and engineered targeting spacers on a plasmid-based pCRISPR (solid circles). The non-targeting control contains pCRISPR encoding a spacer with no homology to pDL1301 (solid squares, left). Open circles show effect of induction of Cascade and crRNA with 1OOµM IPTG on interference. Data represent conjugation efficiency of plasmid into three independently obtained *V cholerae* exconjugates.

**Table 2.**
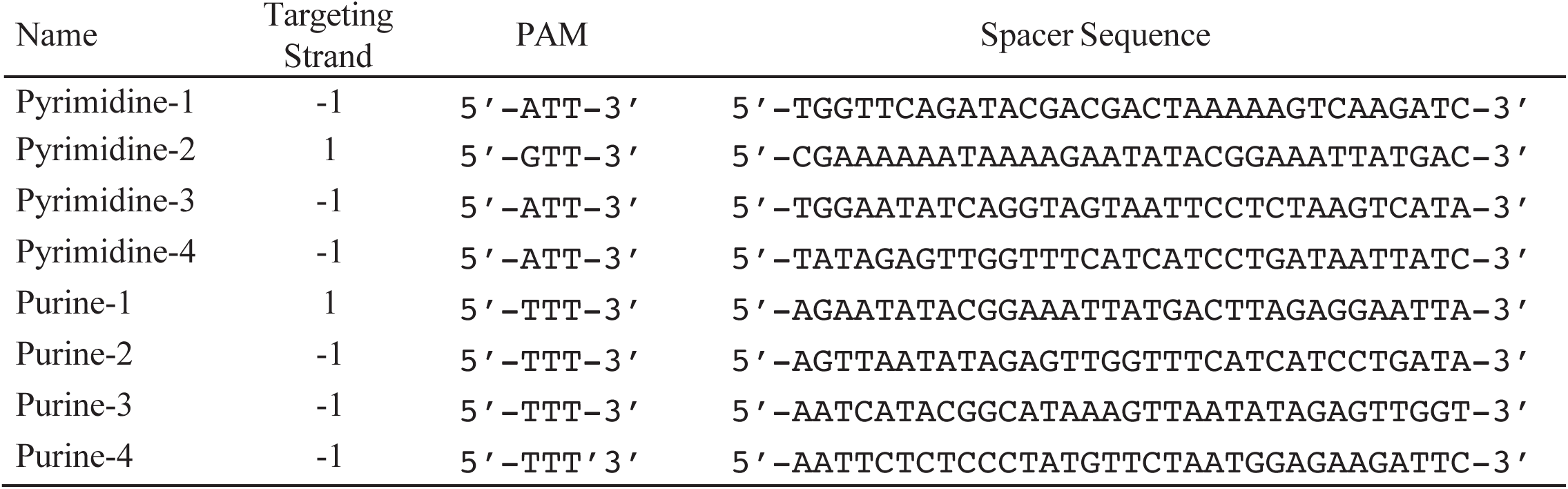
Engineered spacers against the *aad9* gene in target plasmid

To investigate the effect of induction of targeting machinery on conjugation efficiency, conjugation was also performed in the presence of IPTG on the selection plate. Induction was able to rescue the weak targeting activity of 5’-pyrimidine-spacer 2 to the same level as other 5’-pyrimidine spacers (Figure 2). Additionally, induction was able to elicit targeting in 5’-purine spacers 1 and 2, but not 3 and 4. These results suggest that induction of both Cascade and crRNA may rescue poorly targeting spacers, a phenomenon observed in other CRISPR systems (23, 24).

### Transversion mutation at the +1 targeting position reverses interference efficacy

The results in Figure 2 show that a pyrimidine in the +1 crRNA spacer position with a complementary purine in the protospacer is necessary for efficient protospacer targeting, which is show schematically in Figure 3A. However, several spacer attributes may influence targeting, including GC content, crRNA stability and processing, and other poorly understood parameters (25–27). To control for confounding intrinsic spacer attributes, the 5’ nucleotide in each pCRISPR targeting construct was mutated from a pyrimidine to a purine, or vice-versa (crRNA*). Then, the target *aad9* gene in donor plasmid pDL1301 was individually and separately silently modified so that base pairing is restored at the +1 position with each new targeting spacer (*aad9*\*). Thus, each *aad9* mutation additionally changes the putative targeting motif from 5’-RTT-3’ to 5’-YTT-3’, and vice versa. The new spacers, PAMs, and corresponding *aad9* mutations are listed in Table 3. The conjugation assay was then done on each matching pair of mutated pDL1301 and targeting strain and compared to data from Figure 2 (Figure 3B). Switching a 5’ pyrimidine spacer/ 5’ purine PAM pair to a purine/pyrimidine pair abolished targeting. In contrast, switching a 5’ purine spacer/5’ pyrimidine PAM pair to a pyrimidine/purine pair provided modest increases in interference for 5’-purine spacers 1 and 4, and substantial increases for spacers 2 and 3. In total, these results confirm the necessity of a 5’-pyrimidine in the +1 spacer position and/or an opposite complementary purine in the target for efficient CRISPR interference activity, independent of individual spacer characteristics.

**Figure 3.**
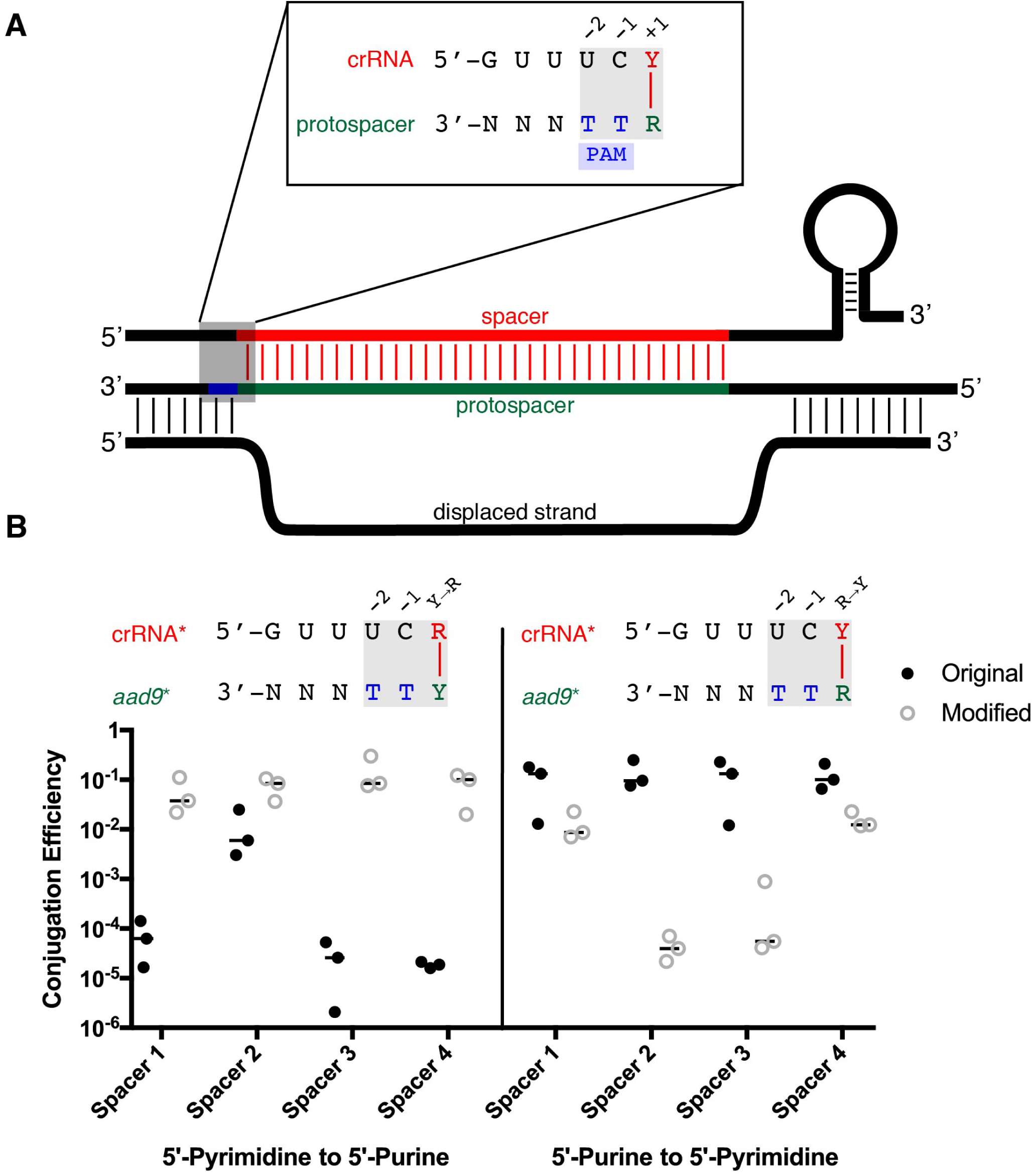
Mutating the 5’ spacer nucleotide and the corresponding *aad9* target PAM reverses crRNA targeting efficacy. **A.** Schematic of crRNA strand invasion and protospacer binding in *V cholerae.* The published protospacer-adjacent motif (PAM), 5’-TI-3’, is shown in blue at the -1 and -2 positions. The 5’ spacer nucleotide is highlighted in red at the +1 position, where it complements with the protospacer in green. **B.** Effect of transversion mutation at the +1 site. The *aad9* gene in pDL1301 was silently mutated at the +1 site from a purine to a pyrimidine, or vice versa, at targeting sites to create eight variations of donor plasmid *(aad9*).* The corresponding spacer in targeting strains was then changed to match each new donor plasmid to preserve base pairing at the +1 position, creating eight new targeting strains that individually pair to its donor strain (crRNA*). Conjugation efficiency of these eight new pairs was obtained. Data for the original, unmodified conjugation (solid circles) are reproduced from Figure 2 for the sake of comparison. The crRNA-PAM diagrams above represent the modified condition. Data represent mating efficiencies of three independently obtained *V cholerae* exconjugates.

**Table 3.**
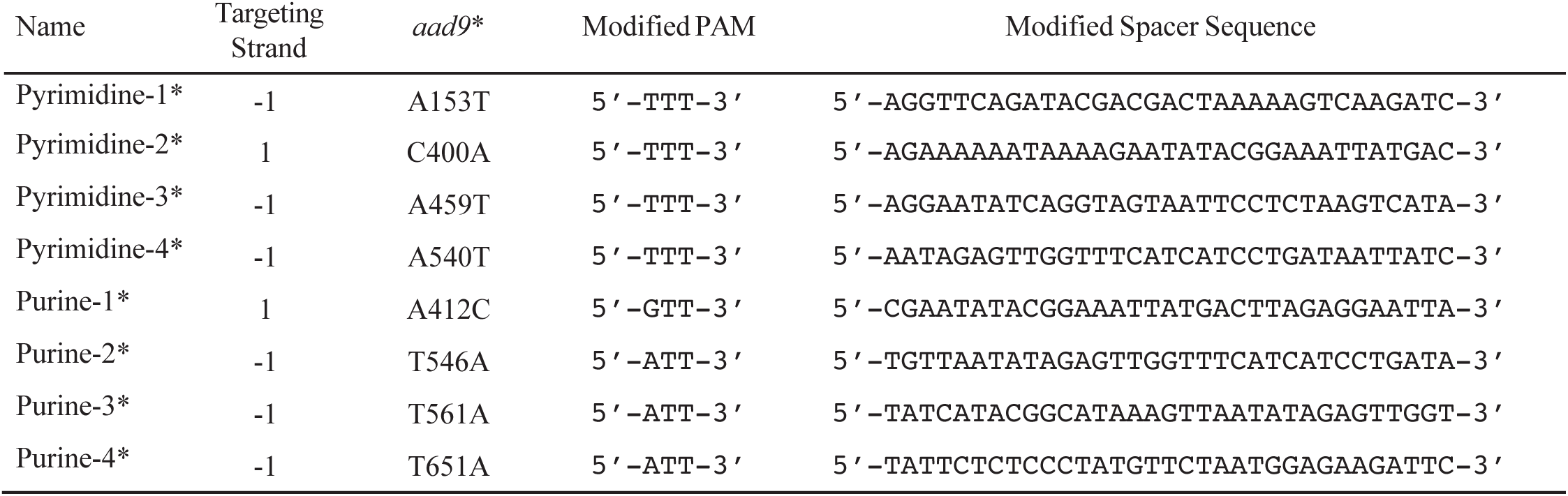
Modifications of targeting spacer and corresponding silent mutations in *aad9* gene

### Disruption of base pairing at the +1 position affects interference

Thus far, our results have shown that, for effective interference, there must be a pyrimidine in the +1 spacer position complementary to a target purine. It is unclear which sequence requirement is dominant – that is, if the conserved spacer pyrimidine occurs as a consequence of a stringent purine requirement in the PAM, or vice-versa. To independently examine the role of the 5’ spacer pyrimidine and the 5’ PAM nucleotide in interference, the unmodified pDL1301 was mated into 5’-pyrimidine mutated targeting constructs pyrimidine-1* through pyrimidine-4*, and the modified *aad9*\* pDL1301 plasmids were mated into the respective unmodified targeting constructs purine-1 through purine-4. These combinations preserve a 5’-RTT-3’ PAM in the target plasmid in all cases but disrupt base pairing at the +1 PAM position as the crRNA spacers contain 5’-purines (Figure 4A). The conjugation efficiencies of these matings were compared to previous efficiencies obtained when mating combinations preserve base pairing (Figure 4B). We observed that all constructs that target a 5’-RTT-3’ PAM irrespective of 5’ spacer identify reduced conjugation efficiency by at least 10-fold, and up to 10^4^-fold. In total, seven of eight 5’-RTT-3’ PAMs were targeted efficiently in the absence of induction while earlier, we showed that under the same conditions, none of the eight 5’-YTT-3’ PAMs were targeted efficiently (Figure 2, Figure 3B). Together, these results support the conclusion that in *V. cholerae*, the true PAM for interference is 5’-RTT-3’.

**Figure 4.**
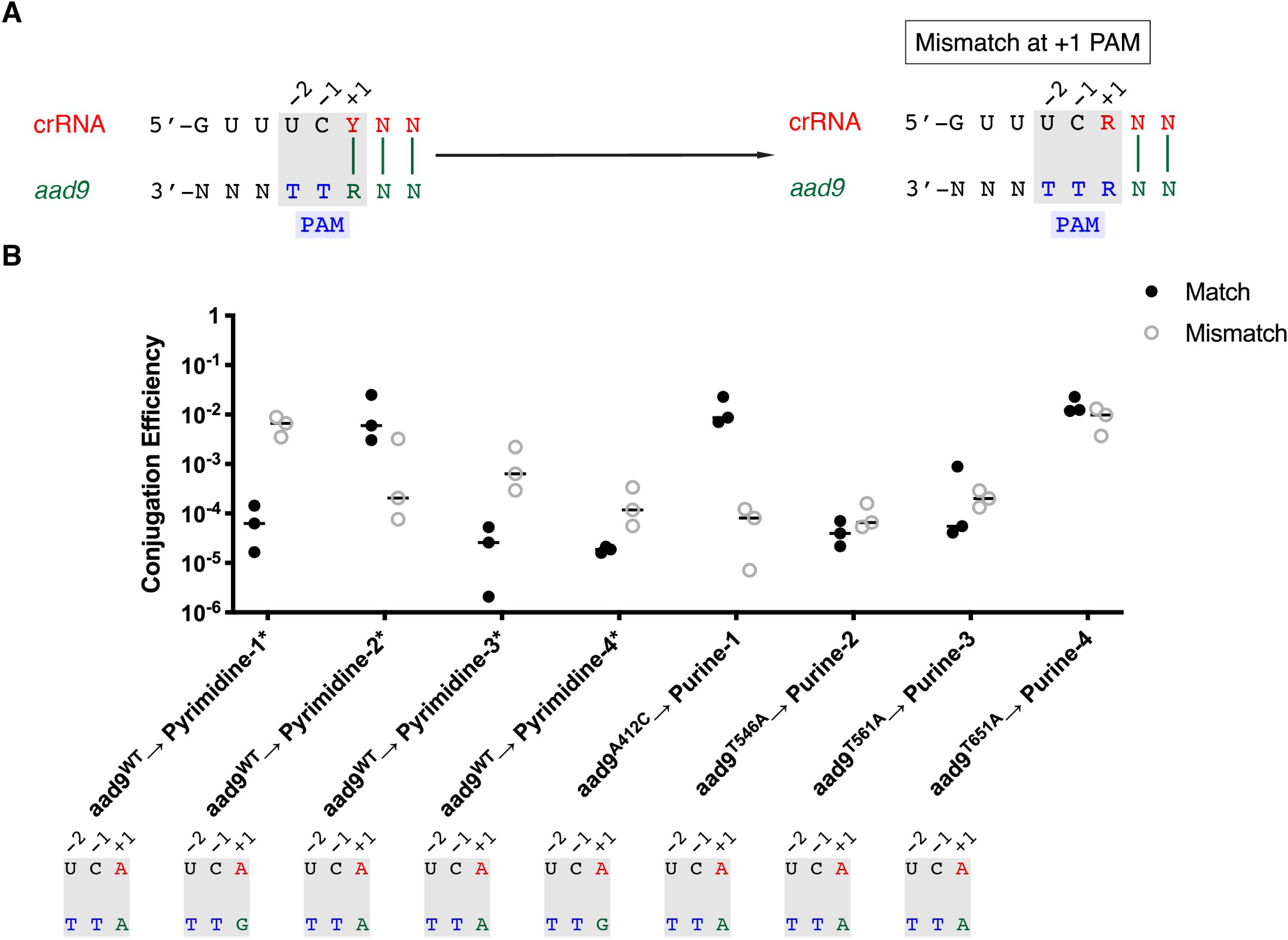
Effect of disallowing base pairing at the +1 PAM nucleotide position on interference activity. **A.** Mating modified 5’-pyrimidine* constucts against WT *aad9* (open circles, *aadfJNT)* or mating PAM-modified pDL1301 into corresponding 5’-purine spacer constructs (solid circles, *aad9*)* disrupts base pairing with PAM at+ 1 position while preserving 5’-RTT-3’ PAM. **B.** Conjugation efficiencies of matings before and after mutation resulting in mismatch. Data for the conjugation efficiencies of *aadfJNT* into unmodified pyrimidine targeting strains (solid circles, *aadrJNT)* are reproduced from Figure 2 for the sake of comparison. Data for the matched conjugation pairs of mutated *aad9* to corresponding 5’-modified purine targeting strains (open circles, *aad9*)* are reproduced from Figure 3B for the sake of comparison. The crRNA-PAM architecture for the mismatch experiment is shown below each sample. Data represent conjugation efficiencies of three independent experiments.

We found that in three of eight cases a decrease in targeting efficiency was observed when targeting a 5’-RTT-3’ PAM with a 5’ spacer purine (Figure 4B). The most extreme case was spacer pyrimidine-1*, in which a 1000-fold targeting efficiency was observed when base pairing was allowed but only a 10-fold efficiency occurred when base pairing was disallowed. In contrast, two combinations, *aad9*^WT^ → pyrimidine-2* and *aad9*^T546A^ → purine-1*, showed the opposite effect as targeting worked approximately 100-fold better when base-pairing was disallowed. One difference between these spacers and the others is that in the latter cases these spacers target a 5’-GTT-3’ PAM on the *aad9* coding strand, while in the former cases the spacers target a 5’-ATT-3’ PAM on the noncoding strand. Perhaps the 5’-GTT-3’ PAM or presence of the protospacer on the coding strand may influence the effect of base pairing with the +1 PAM position (Figure 4B, Table 2). Regardless, these results show that base-pairing between the +1 of crRNA and the PAM purine is not required for interference. Further experiments will be needed to fully understand the mechanistic importance of base pairing between the first position of the spacer and the first position of the PAM in interference.

## Discussion

The molecular history recorded in CRISPR/Cas regions of bacterial genomes is a valuable resource for researchers studying molecular epidemiology and encounters with foreign DNA. Numerous databases and web servers have been developed to detect CRISPR arrays and Cas gene clusters (28, 29). We opted for a straightforward approach to obtain spacer content from arrays across multiple isolates of *V. cholerae*. Our script allows us to automate spacer extraction, protospacer/PAM detection, and extraction of annotation information from the target regions to assist in analysis. The running parameters are easily modifiable to permit mining of spacer, protospacer, and PAM-associated information of any bacterial species with any known repeat sequence.

Horizontal transfer of mobile genetic elements has played a key role in the success of *V. cholerae* as a human pathogen. For example, genes encoding for cholera toxin are encoded by the lysogenic bacteriophage CTXΦ (30), and an integrative conjugatable element SXT provides resistance to multiple antibiotics (31). Many isolates with detected CRISPR repeats do not possess these and other key virulence factors necessary for human infection. One hypothesis is that CRISPR/Cas provides a barrier to the acquisition of advantageous traits that aid infection. CRISPR/Cas is negatively associated with virulence in *E. coli*, and negatively impacts natural transformation in *Streptococcus pneumoniae* (32–34). Consistent with this, we discovered spacers against several plasmids and filamentous CTXΦ-like prophages. We also found spacers targeting prophage genes homologous to *rstA2* and *rstB2* of RS2-CTXΦ, genes that are necessary for replication and integration of CTXΦ (35). Increased exposure in *V. cholerae* of the El Tor biotype to incoming genetic elements that provide selective advantages may outweigh any disadvantages that occur from the loss of adaptive immunity directed against predatory phages. (36, 37). Indeed, some *V. cholerae* strains have acquired specific genomic islands, termed PLEs, that inhibit replication of the ICP1 vibriophage and thereby obviate the need for CRISPR/Cas in that setting (38). Similarly, the SXT element resident in the strain that caused the cholera epidemic in Haiti contains genes highly homologous to those from a known anti-phage system (39). The CRISPR/Cas system in *V. cholerae* was thought to be limited to strains in the classical biotype. Since this biotype is thought to be extinct (12, 40, 41), it was assumed that its CRISPR/Cas system was also extinct. Here we found direct evidence to the contrary, as we identified multiple environmental strains isolated in the last three years from locations as diverse as Bangladesh, Russia and Australia, all containing the CRISPR/Cas system. Hence, the system continues to impact the evolution of *V. cholerae*.

The Type I-E CRISPR/Cas system in *E. coli* is most commonly described as having 32 bp spacer sequences flanked by 29 bp repeats in its CRISPR array, targeting a 3 bp 5’-CWT-3’ PAM (21, 42). In the crRNA of this system the terminal guanine that could potentially base pair with the cytosine of the 5’-CWT-3’ PAM was initially assumed to be part of the repeat and created by repeat duplication during acquisition (43). However, there is growing evidence that this guanine is instead captured from incoming prespacers of foreign DNA (9, 21, 44). In *V. cholerae,* a pyrimidine in the CRISPR array occupies the position analogous to the guanine in the *E. coli* system. In nearly all mined protospacers, when the array pyrimidine was a cytosine, a guanine was observed in the protospacer PAM, while a thymine in the array was almost always associated with an adenine. Furthermore, within individual CRISPR arrays, 5’-thymine and 5’-cytosine spacers are randomly distributed. Together, these results indicate that the pyrimidine does not result from repeat duplication during spacer integration and is instead protospacer-derived. Our observations in *V. cholerae* may support findings in the analogous *E. coli* system, namely that in the *E. coli* system the guanine is similarly protospacer-derived and should be considered as the first position of the spacer. By that definition, as is the case for *V. cholerae,* the system has 33 bp spacer sequences flanked by 28 bp repeats.

Protospacer-adjacent motifs (PAMs) provide at least three functions: they are required for capture and adaptation, they are required for interference, and they are required for self-discrimination (10). Here we propose that the PAM for the *V. cholerae* system is 5’ RTT 3’. The bioinformatic data presented support the importance of the 5’ purine in acquisition, while experimental data support its importance in interference. However, this nucleotide is not adjacent to the protospacer, and is instead captured as the prespacer terminal nucleotide. Thus, as base pairing is expected at this PAM purine, it is not involved in autoimmunity and the 5’-TT-3’ sequence alone must provide this function. In the majority of Type I and Type II CRISPR/Cas systems, this corresponding +1 PAM position is not retained in the captured spacer following target cleavage (8, 45–47). We propose that the *V. cholerae* and *E. coli* systems differ in that cleavage occurs within the PAM during capture, thereby incorporating a single complementary nucleotide into its spacer. In this context, the PAM might better be referred to as a protospacer-associated motif to reflect its origin and function (10).

Base pairing between spacer and protospacer sequences is crucial for interference (48). As the +1 PAM nucleotide is derived from the protospacer, and the complementary pyrimidine is captured into the CRISPR array, base pairing between the PAM and the 5’ nucleotide of the crRNA spacer is inevitable, but such pairing can be artificially manipulated to investigate its importance. In *E. coli*, base pairing between the 5’ crRNA guanine and +1 PAM cytosine in the phage M13 g8 spacer-protospacer pairing is dispensable for interference (49). In contrast, in *V. cholerae* we show positive, negative, and neutral consequences in interference, depending upon the spacer, when this base pairing is abolished. The effect of base pairing on PAM recognition and interference warrants further investigation, as a greater understanding of this phenomenon may permit better design of synthetic spacers.

In conclusion, we found evidence that the CRISPR/Cas system originally observed in an extinct strain of *V. cholerae* continues to be present in other strains today. By mining deposited sequences, we identified a conserved 5’-pyrimidine in naturally occurring spacers and found that their cognate protospacers contain a complementary purine. This purine was found to be essential for interference when crRNA and Cas proteins are not over-expressed, thus we have redefined the PAM of the system to 5’-RTT-3’. Finally, we demonstrated interference is affected by base pairing between the protospacer-derived 5’ pyrimidine and the complementary +1 PAM purine. Our findings further the understanding of the molecular mechanisms of Type I-E CRISPR/Cas systems. The resulting enhancement in our knowledge of the requirements for interference should allow for more intelligent design of targeting spacers and permit efficient utilization of CRISPR in genomic manipulation of *V. cholerae* and its phages.

## Materials and Methods

### Bacterial strains and culture conditions

Bacterial strains and plasmids used in this study are listed in Table 1. Bacteria were cultivated at 37°C in Luria Broth (LB) agar or in LB. Medium was supplemented with ampicillin (Amp; 50 μg/mL), kanamycin (Kan, 50 μg/mL), spectinomycin (Spec, 100 μg/mL), and/or streptomycin (Sm, 100 μg/mL) when appropriate. For induction, LB agar plates were supplemented with 100 μM isopropyl-β-d-thiogalactopyranoside (IPTG).

### Bioinformatic analysis of CRISPR repeats in deposited sequences of *V. cholerae*

Additional strains containing the 28 bp *V. cholerae* O395 CRISPR repeat (5’-GTCTTCCCCACGCAGGTGGGGGTGTTTC-3’) were identified using BLAST to search the NCBI whole-genome shotgun contigs database, restricting results to the family *Vibrionaceae*. This method would identify O395 CRISPR repeats when present in other *Vibrio* species, however, we found that only *V. cholerae* strains contained the perfect repeat. Extraction of spacer content and protospacer mining was automated using a custom Python script (https://github.com/camillilab/spacer_miner). Briefly, identified contigs containing repeat sequences were retrieved from NCBI. Spacers were extracted if its sequence was no longer than 50 nucleotides, were flanked by perfect repeat sequences, and did not contain ambiguous nucleotides. Each unique spacer was then compared with BLAST to the NCBI non-redundant nucleotide database to identify putative protospacers. Putative protospacers were considered if there was at least 93% identity to the corresponding spacer over 96% of the spacer sequence, which permits up to two mismatches and one missing or additional base in the protospacer hit. Sequence logos were generated using unique spacers and nucleotides 3’ of identified protospacers using WebLogo (50).

### Conjugation assays

Donor plasmid pDL1301, an RP4-conjugatable plasmid that confers resistance to spectinomycin, and its variants were transferred using *E. coli* SM10λpir. Donor and recipient AC6625 were grown to an OD_600_ of 1.0. Then, 500 μL of donor and recipient were pelleted, washed once in phosphate-buffered saline (PBS), and resuspended in 50 μL PBS. A 1:1 mixture was applied to a sterile filter (0.22 μm pore, Millipore) on an LB plate, and incubated at 37°C for 2 hours. Bacteria were recovered from the filter by vortexing in 500 μL PBS. Serial dilutions were plated onto medium selective for recipient *V. cholerae* and exconjugates, and separately in the presence of IPTG. The conjugation efficiency was calculated as the number of exconjugates divided by the total number of viable *V. cholerae* recipients.

### Construction of targeting plasmids

CRISPR targeting plasmids were generated using the previously constructed pCRISPR backbone (15). New spacers were constructed using annealed and phosphorylated oligonucleotides that included the targeting region flanked by appropriate 5’ and 3’ overhangs to facilitate ligation into pCRISPR. The resulting double-stranded DNAs were cloned into pCRISPR by Golden Gate cloning using BsaI-HF (New England Biolabs). Ligation products were purified and used to transform electrocompetent *E. coli* SM10λpir. DNA from individual clones was isolated and the targeting sequence was verified by Sanger sequencing. These plasmids were then mated into the *V. cholerae* targeting strain AC6625.

### Generation of transversion mutant donor and targeting plasmids

The target *aad9* gene in donor plasmid pDL1301 was modified in eight independent sites by site-directed PCR mutagenesis to yield eight new donor plasmids possessing one silent mutation each (*aad9\**). The location of each of these mutations is listed in Table 3. Eight new spacers that target these mutated sites were introduced into targeting plasmid pCRISPR as previously described to yield eight new targeting plasmids (crRNA*).

## Acknowledgements

This work was supported by NIH grants AI055058 (AC), AI147658 (AC), GM139722 (JB) and GM008448 (JB).

## Supplementary Tables

**Supplementary Table 1:**
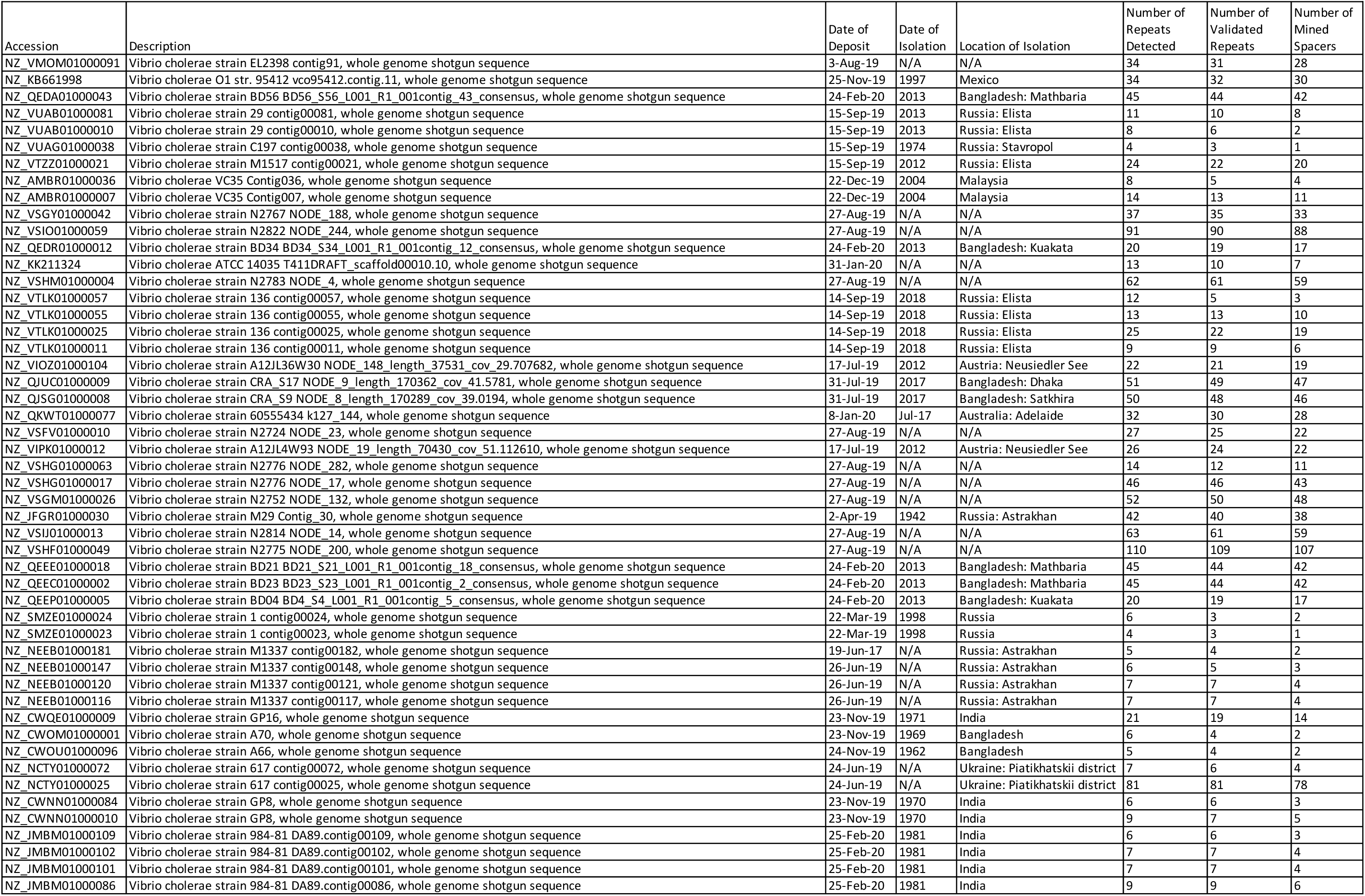
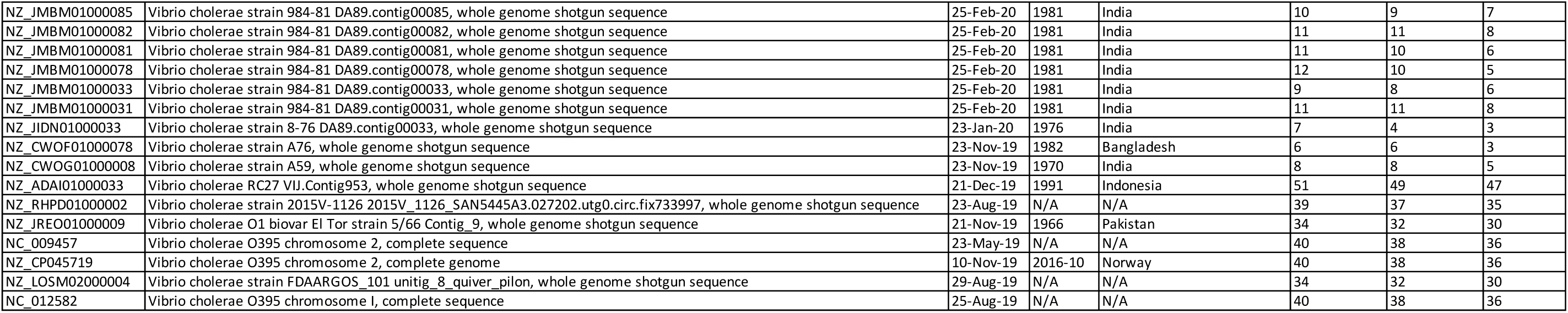
Overview of deposited *V. cholerae* sequences harboring the Type I-E CRISPR/Cas 28 bp repeat sequence.

**Supplementary Table 2:**
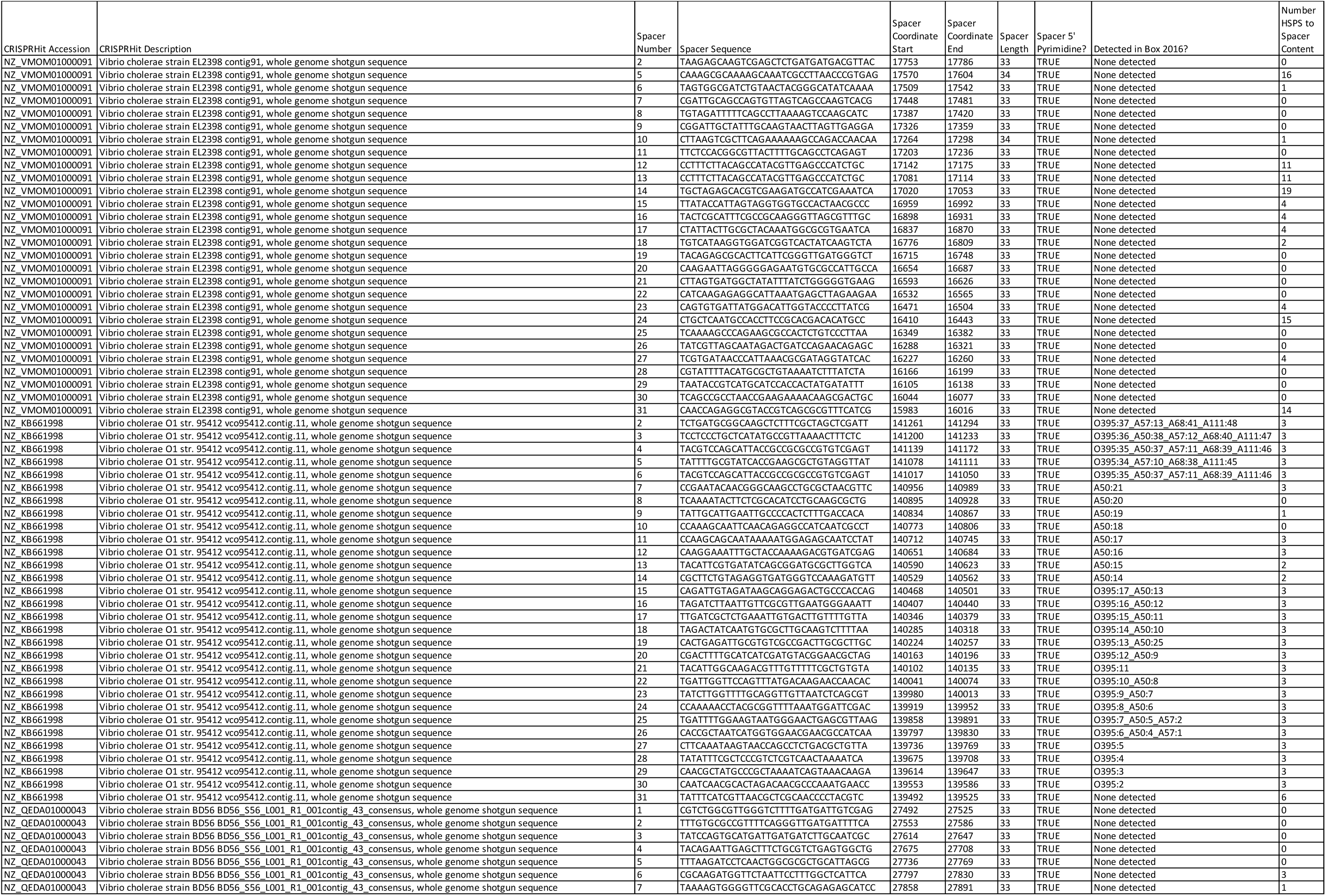
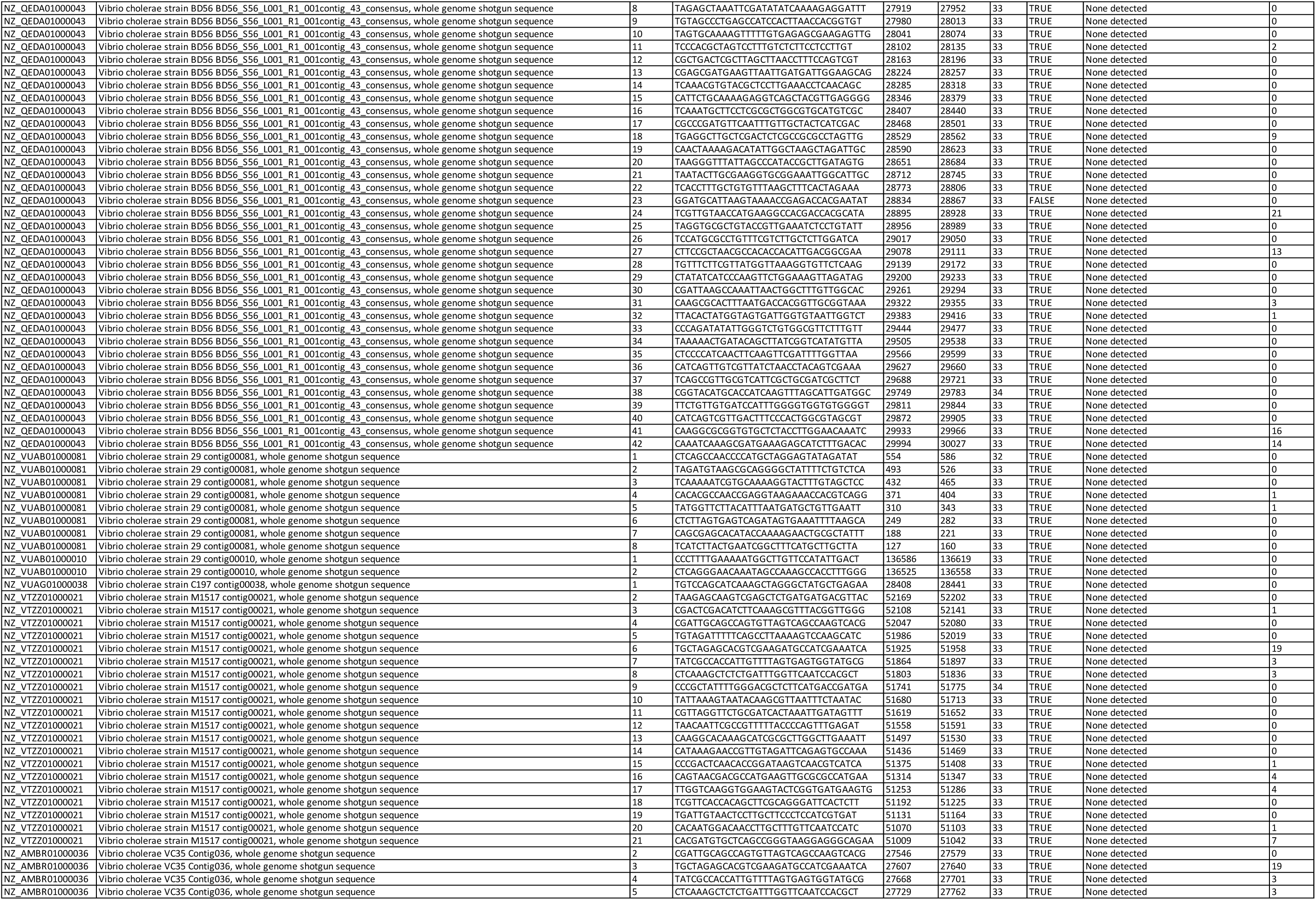
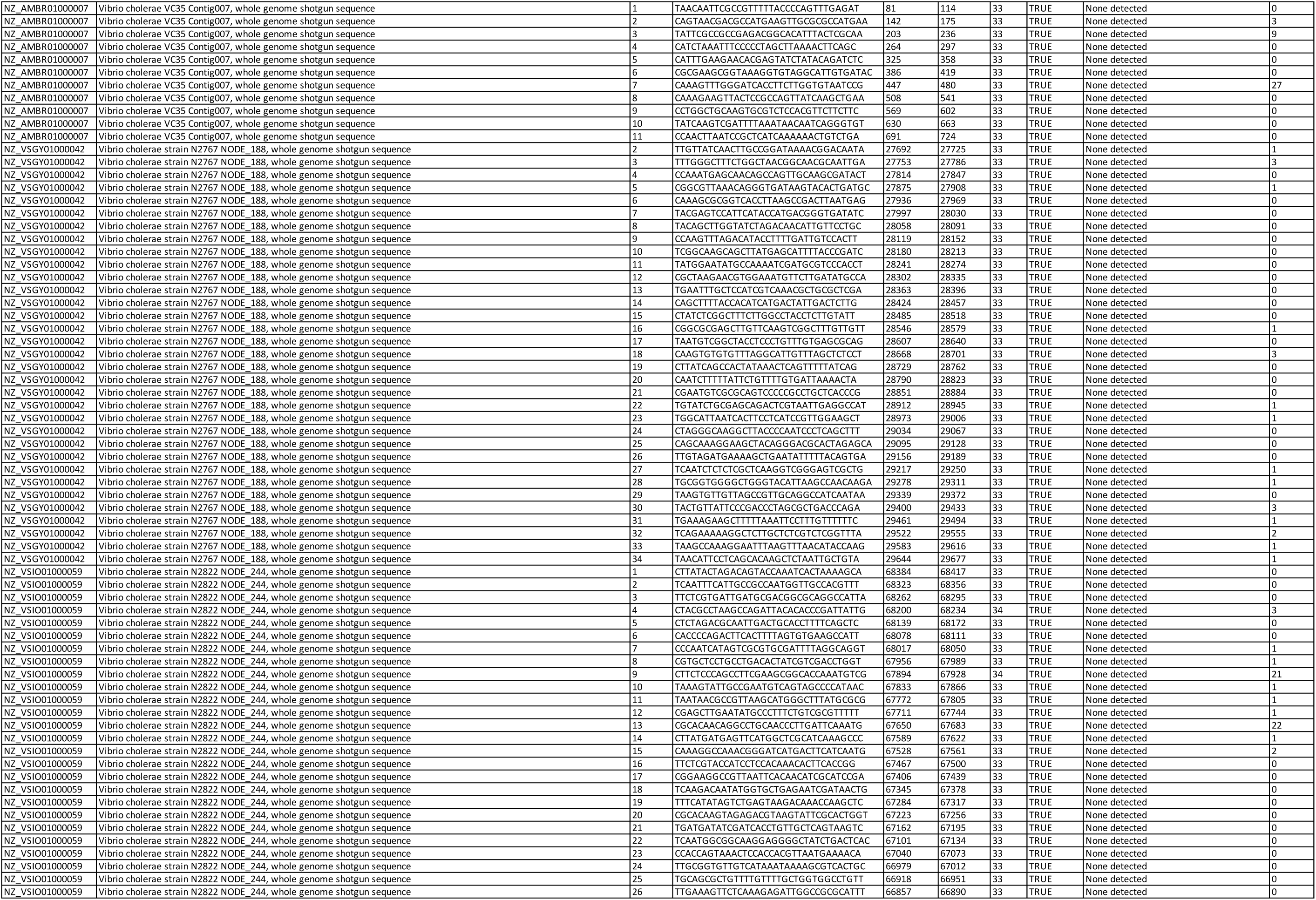
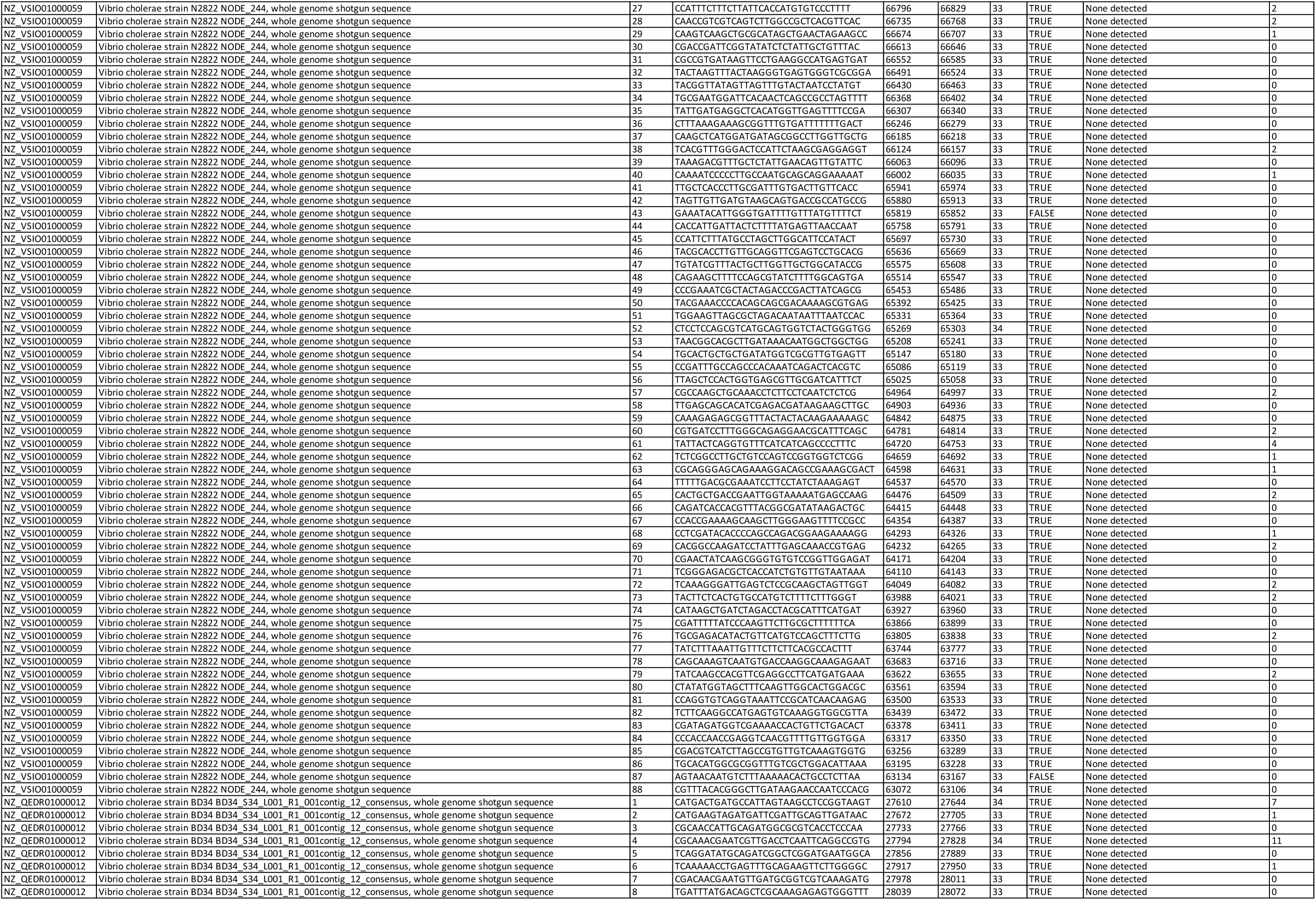
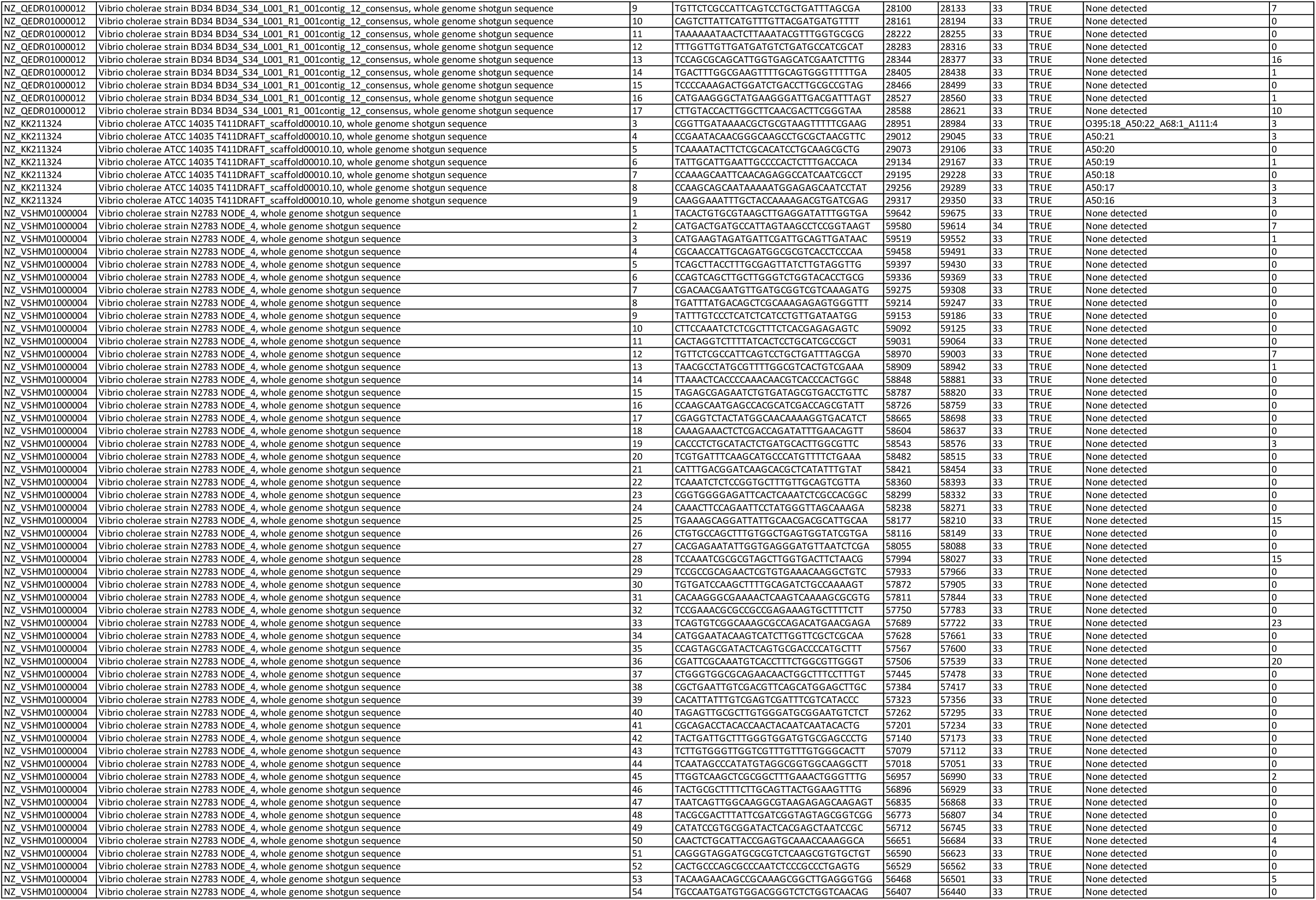
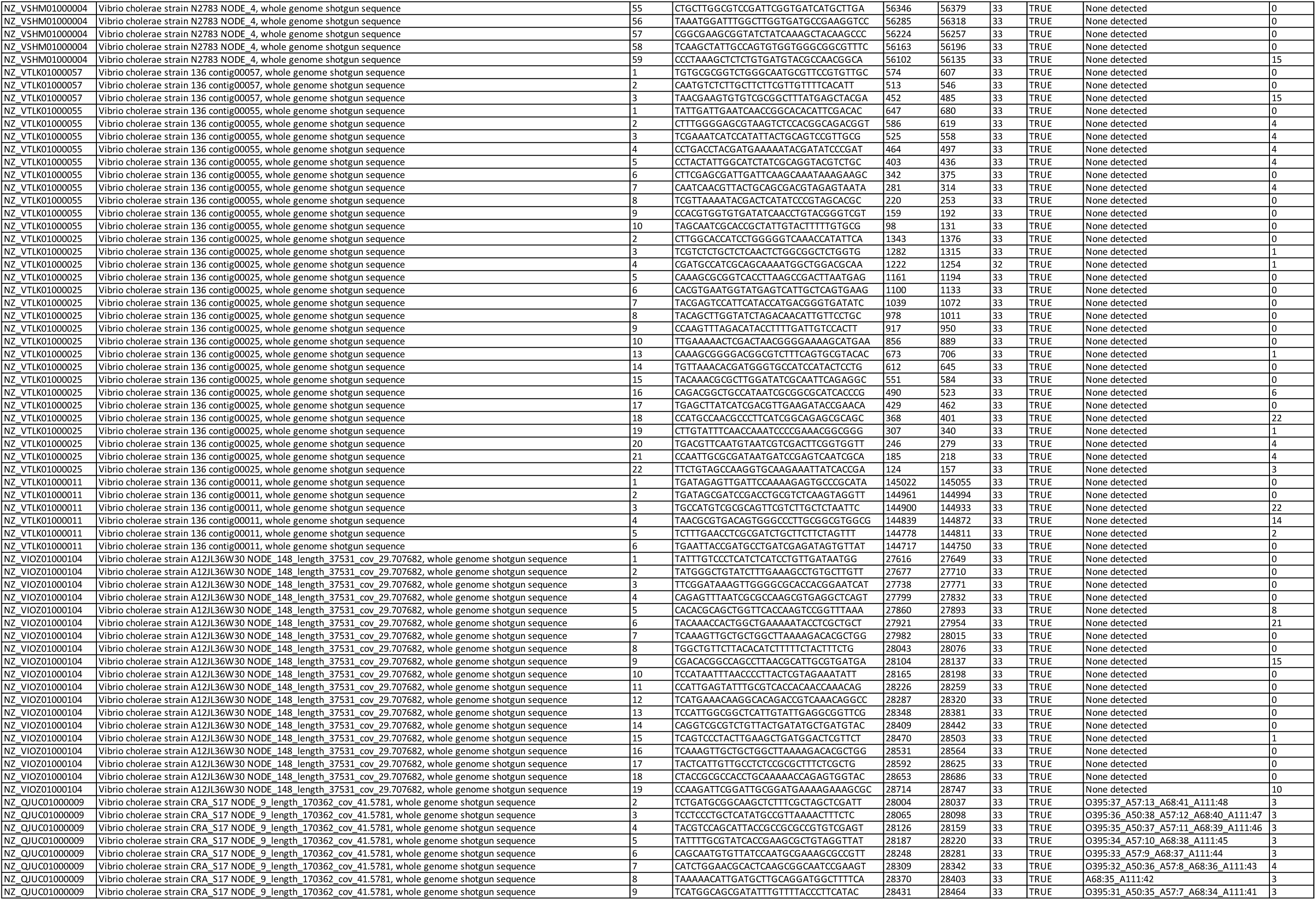
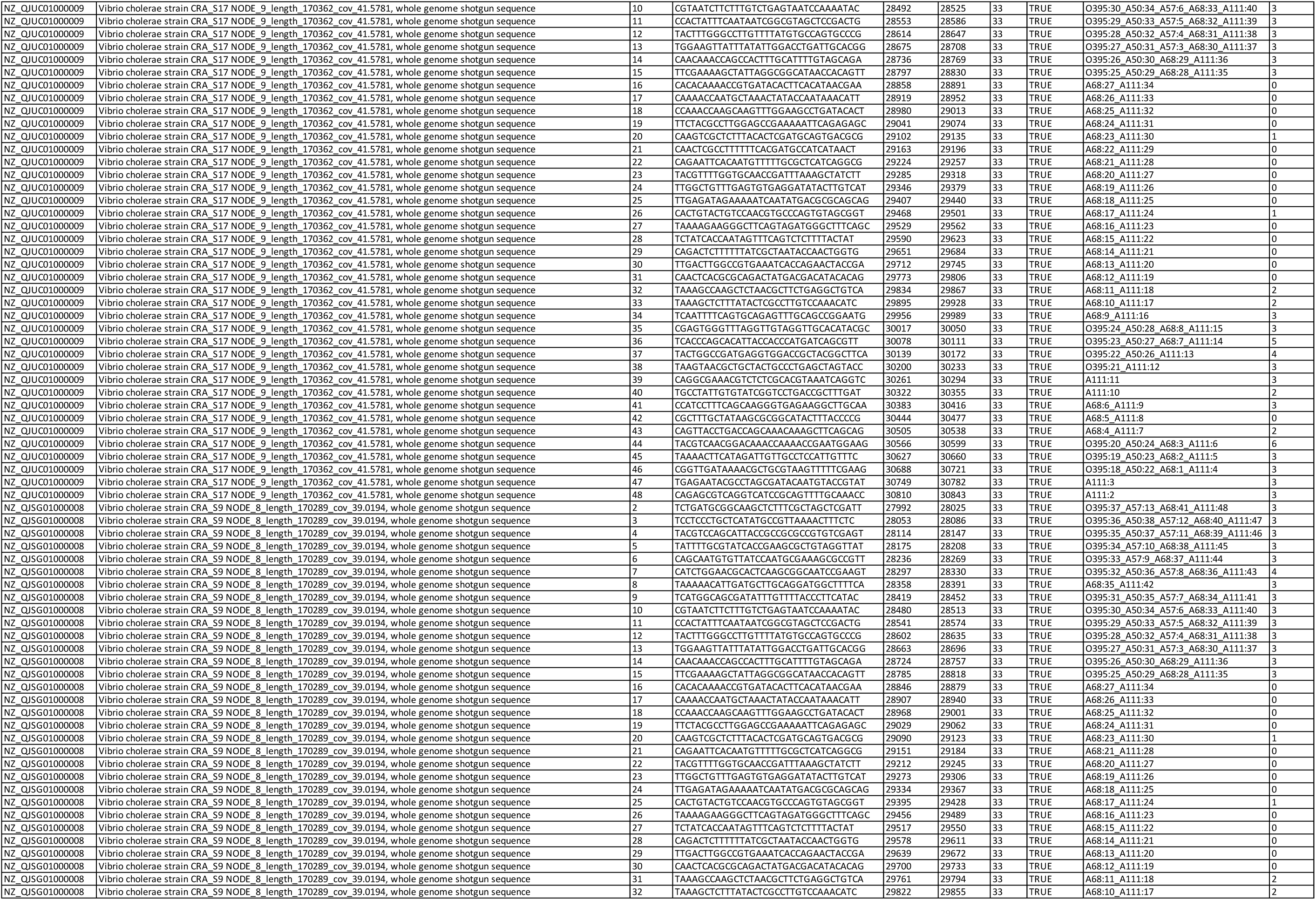
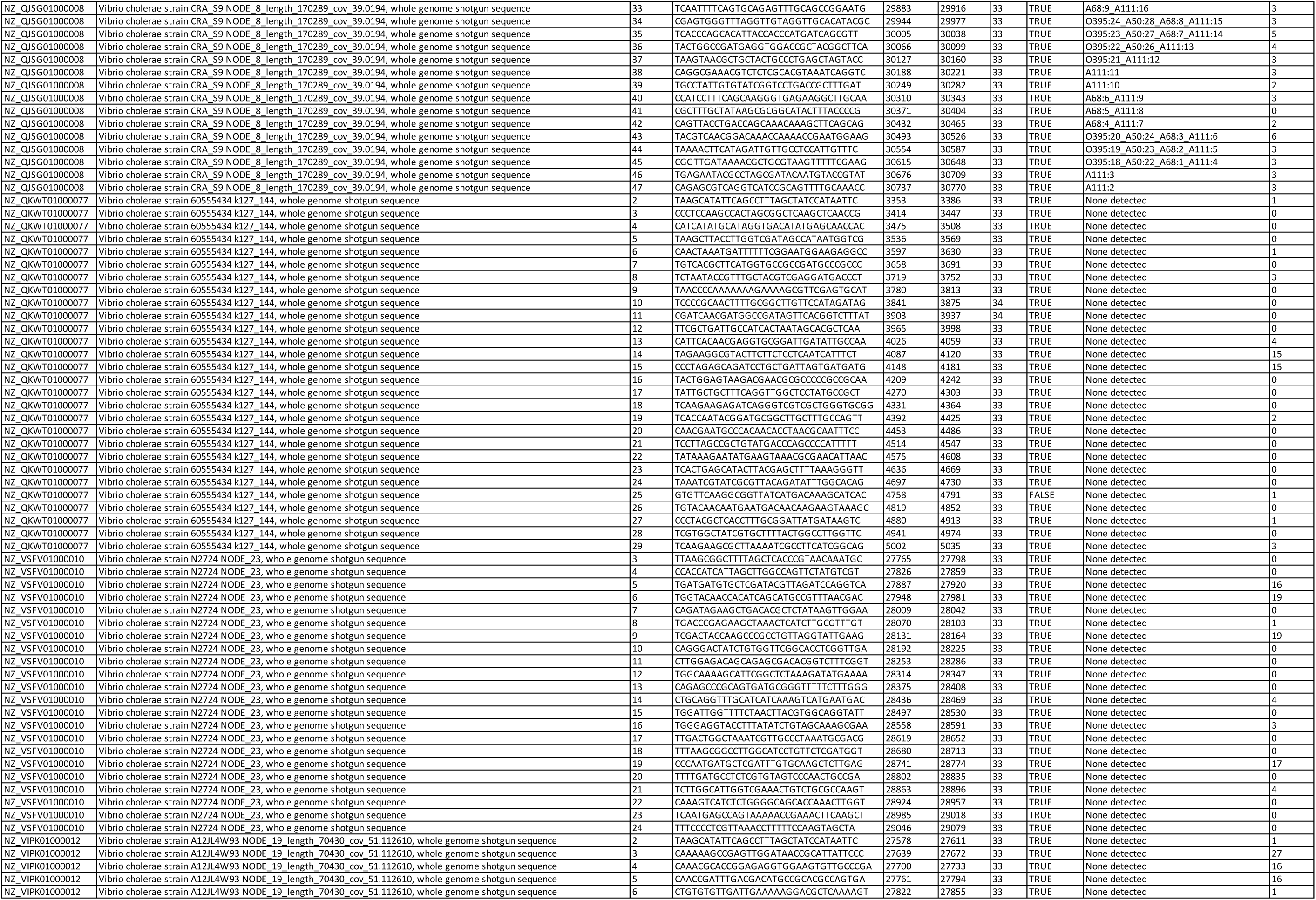
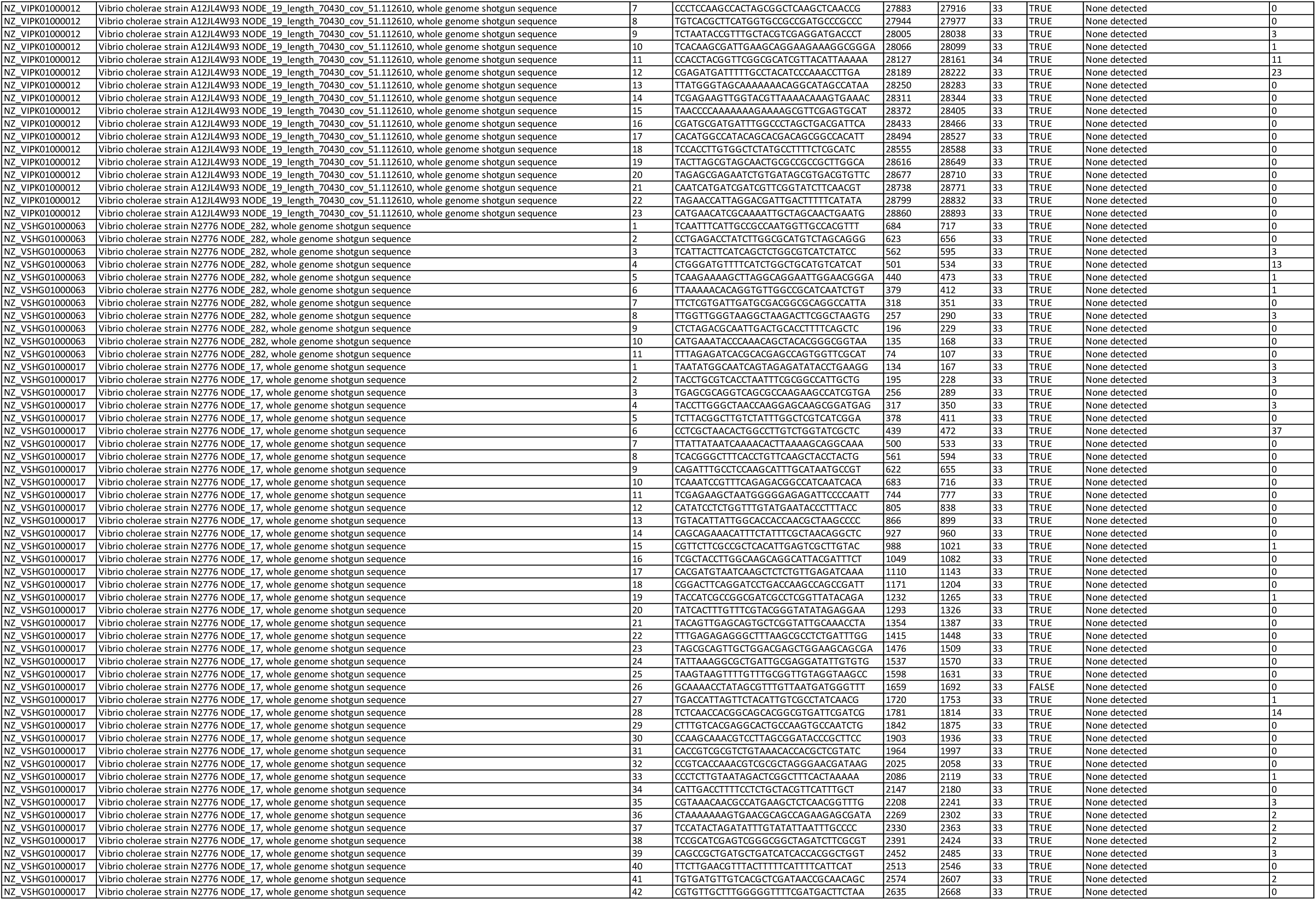
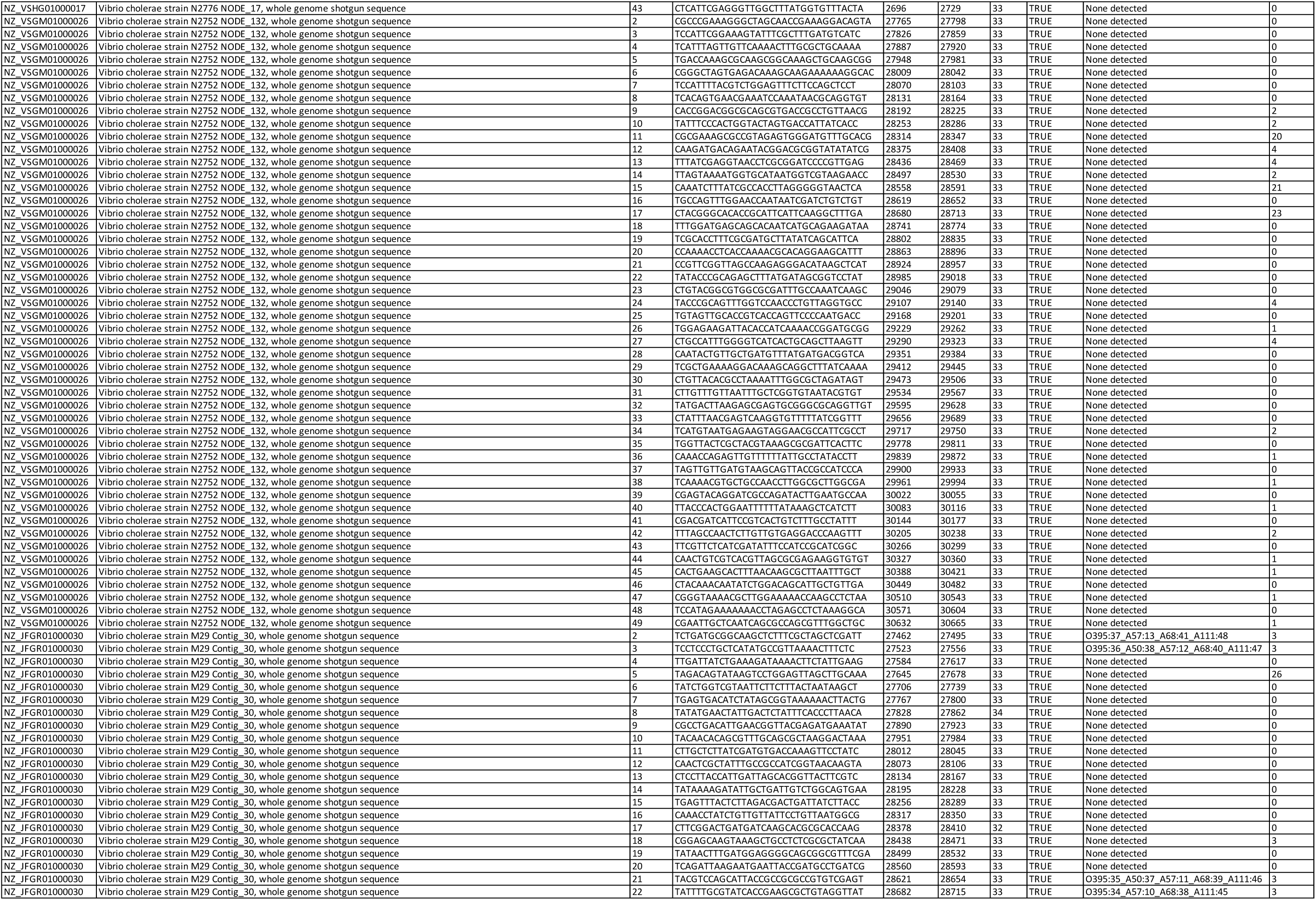
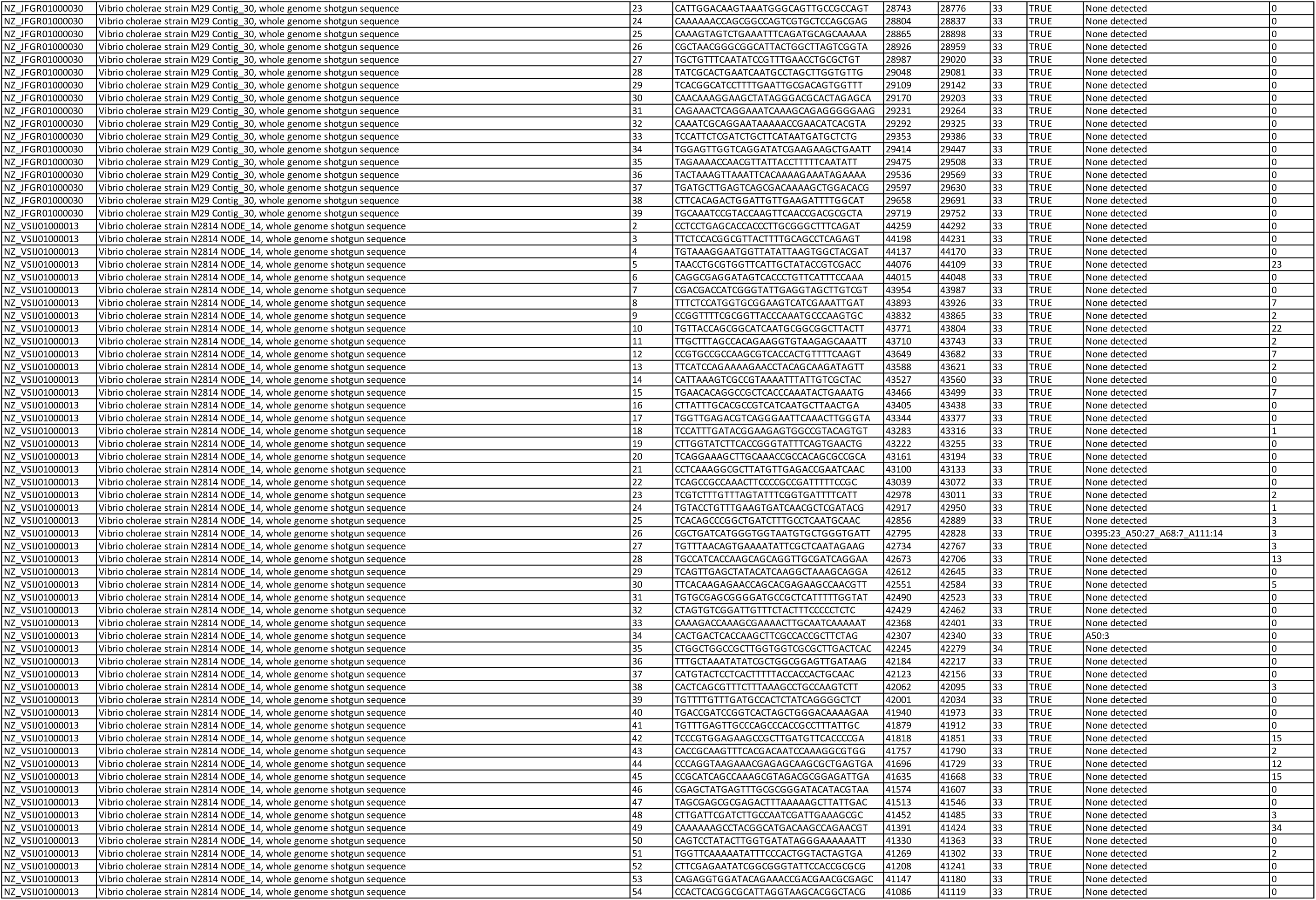
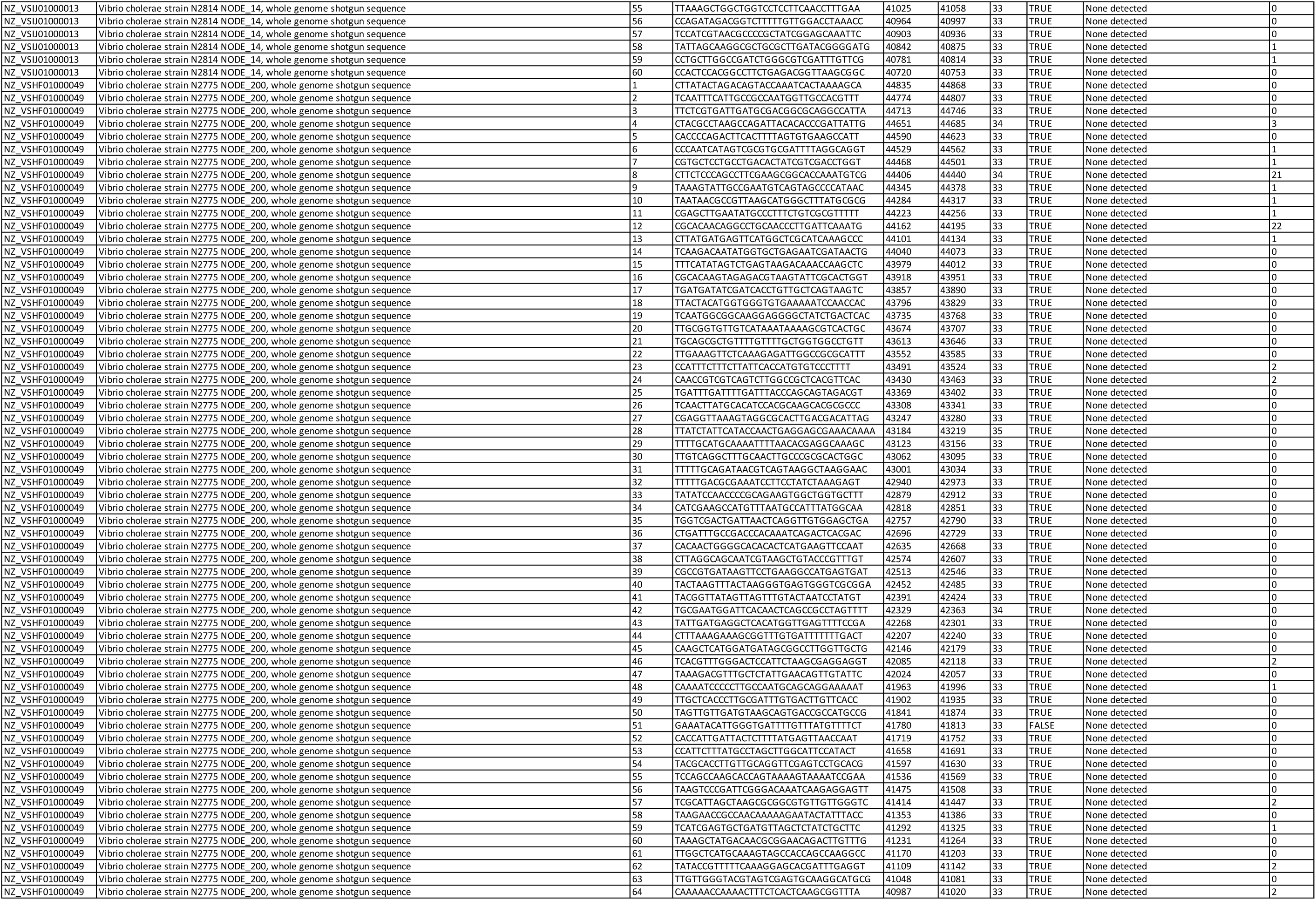
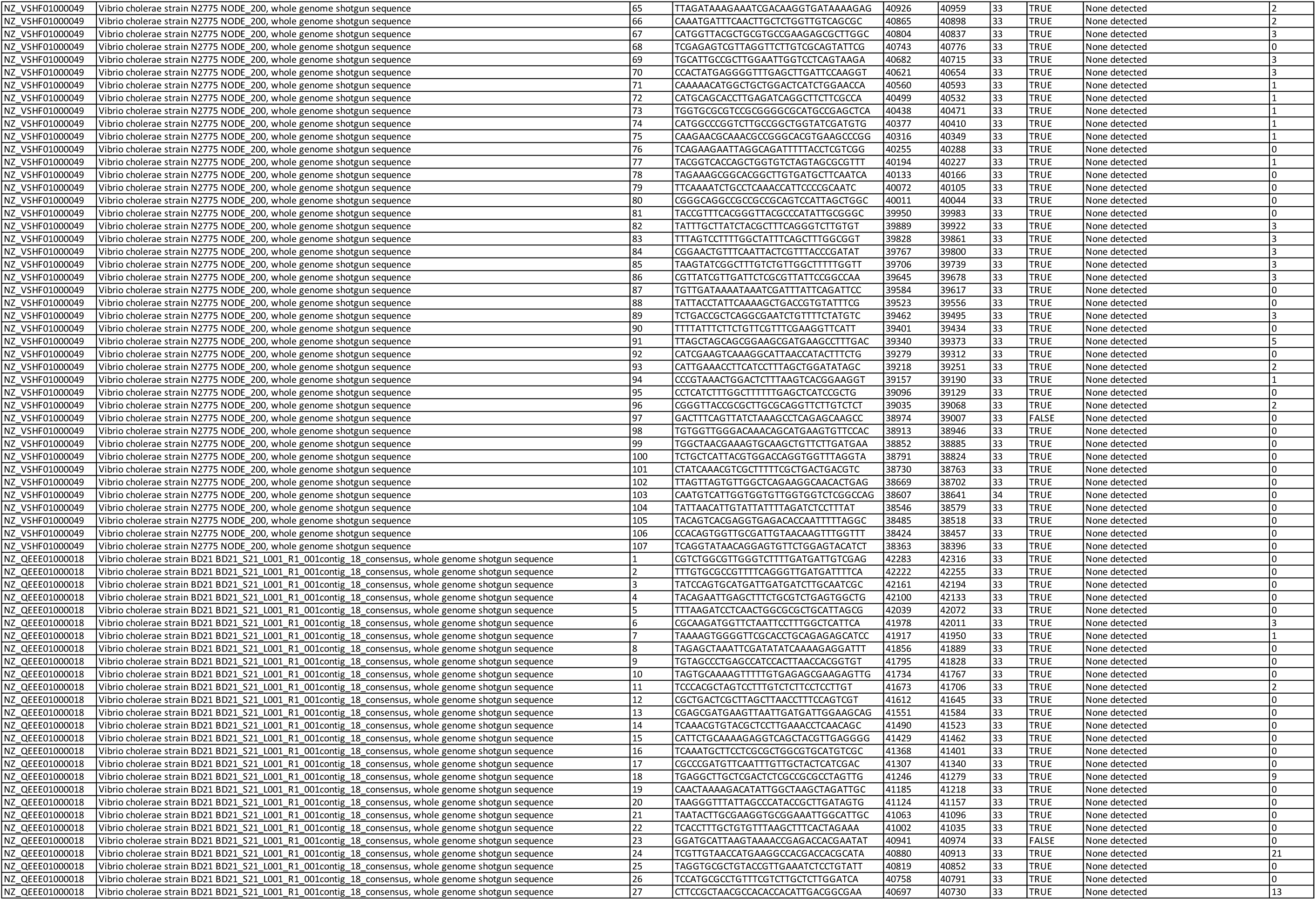
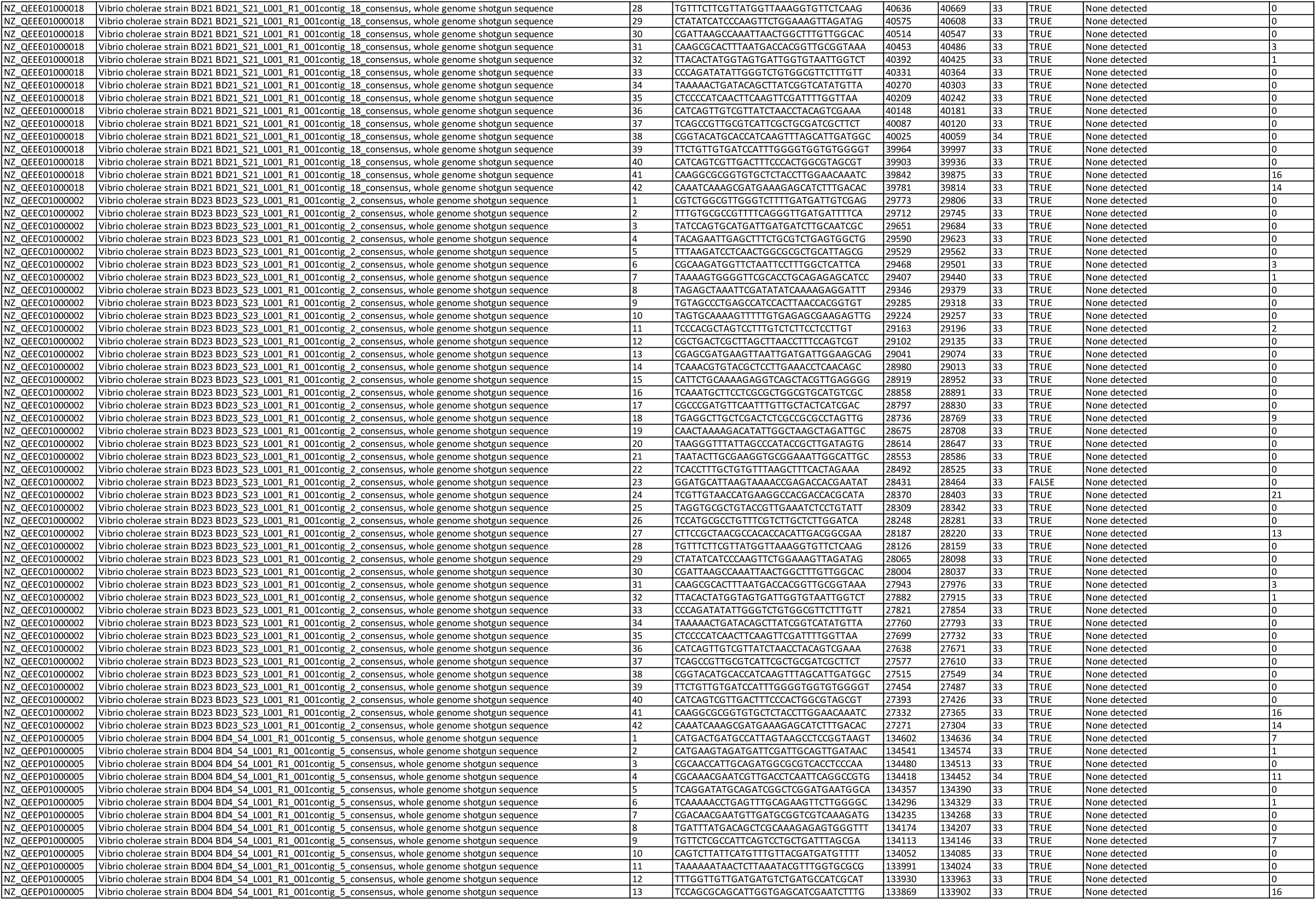
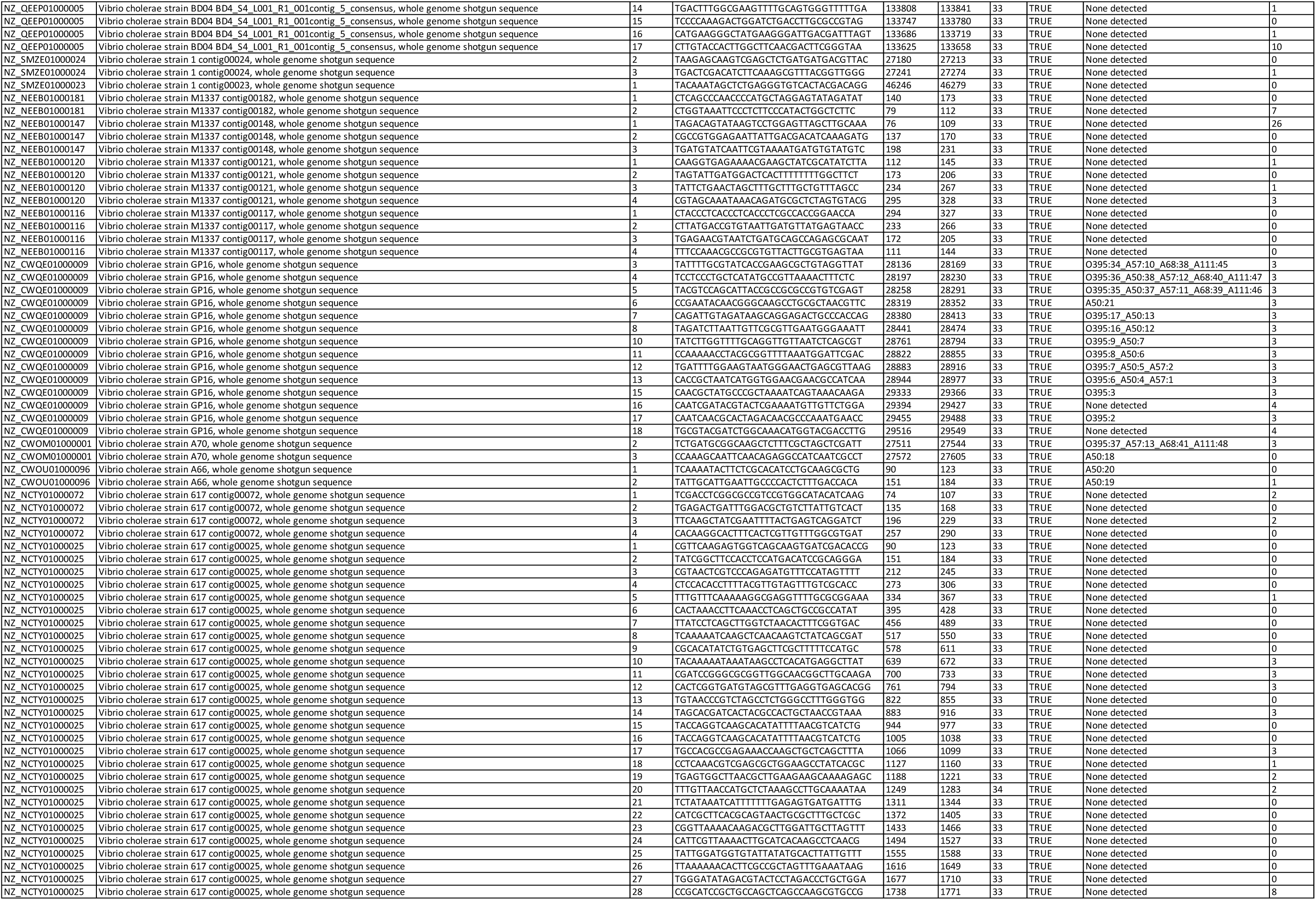
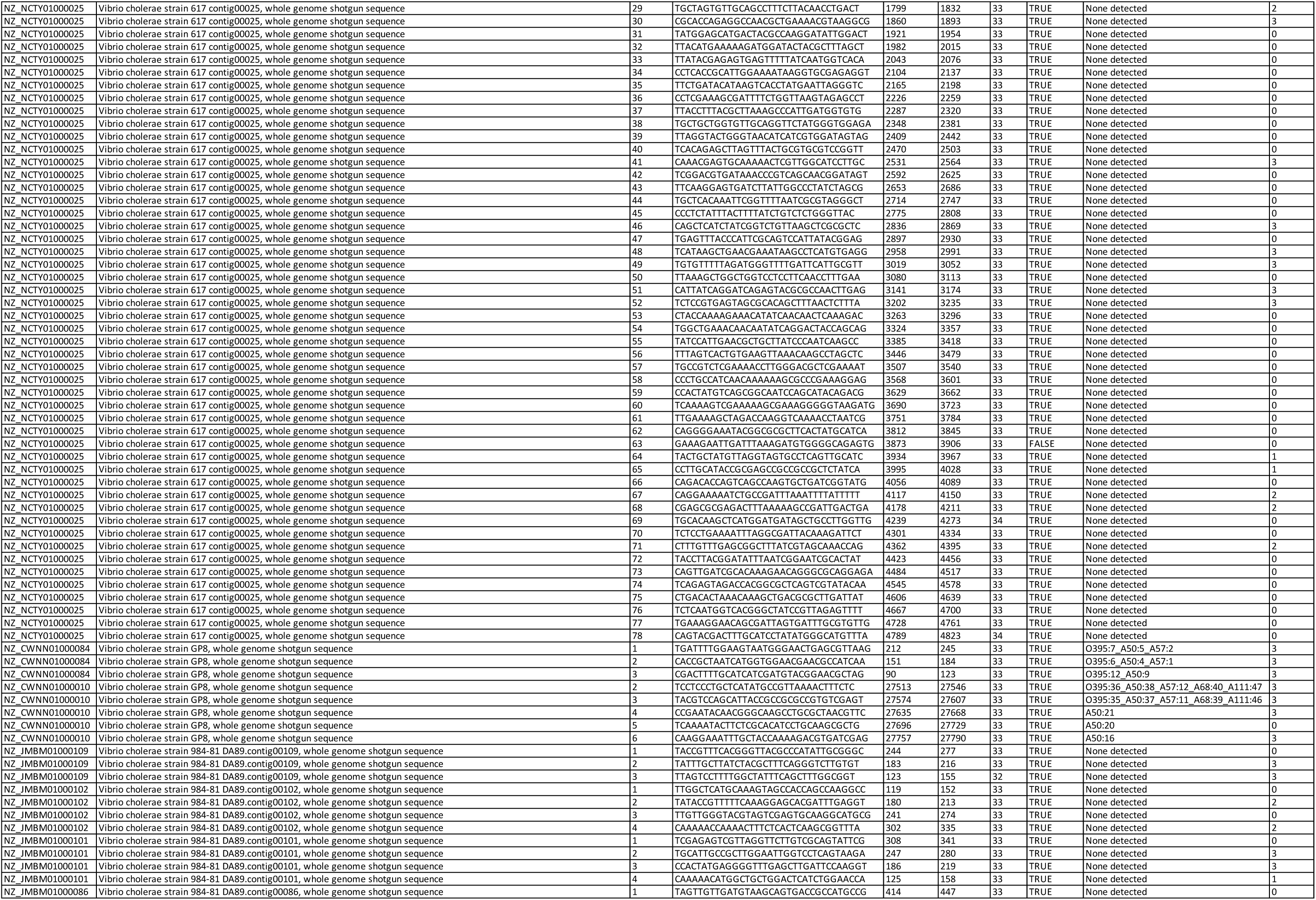
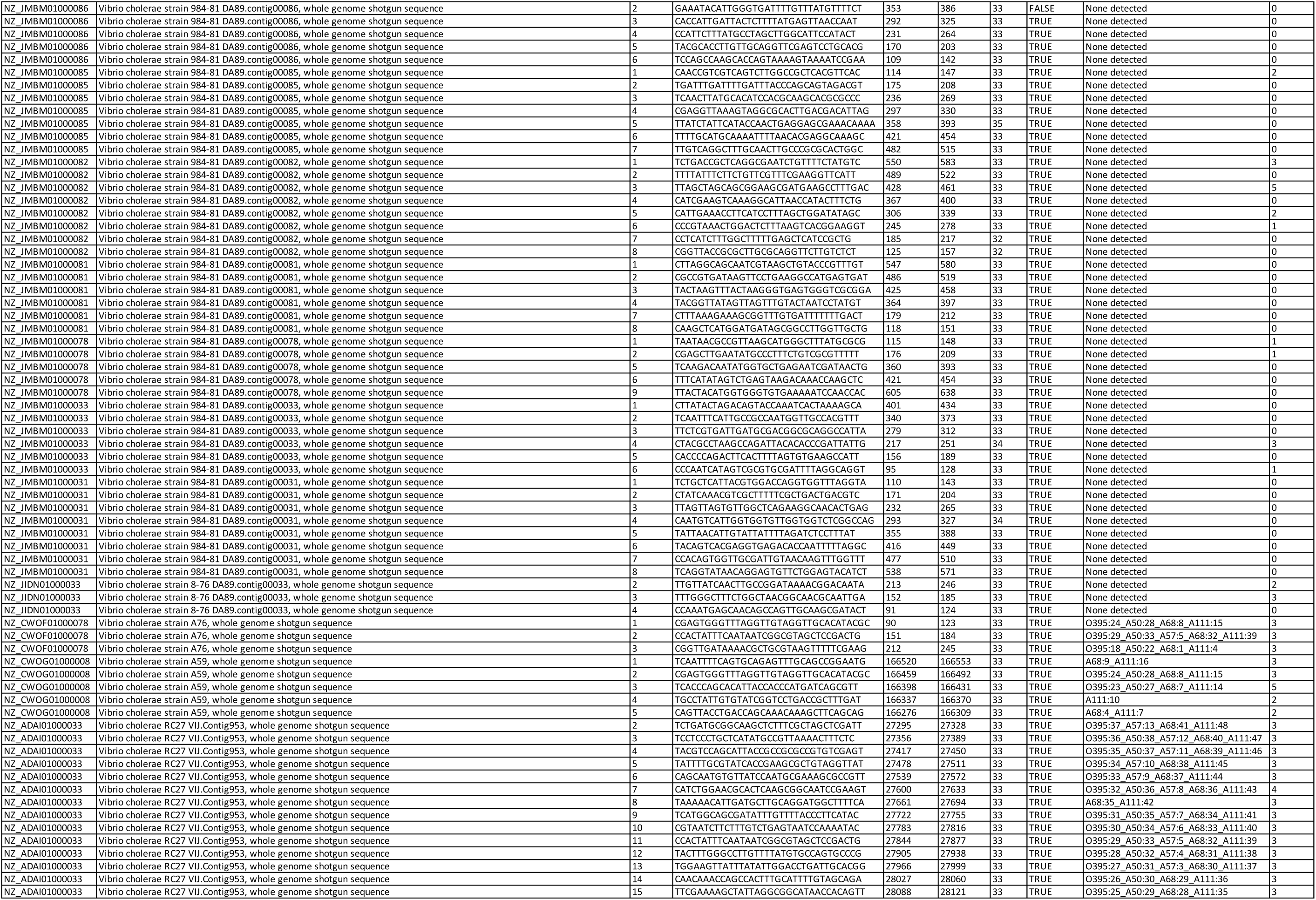
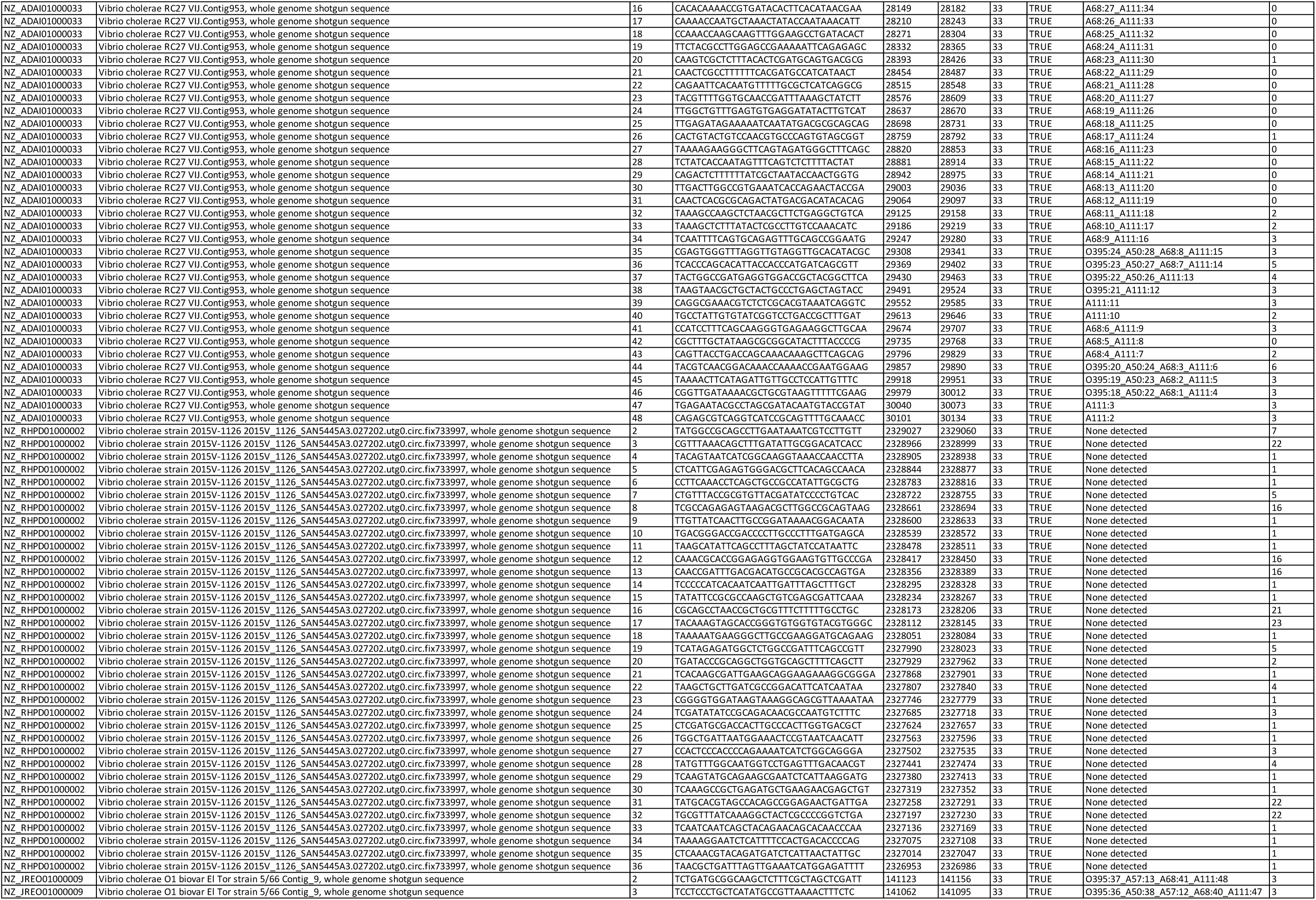
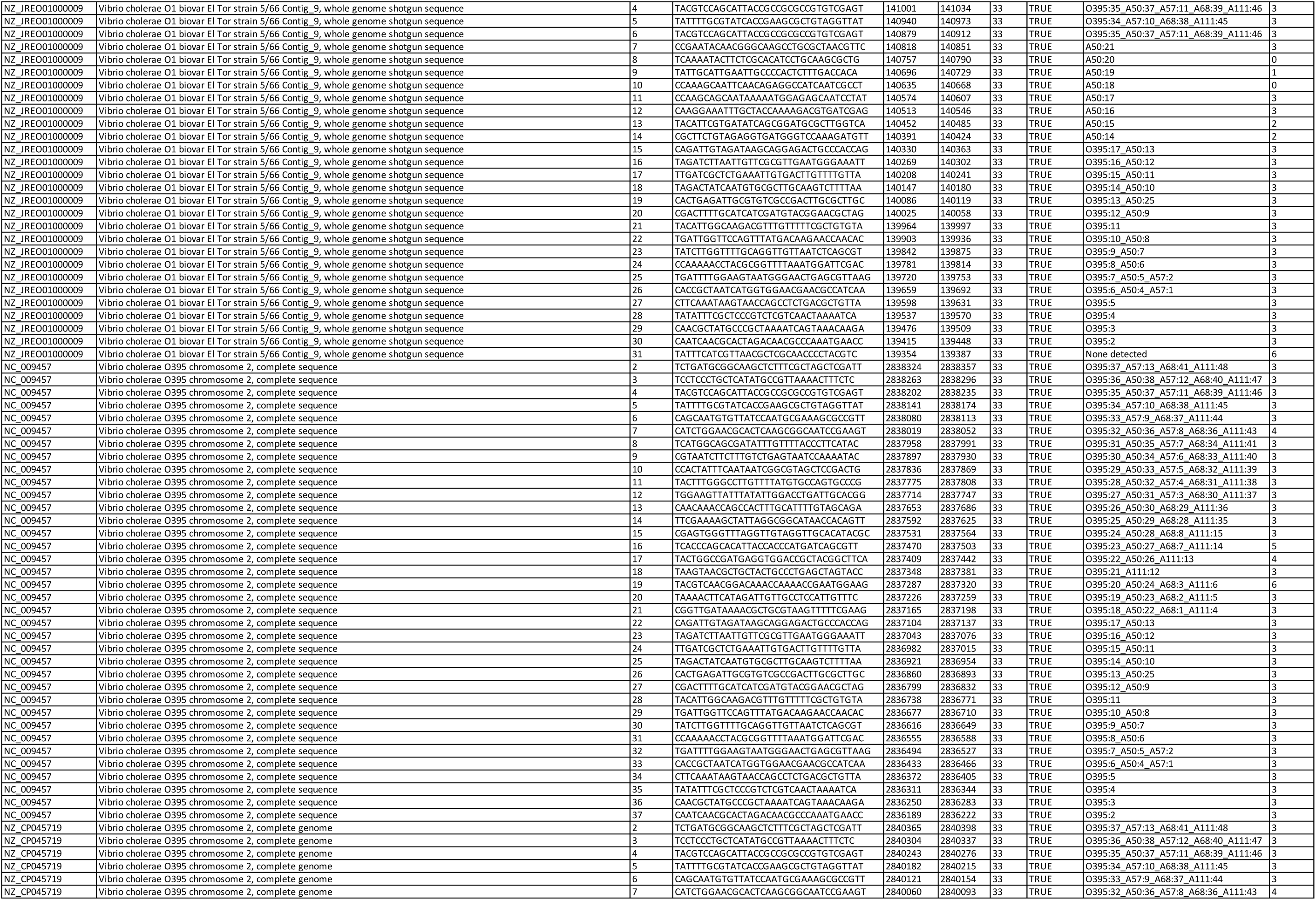
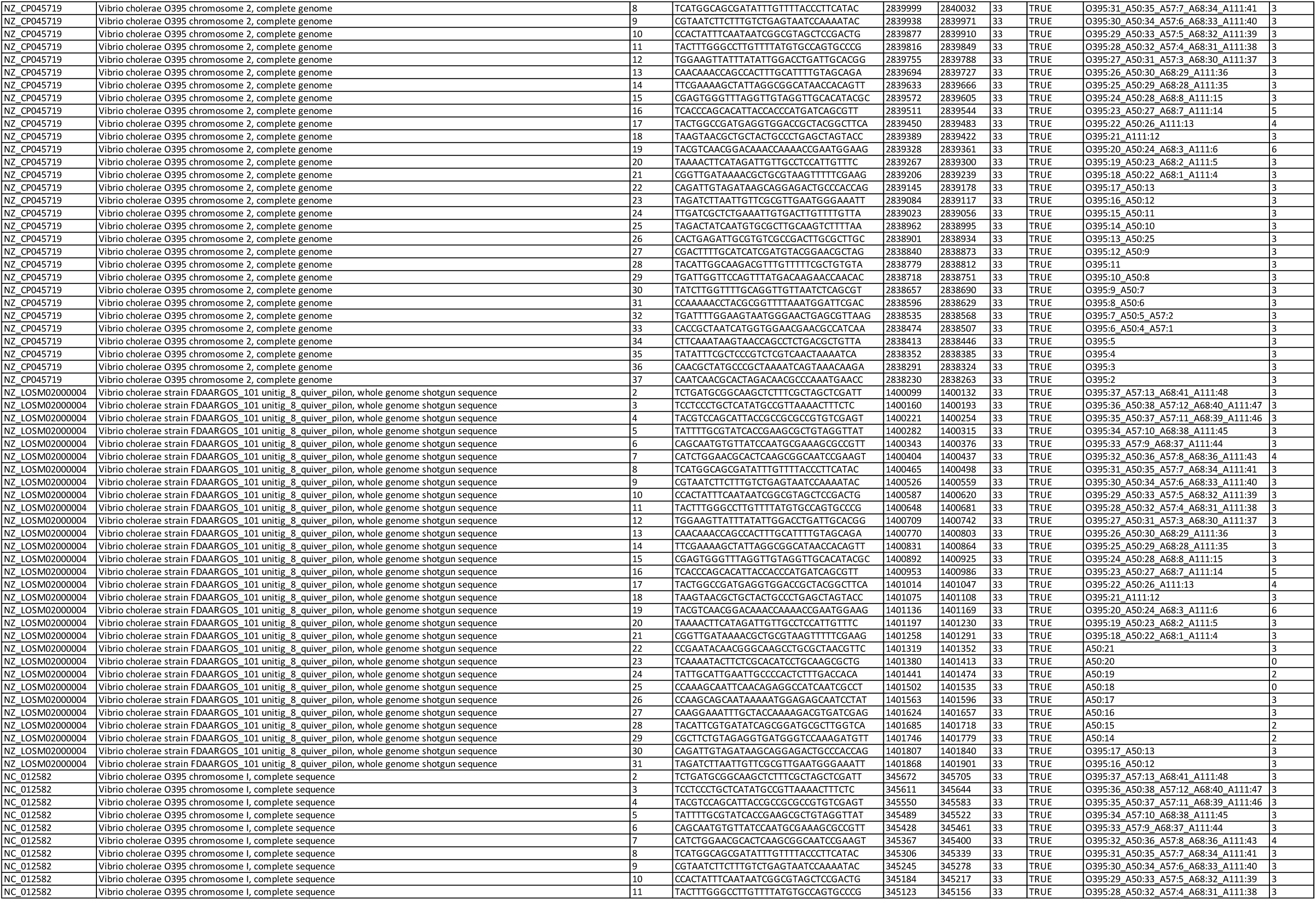
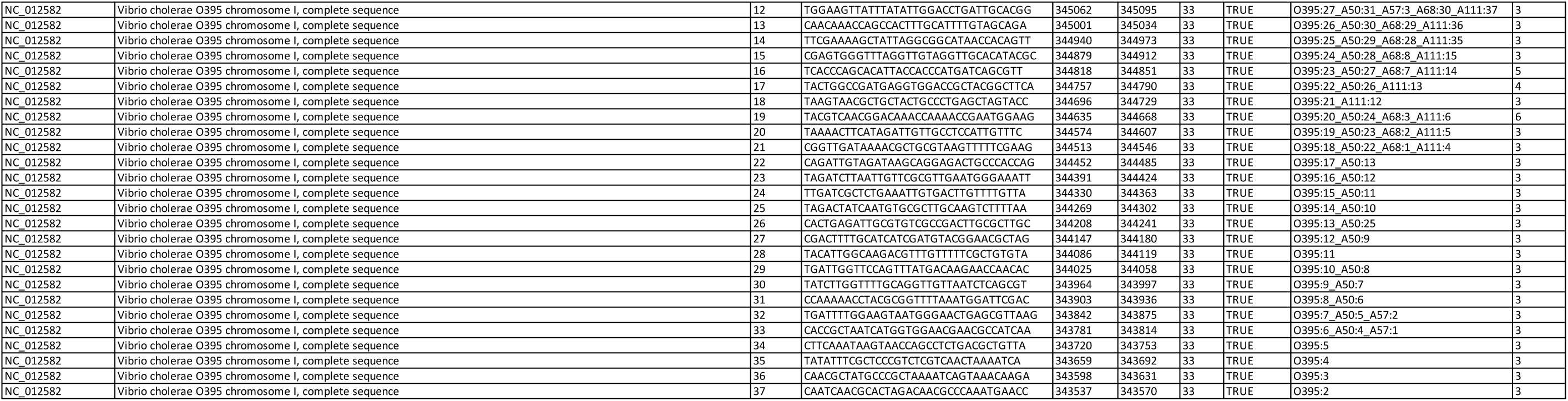
Overview of mined spacers from identified *V. cholerae* sequences harboring the Type I-E CRISPR/Cas system.

**Supplementary Table 3:**
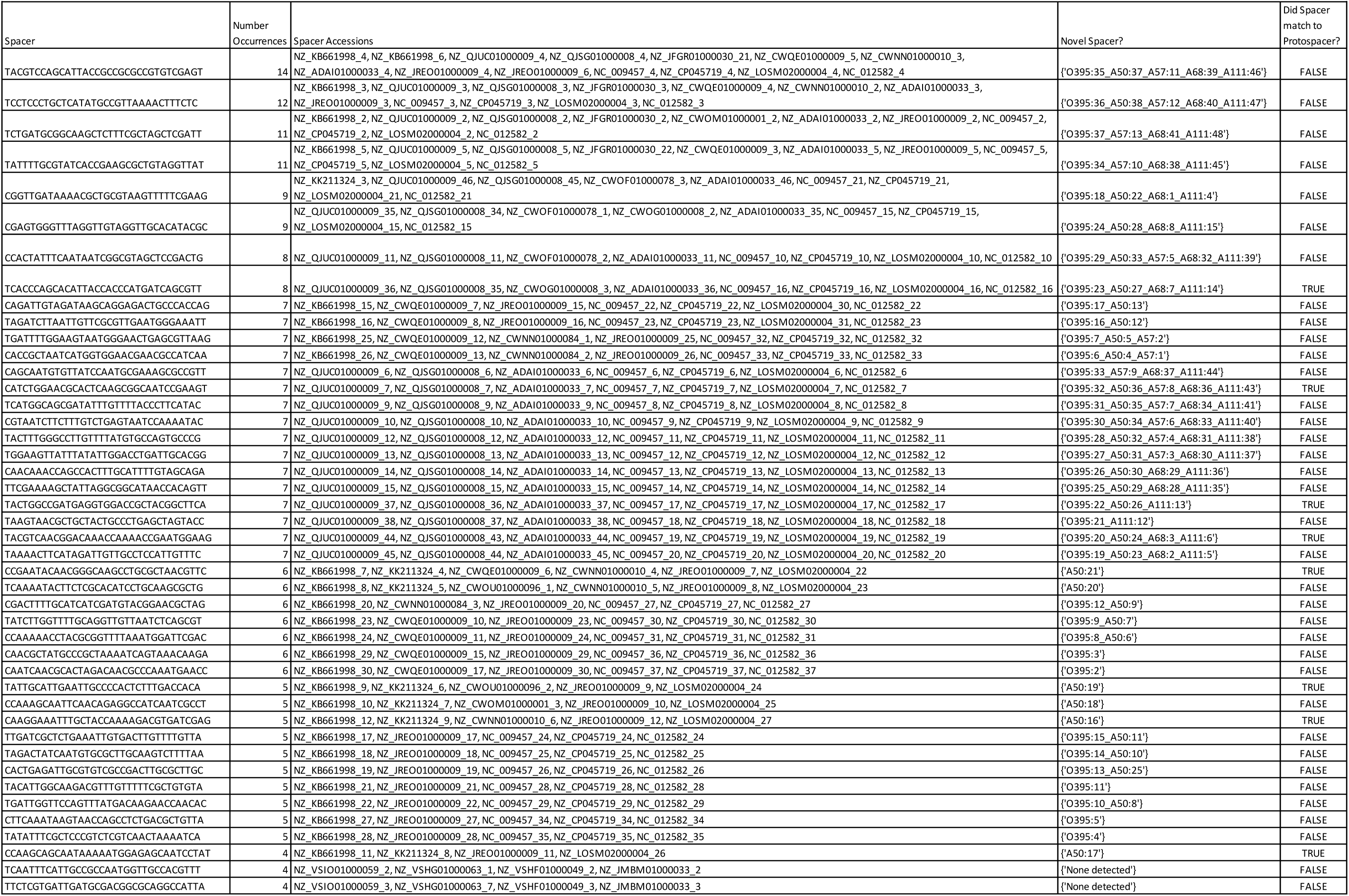
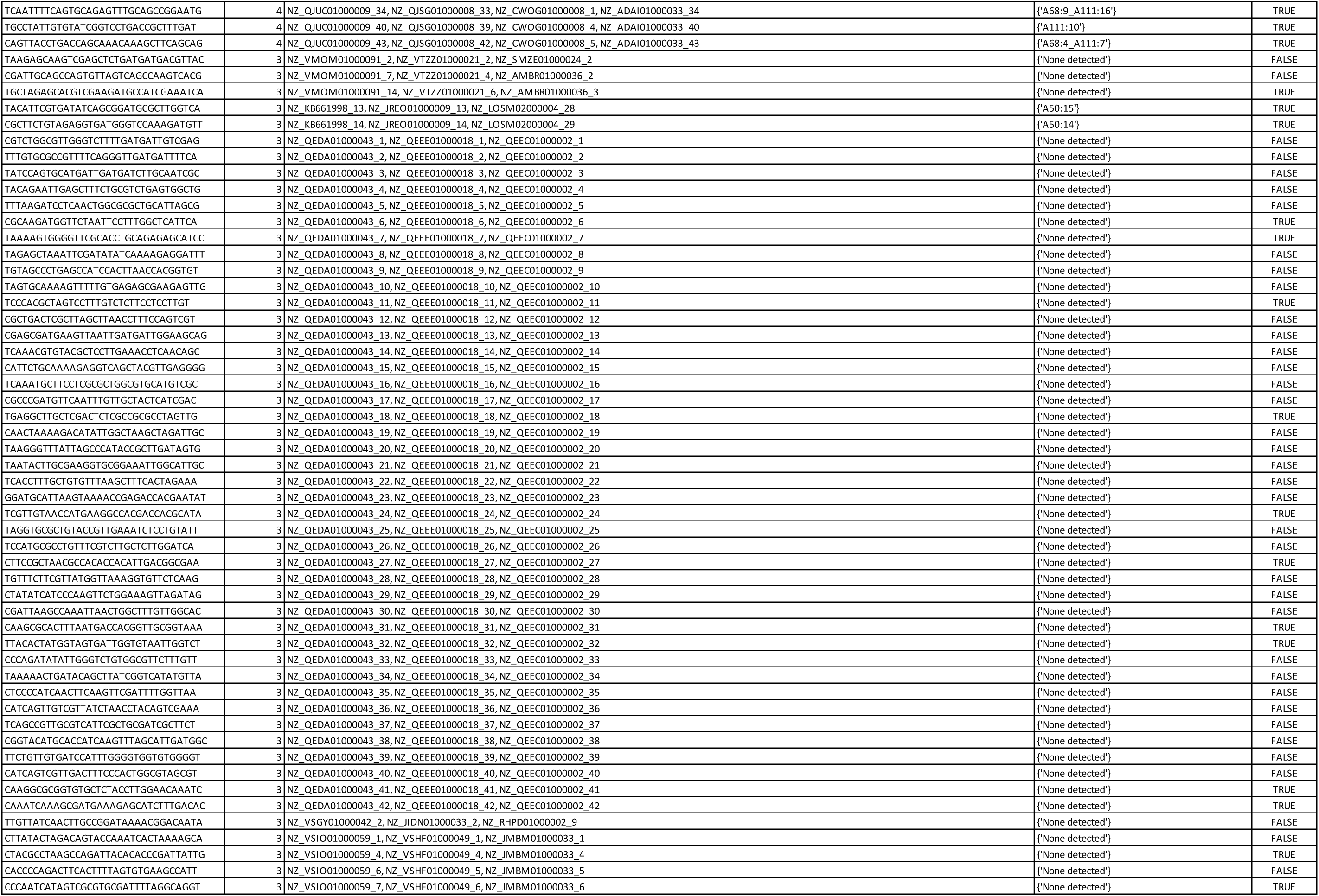
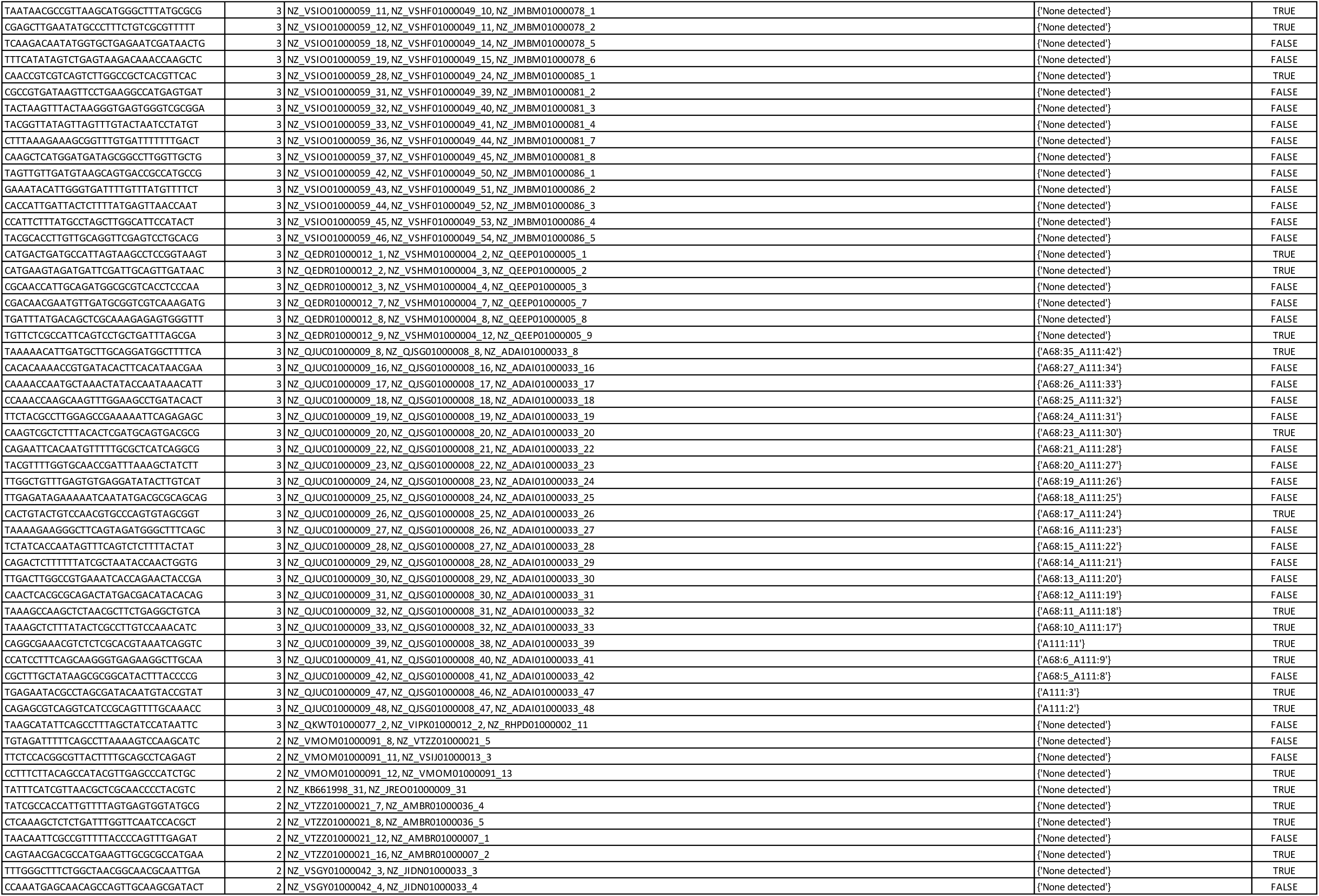
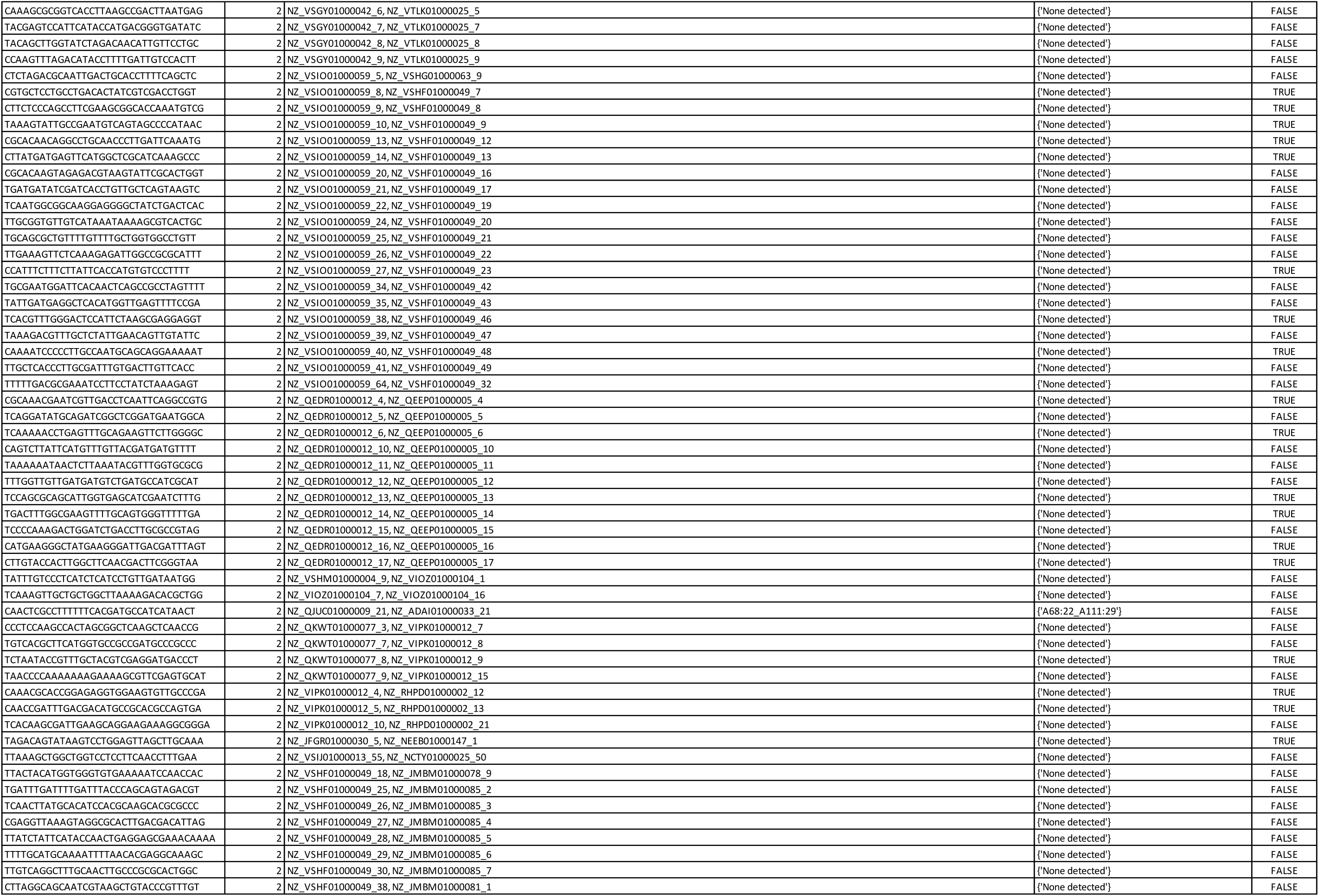
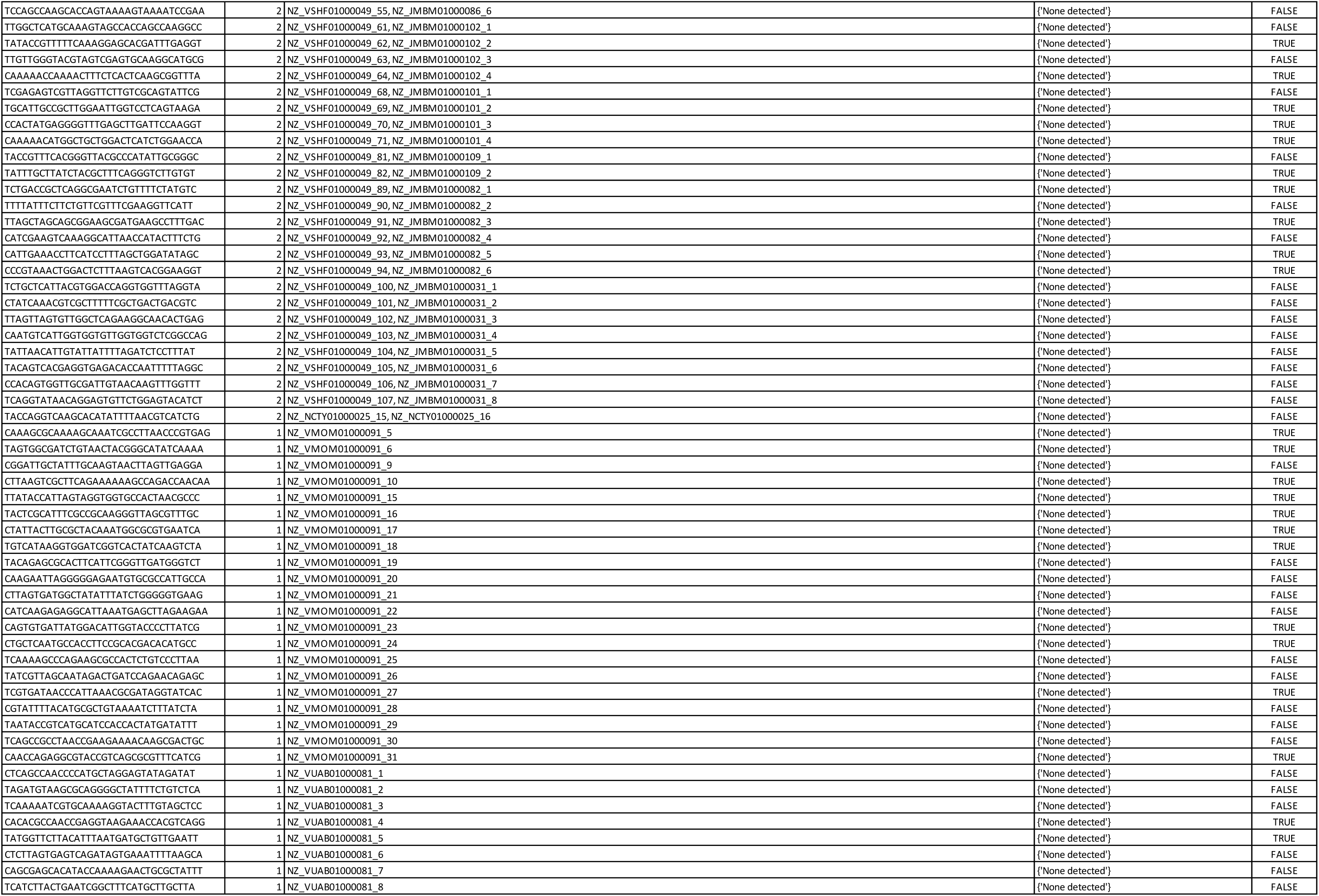
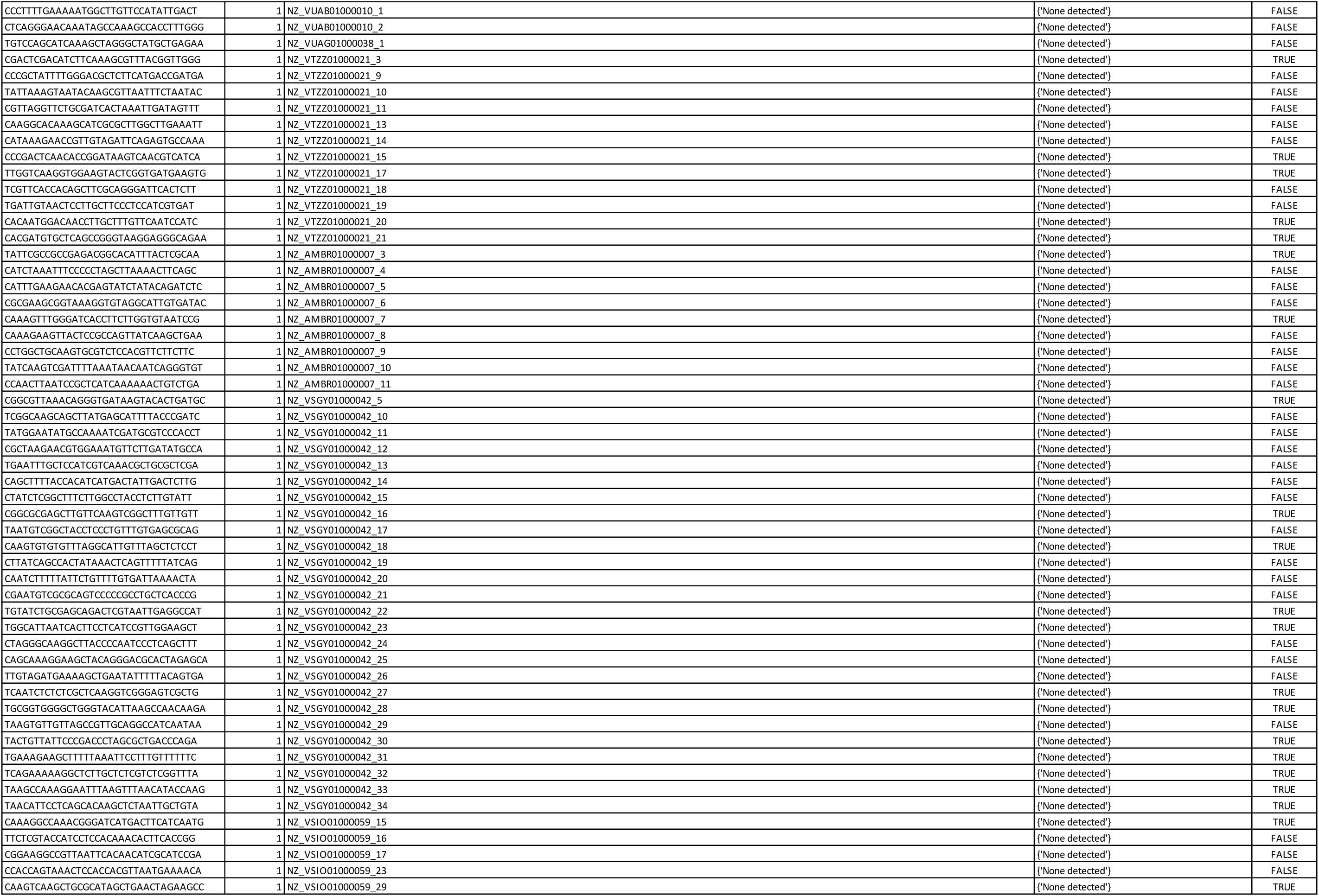
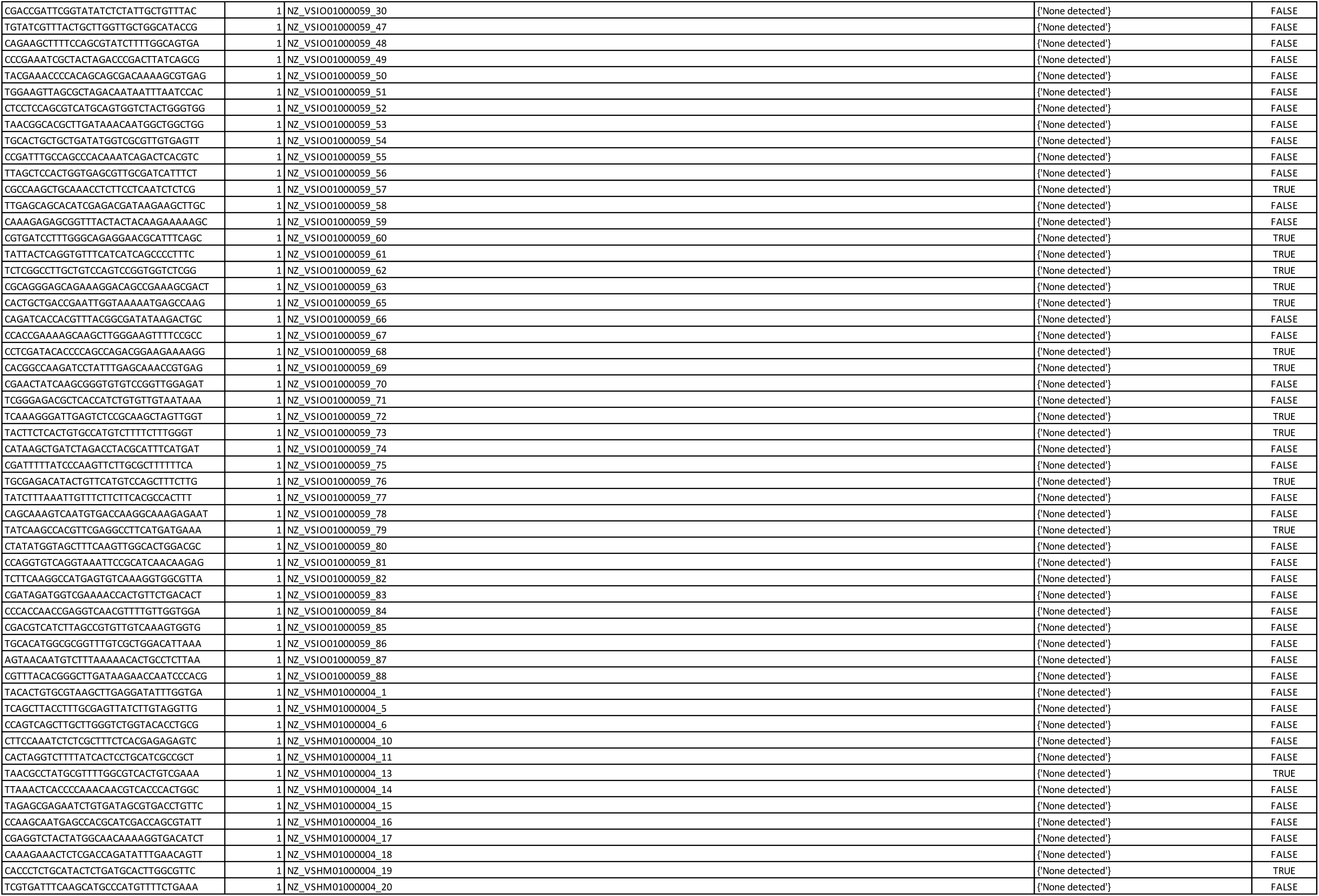
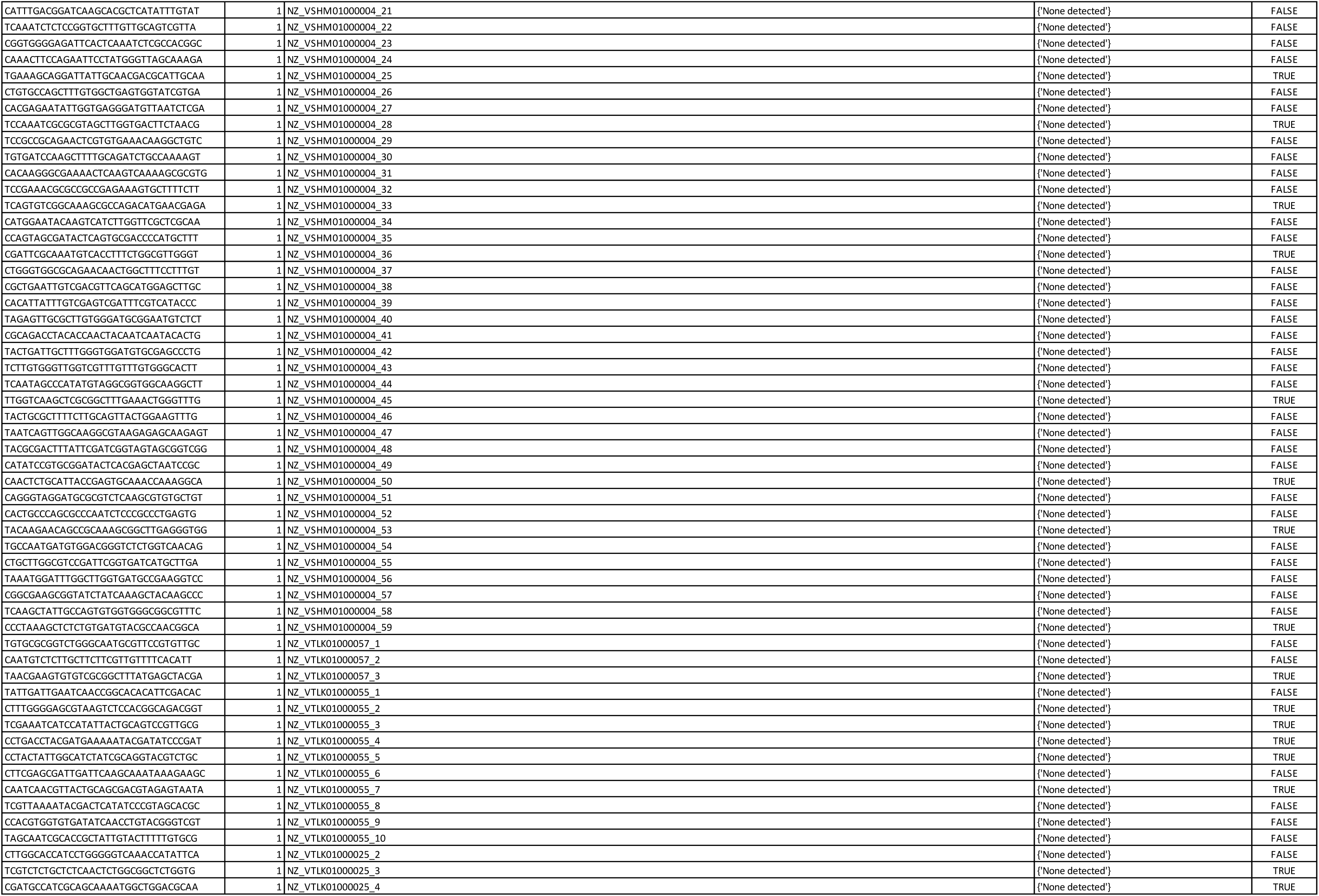
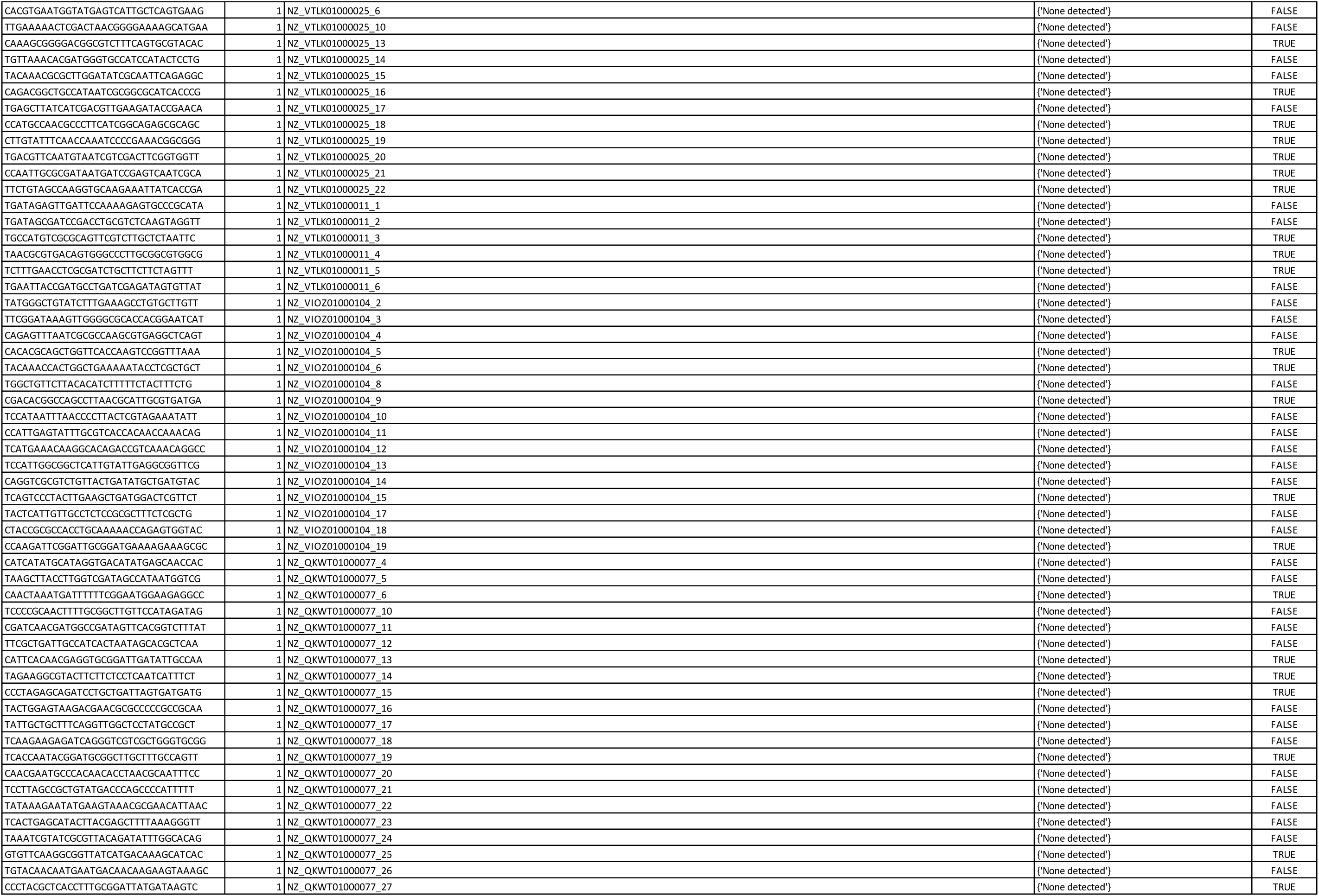
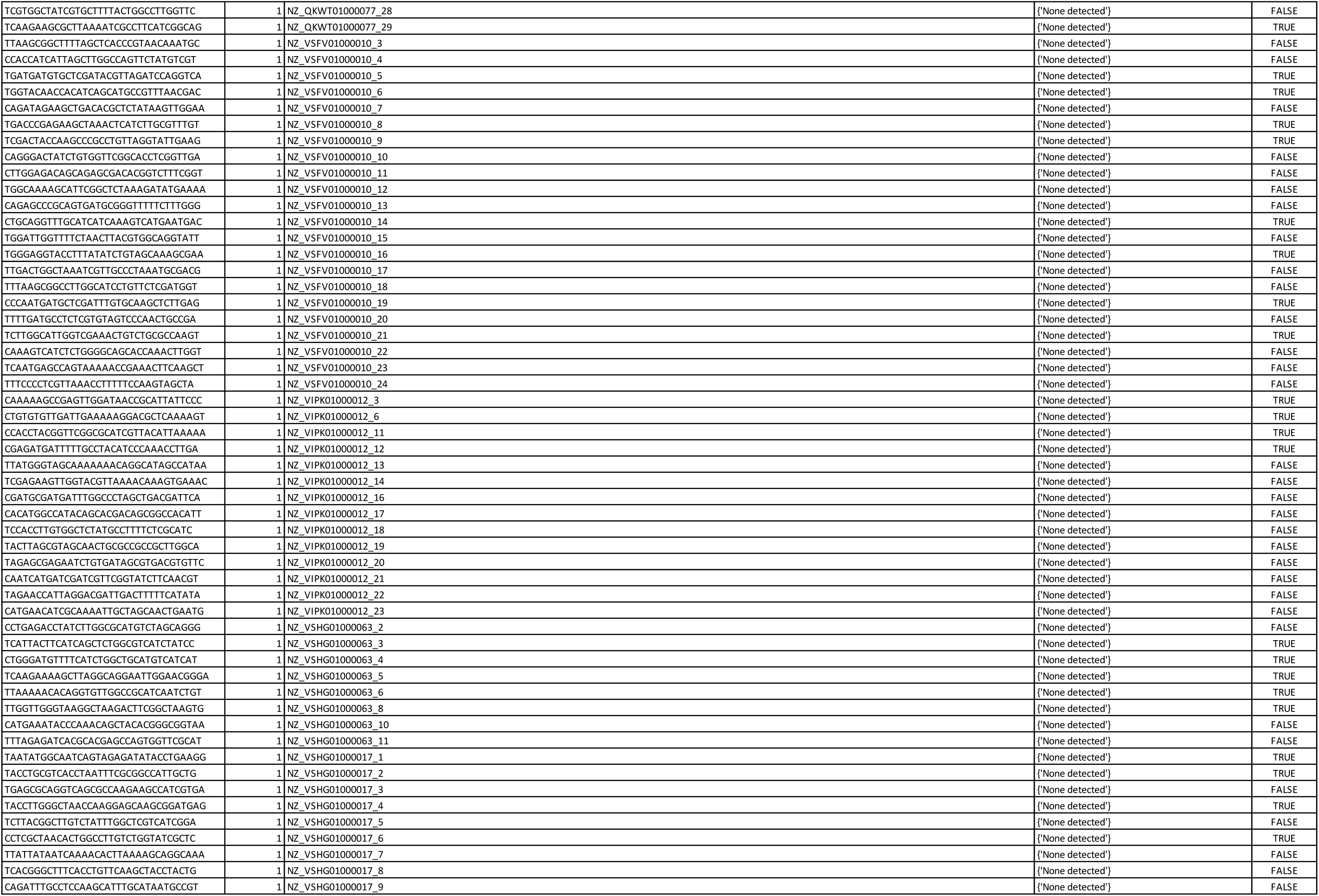
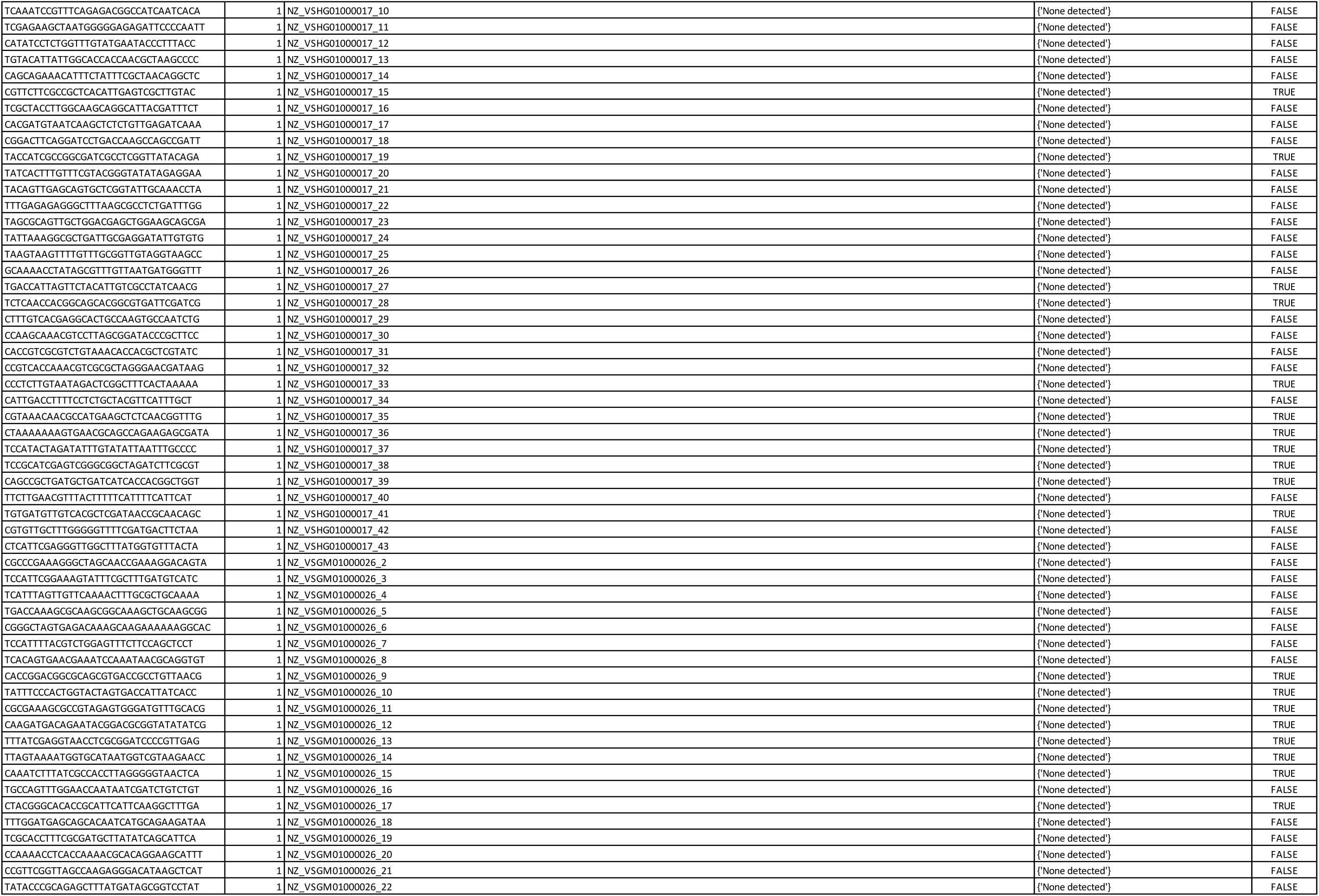
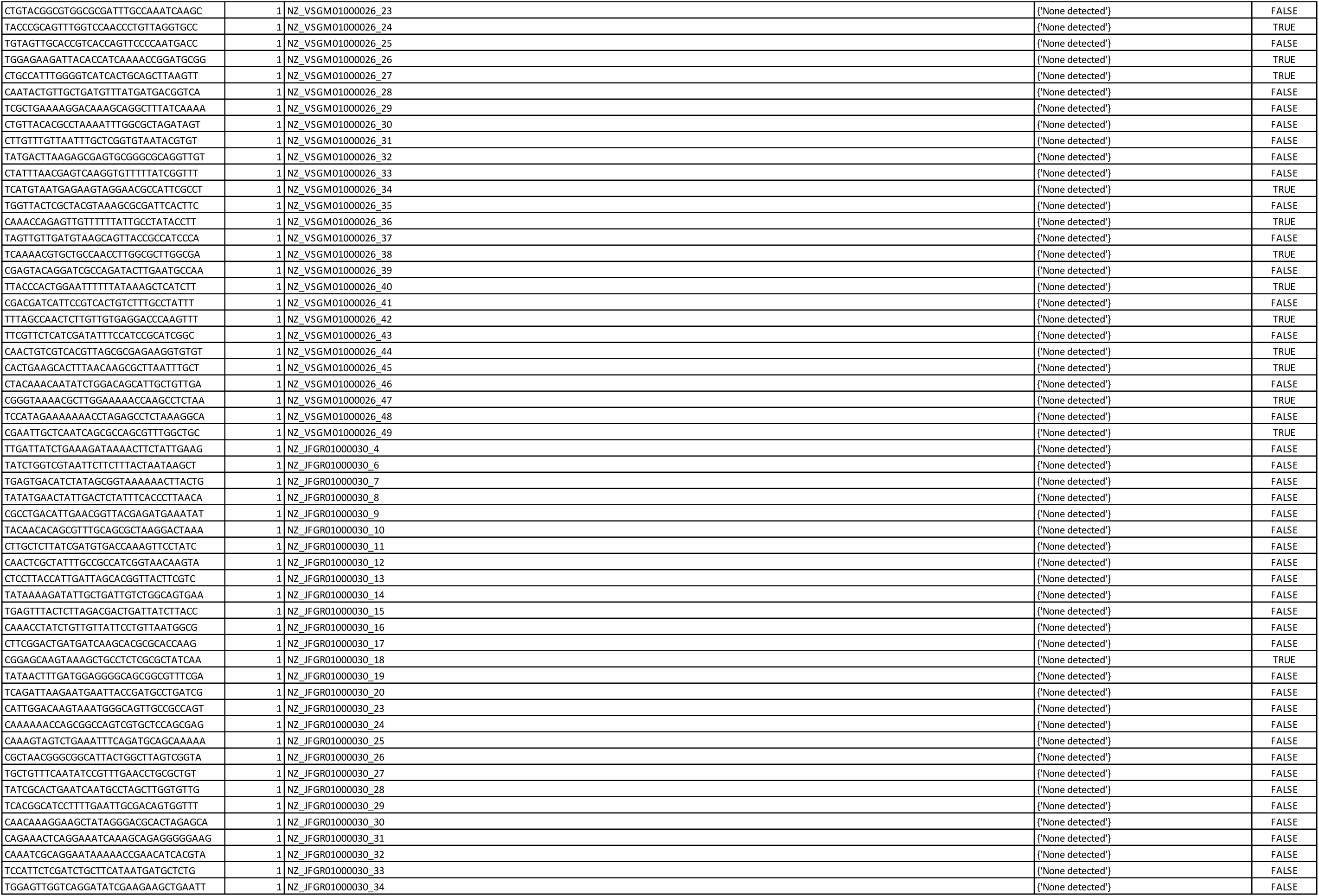
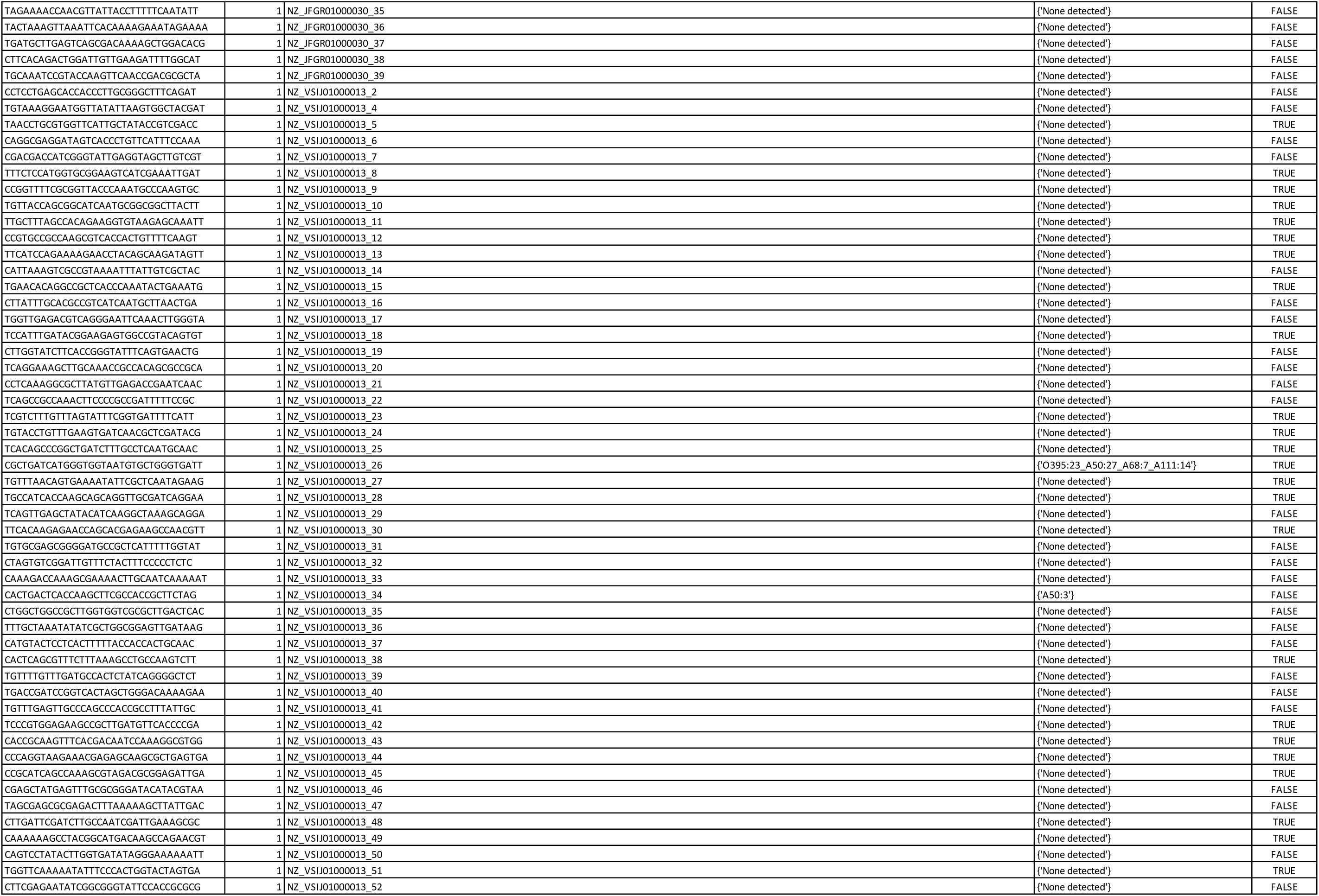
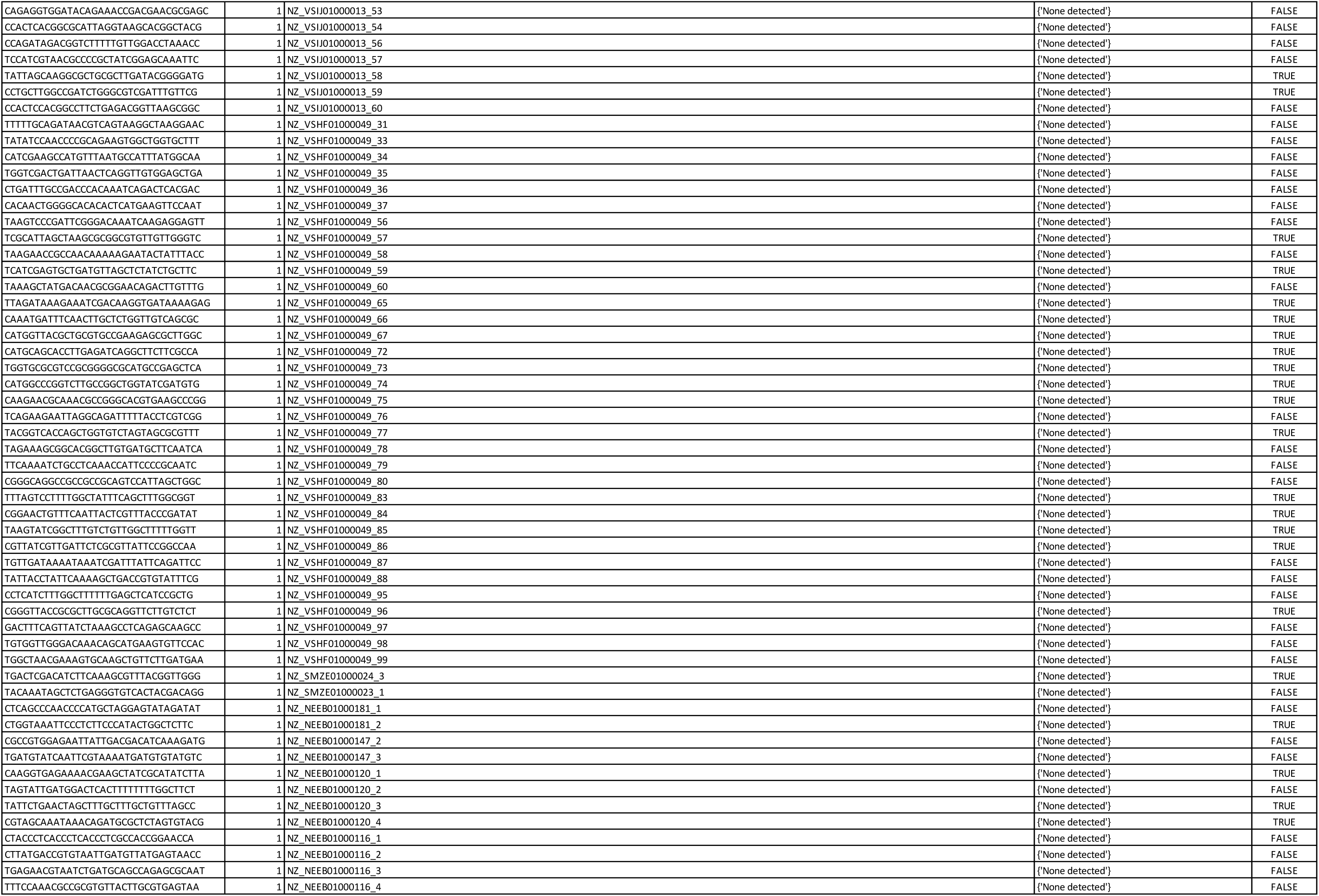
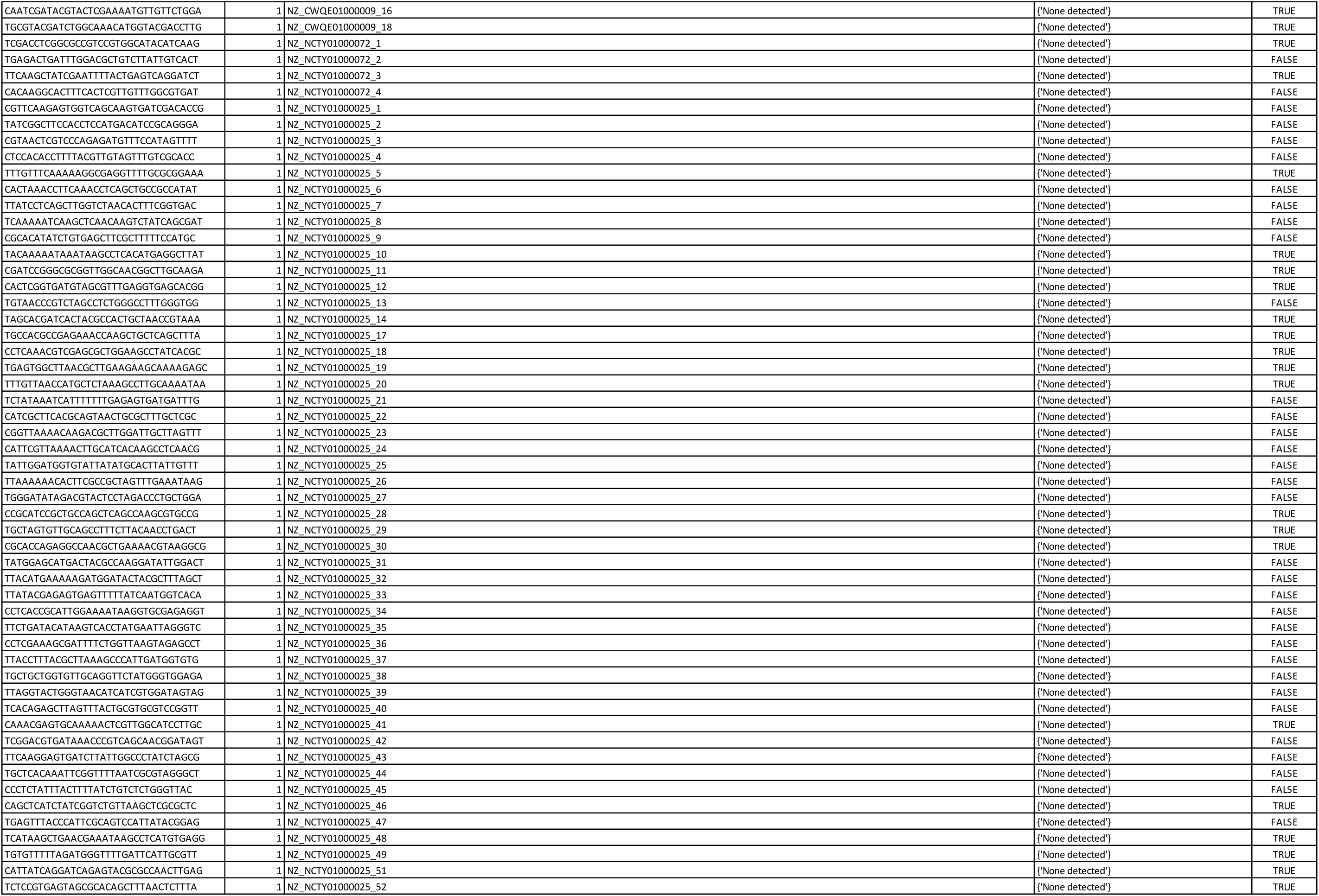
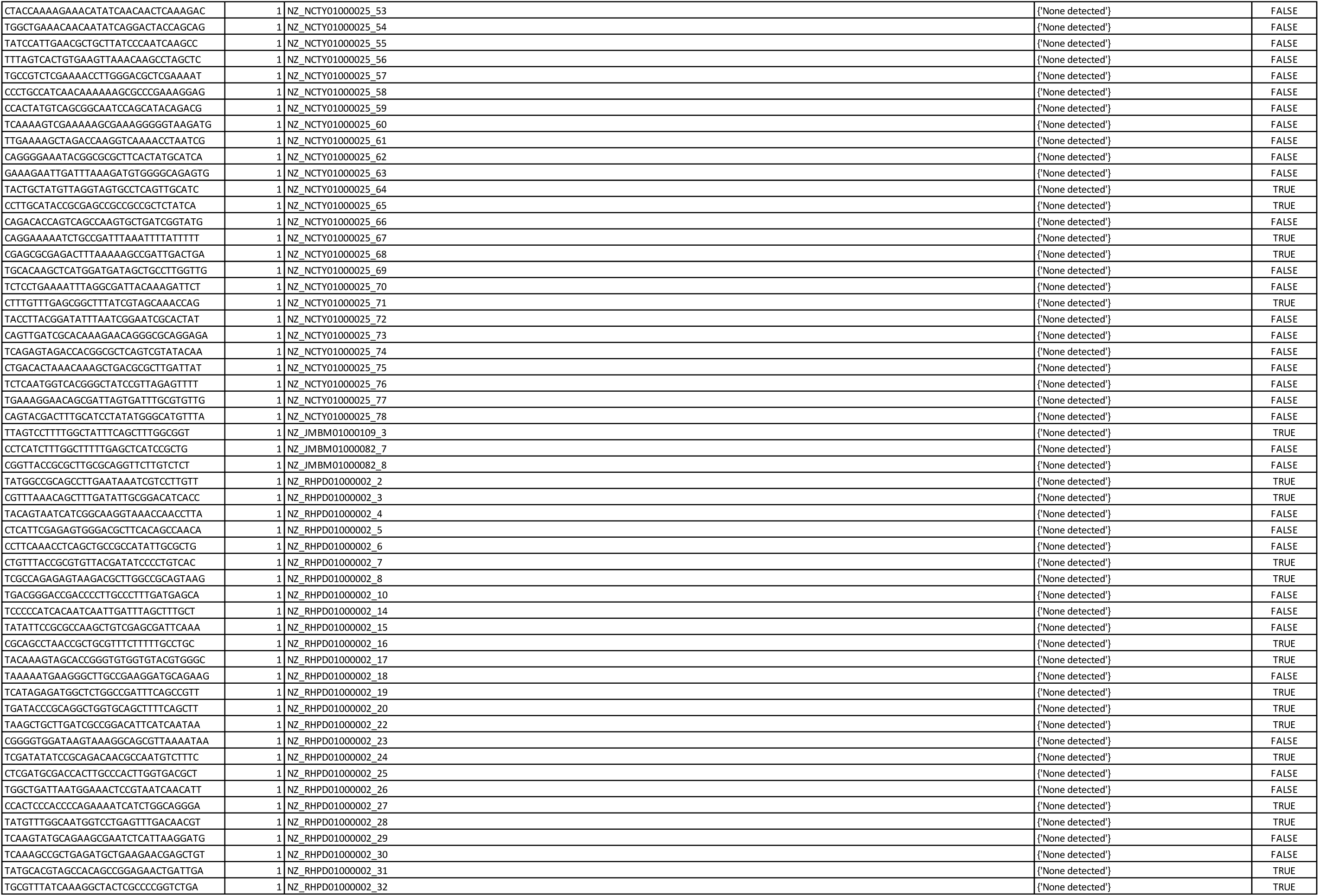
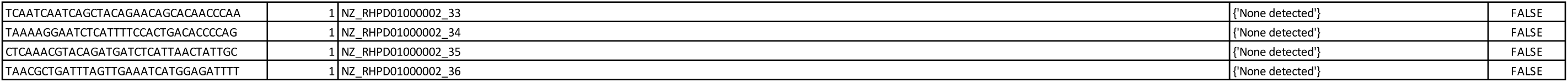
Frequency of mined spacers in identified *V. cholerae* sequences harboring the Type I-E CRISPR/Cas system.

**Supplementary Table 4:**
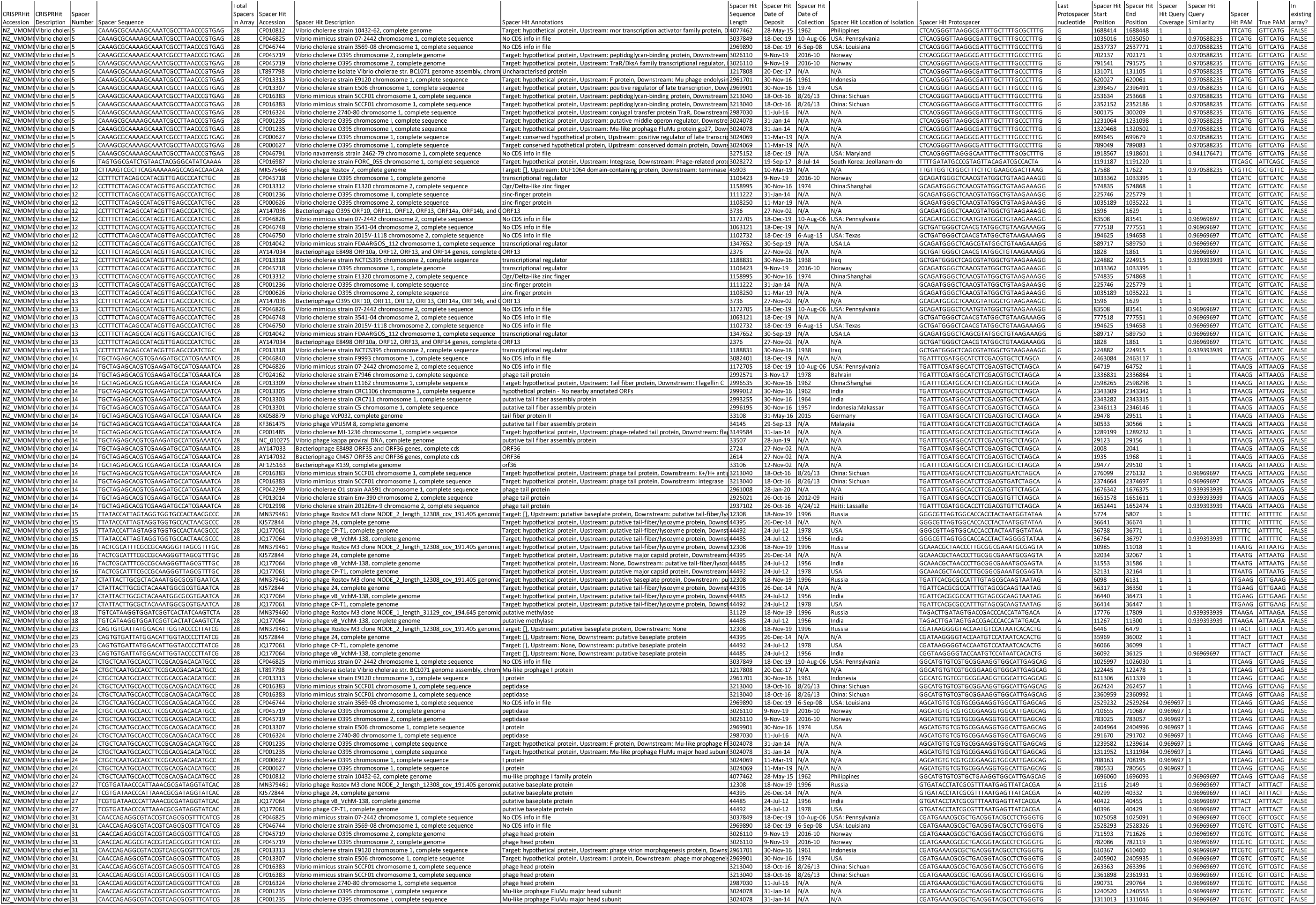
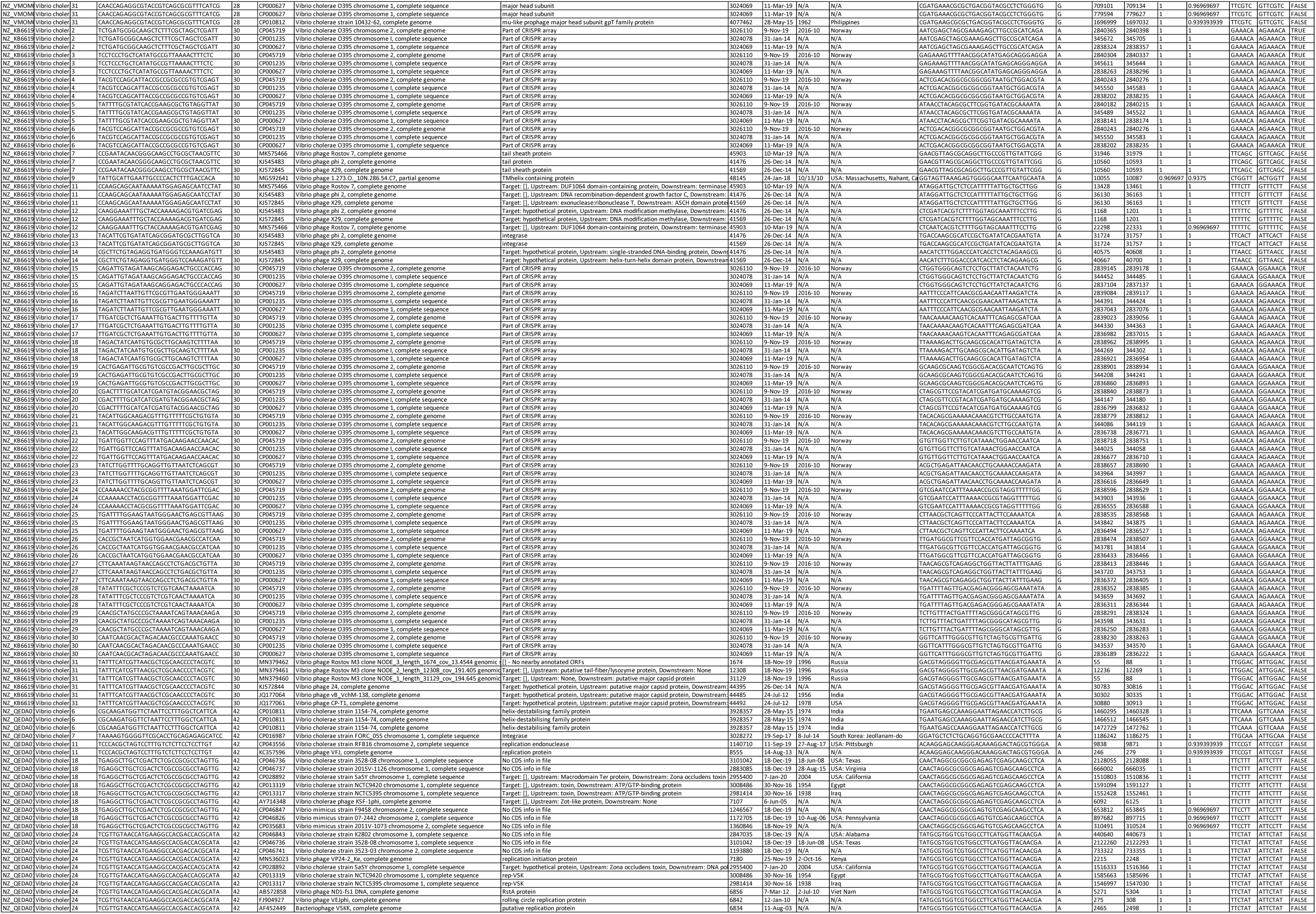
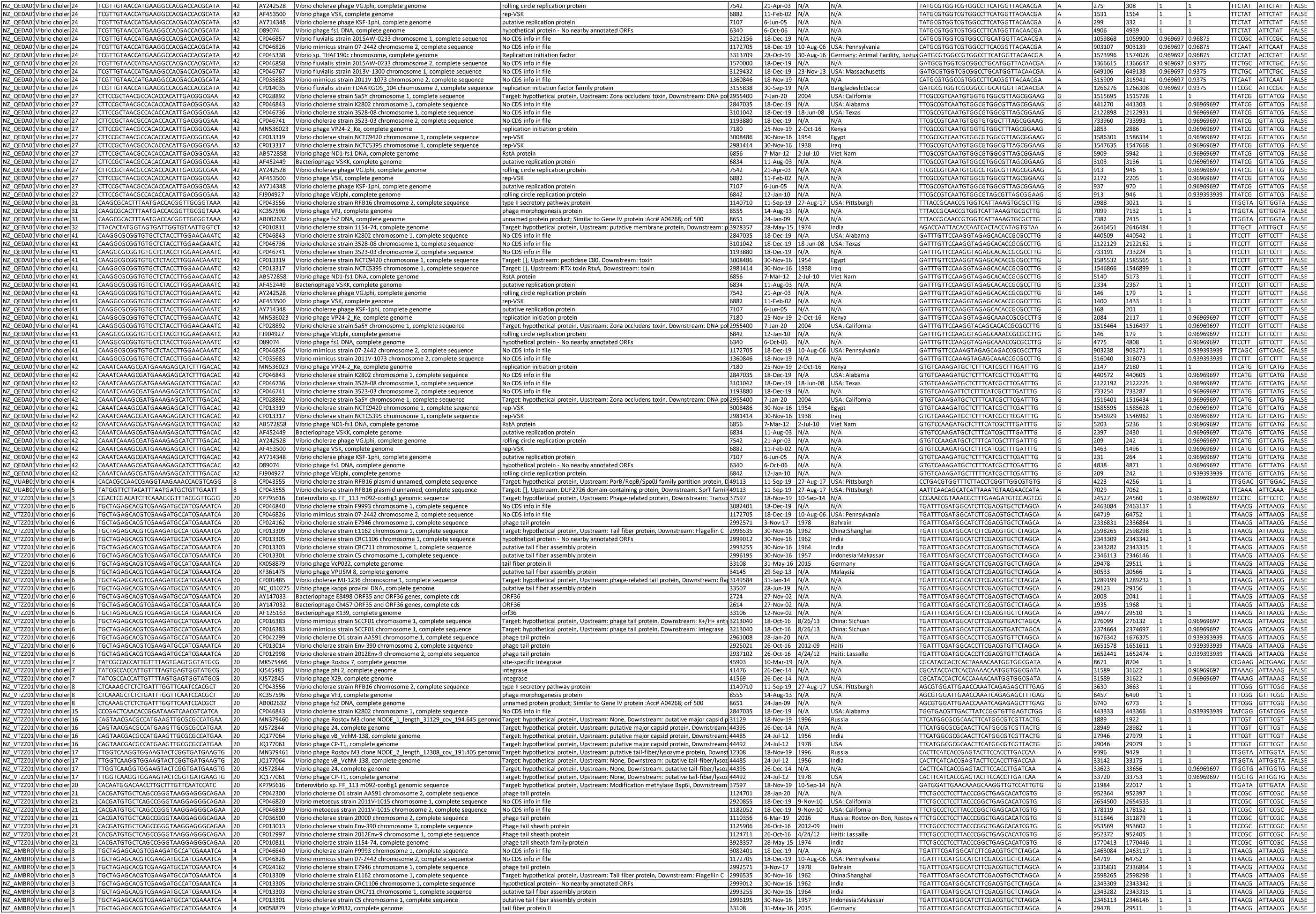
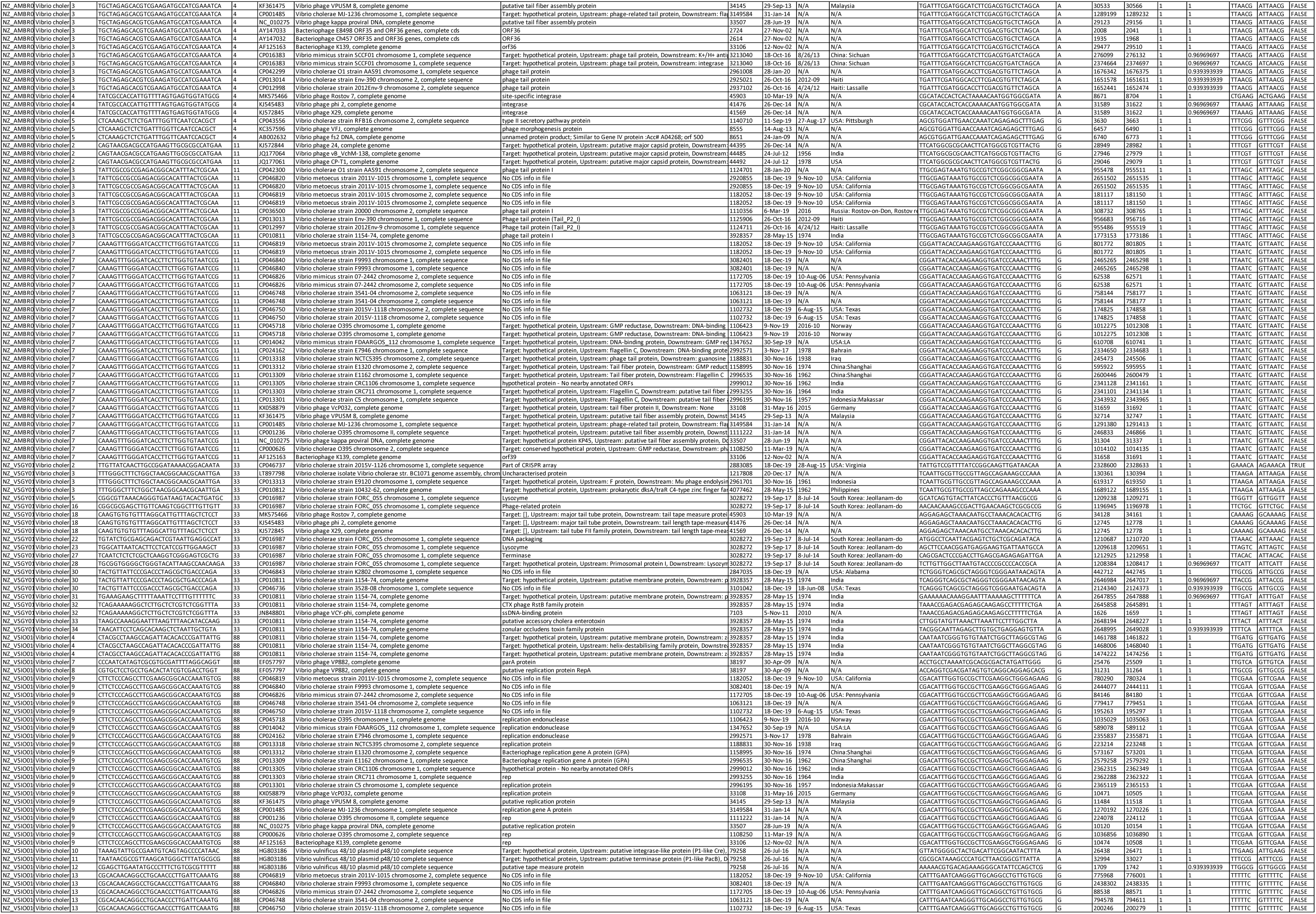
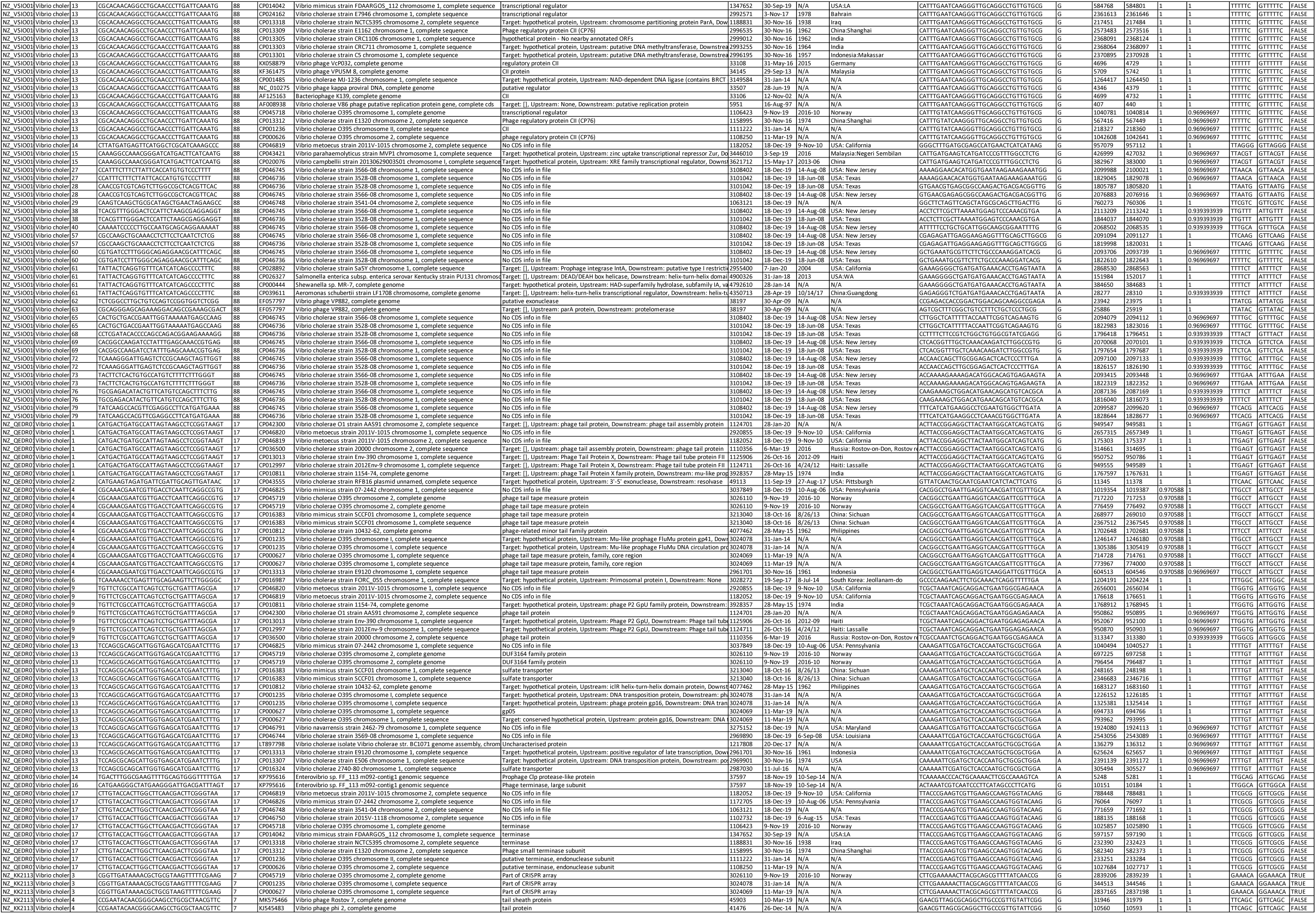
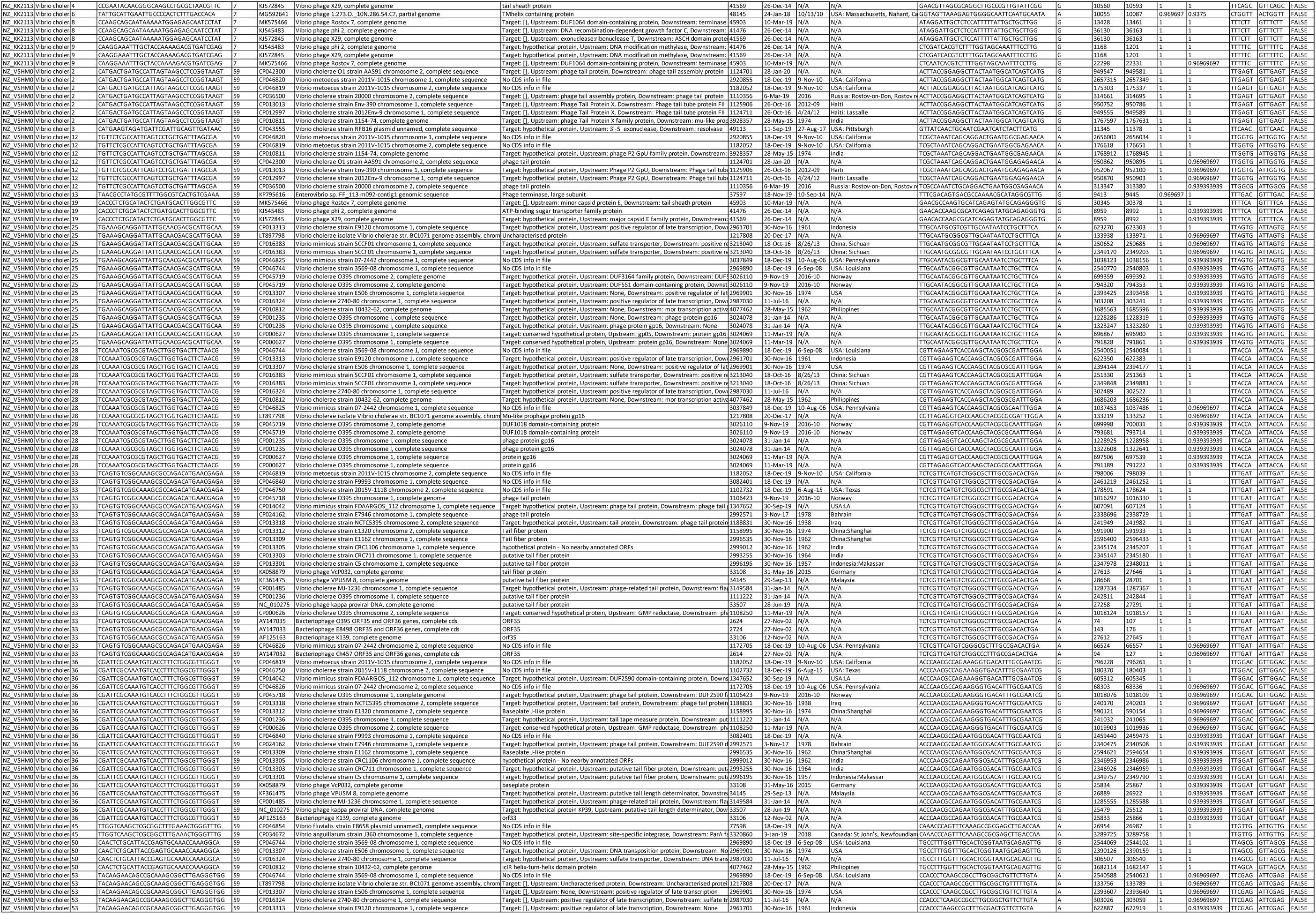
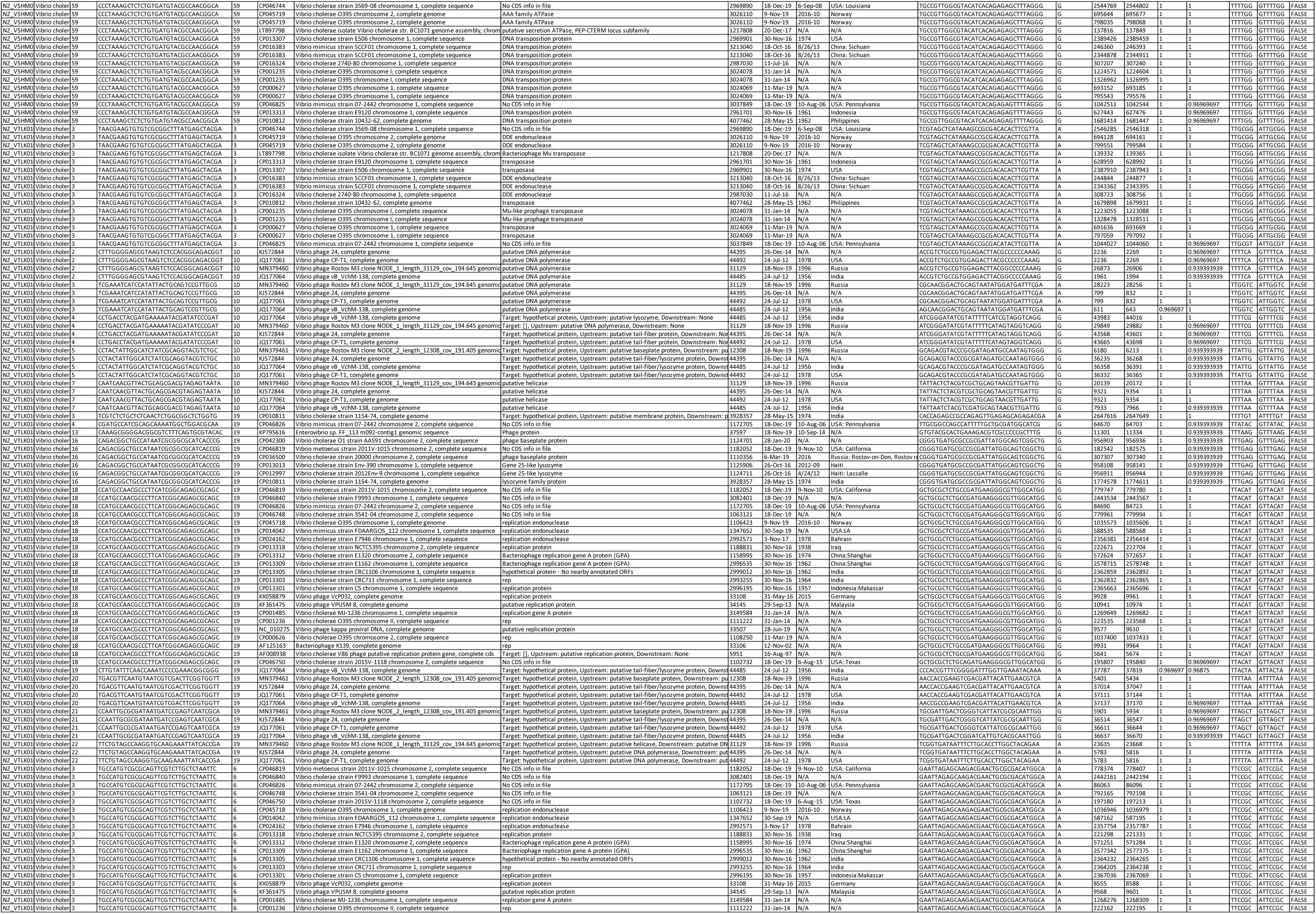
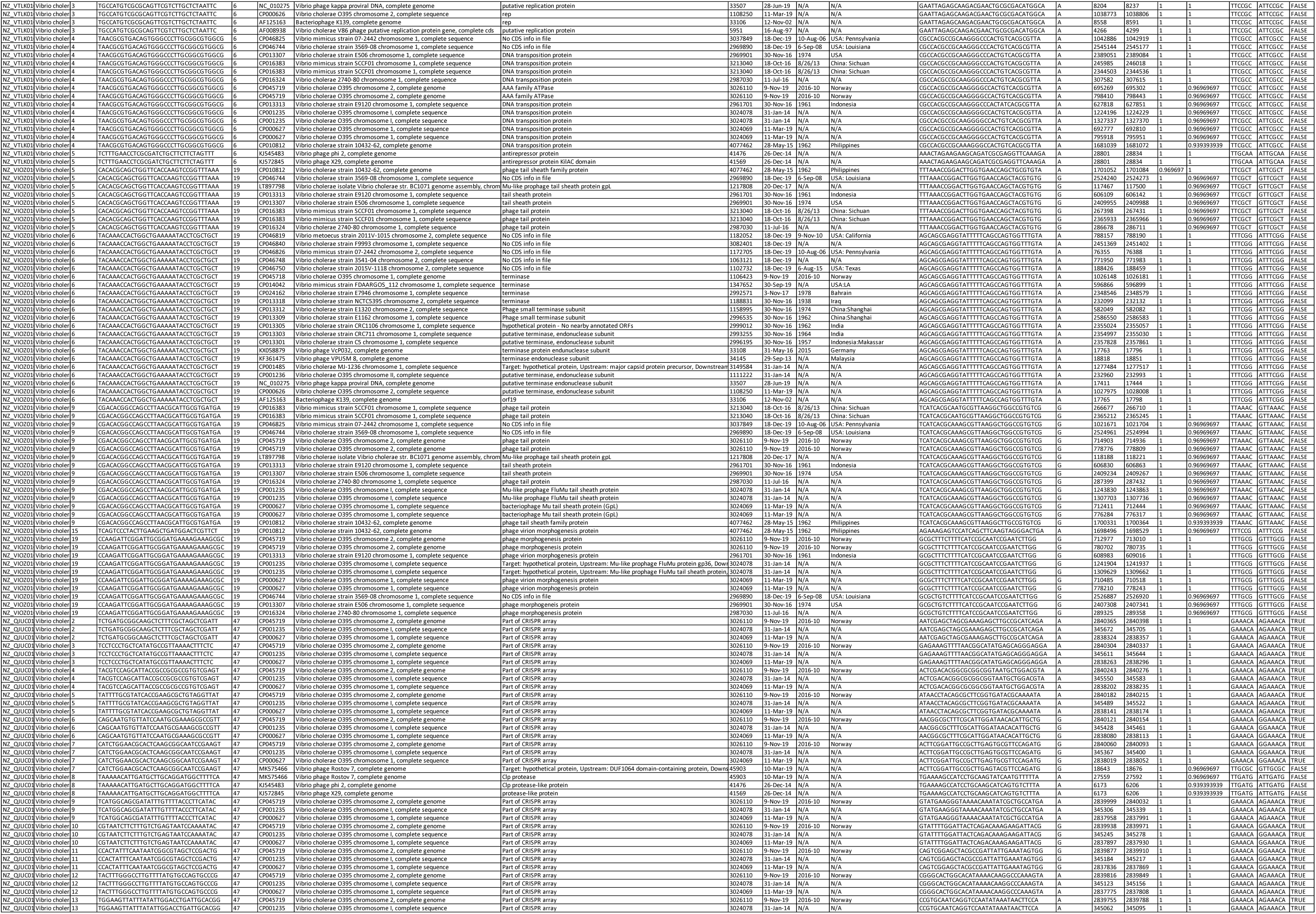
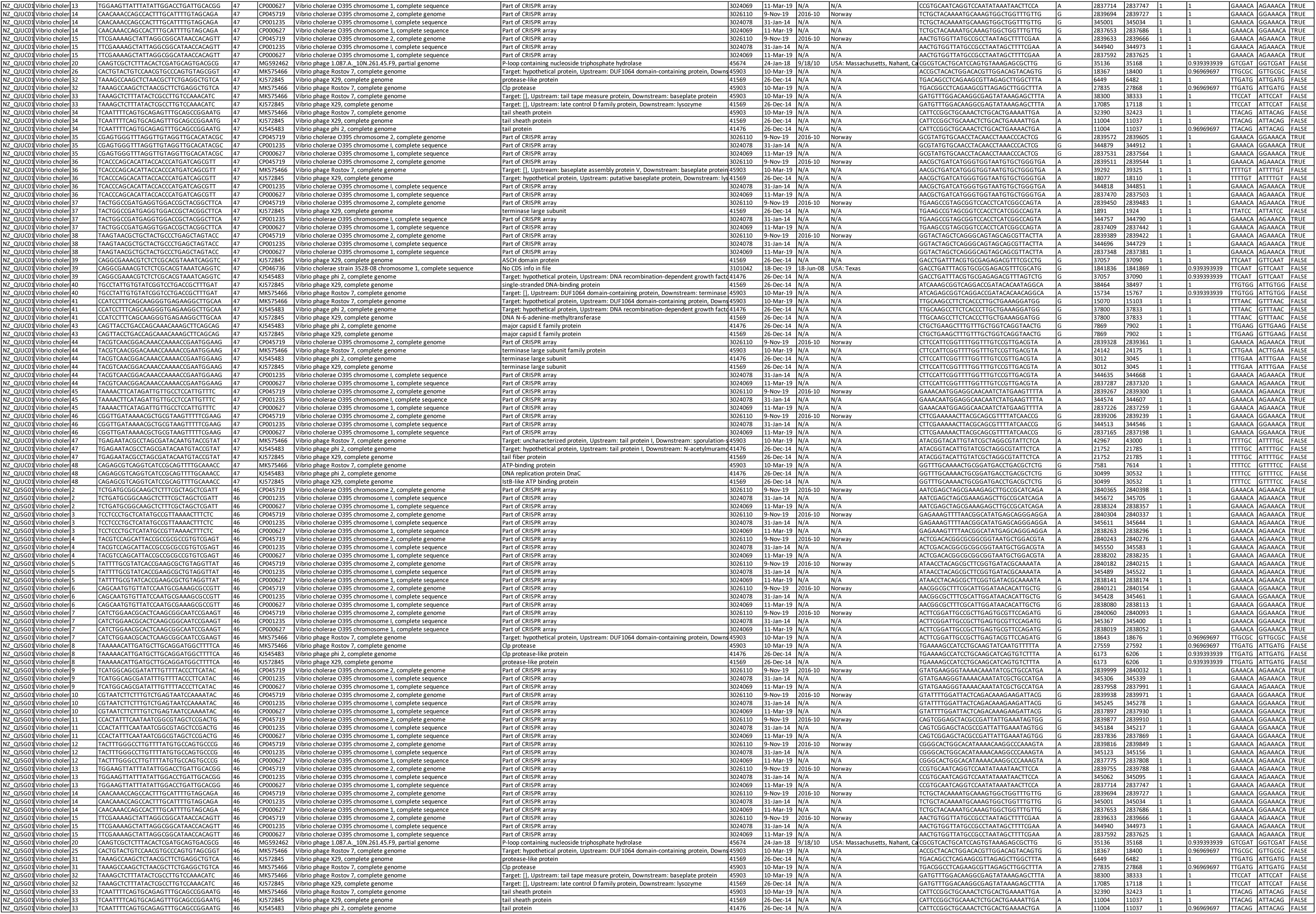
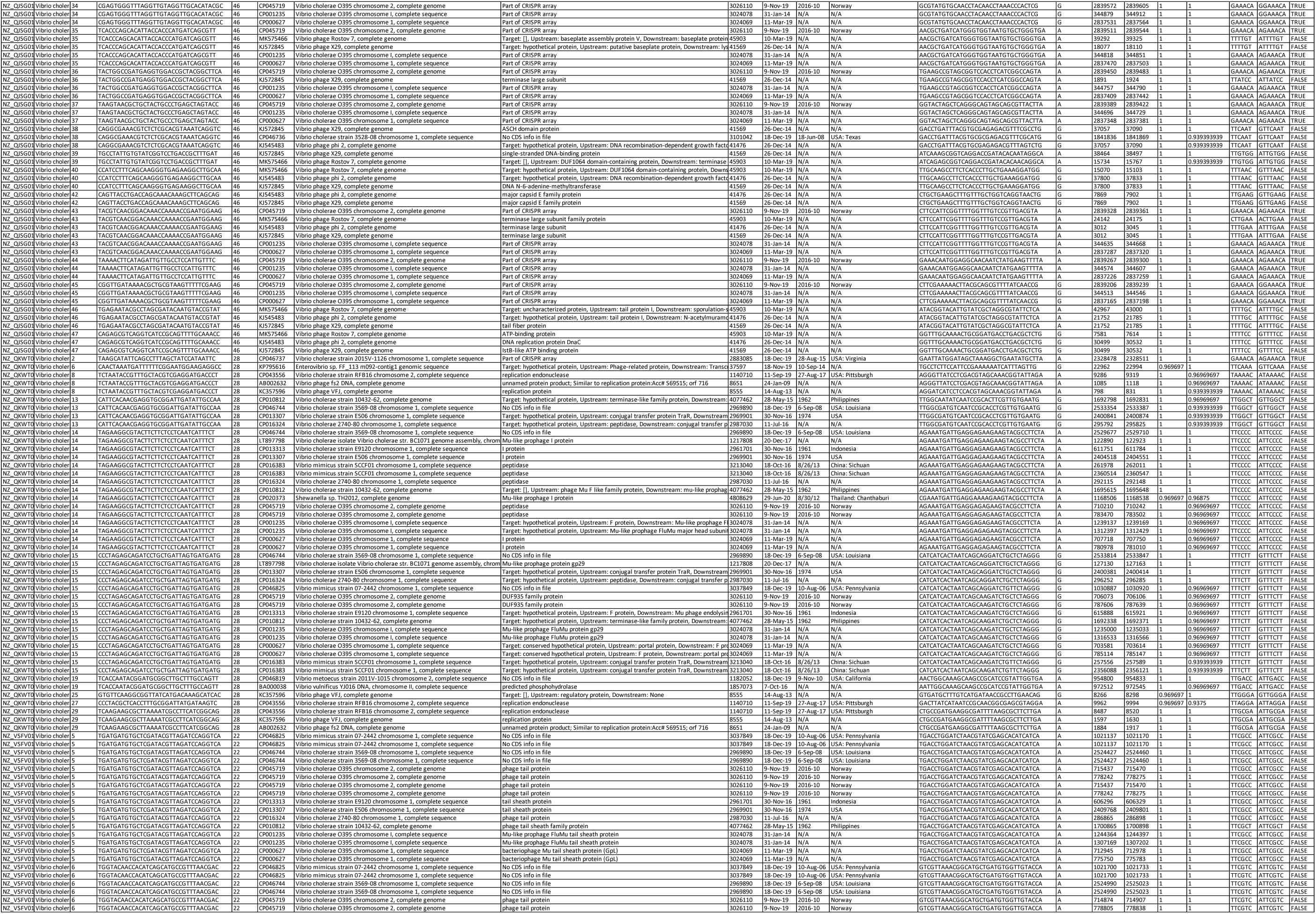
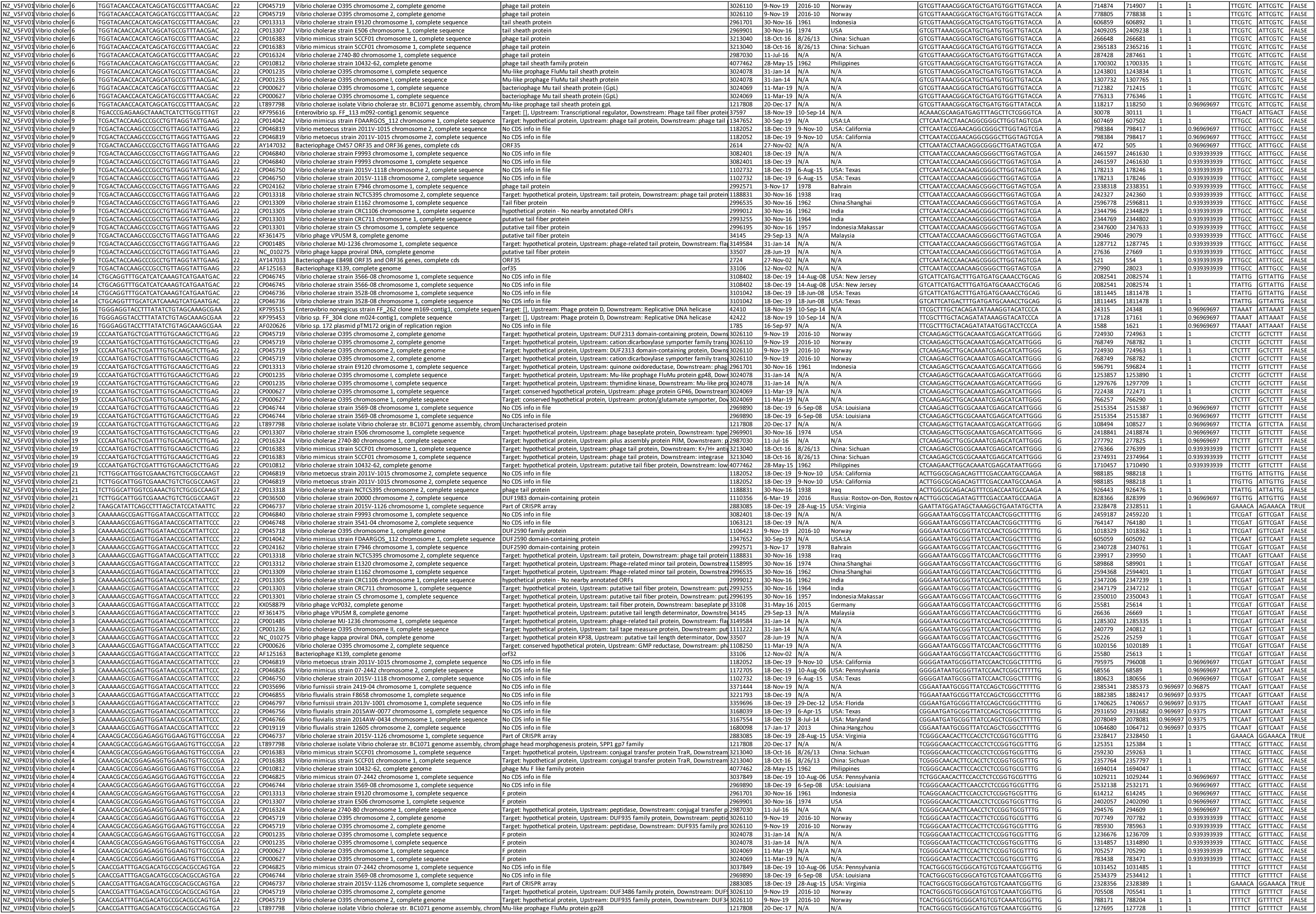
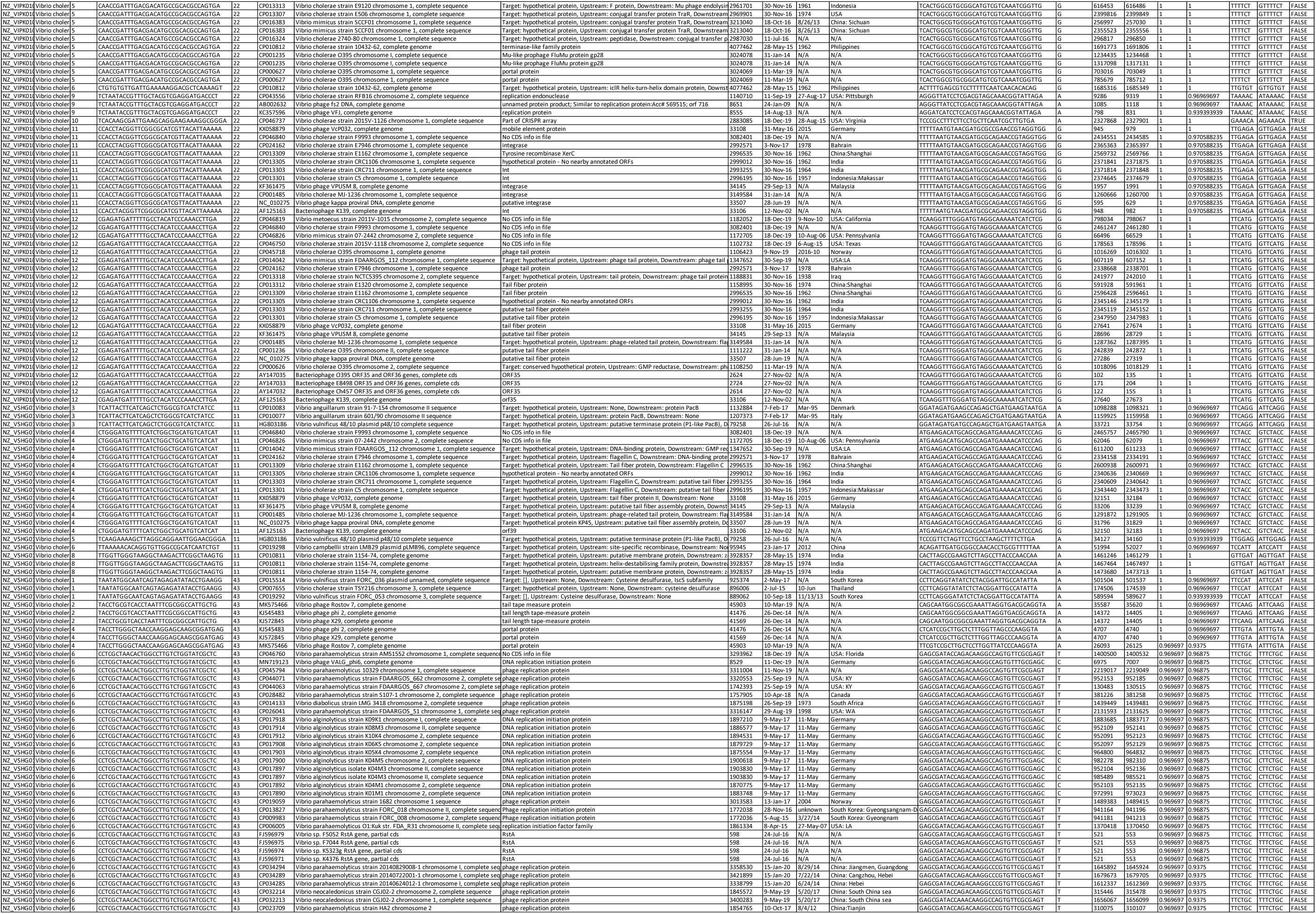
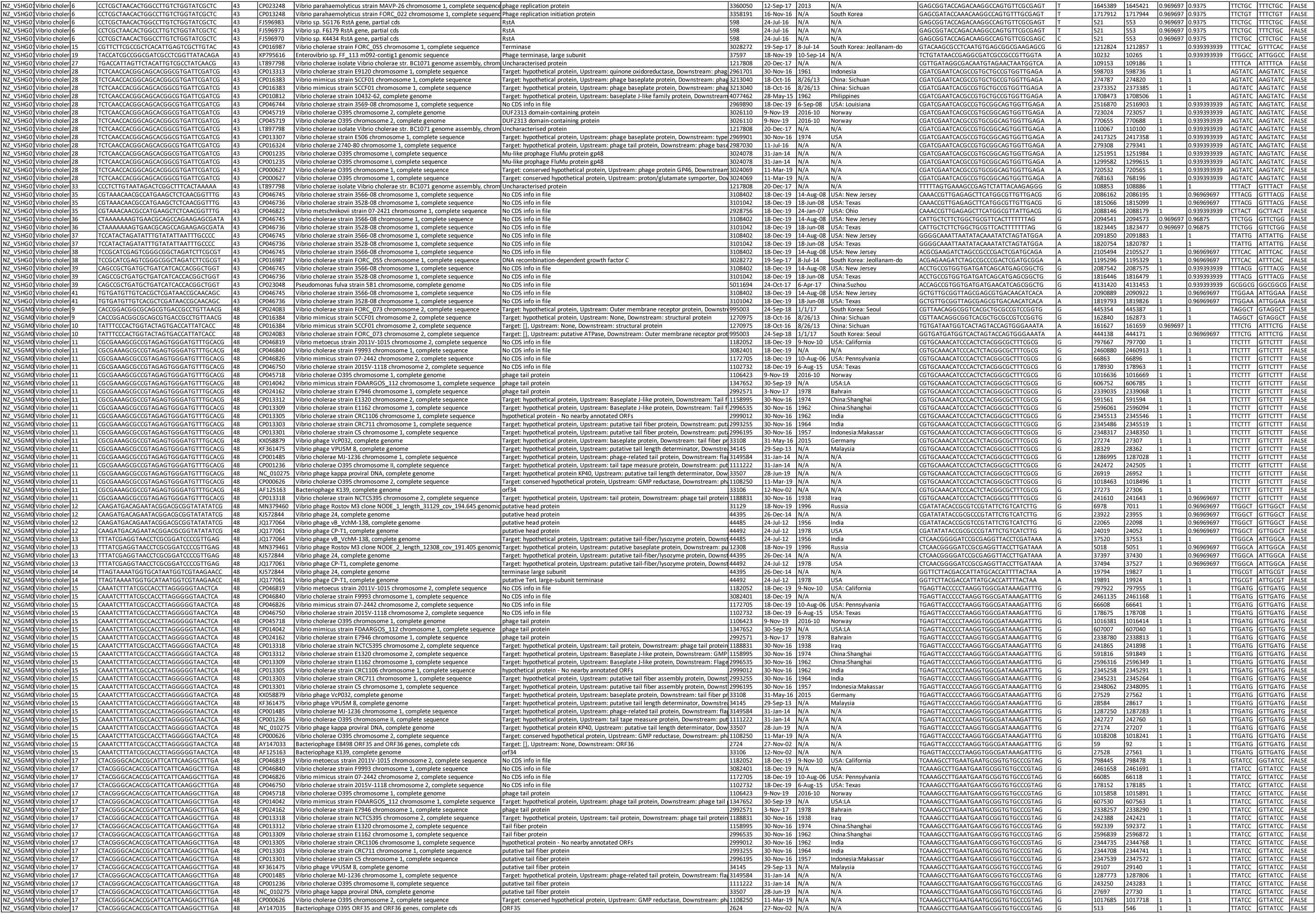
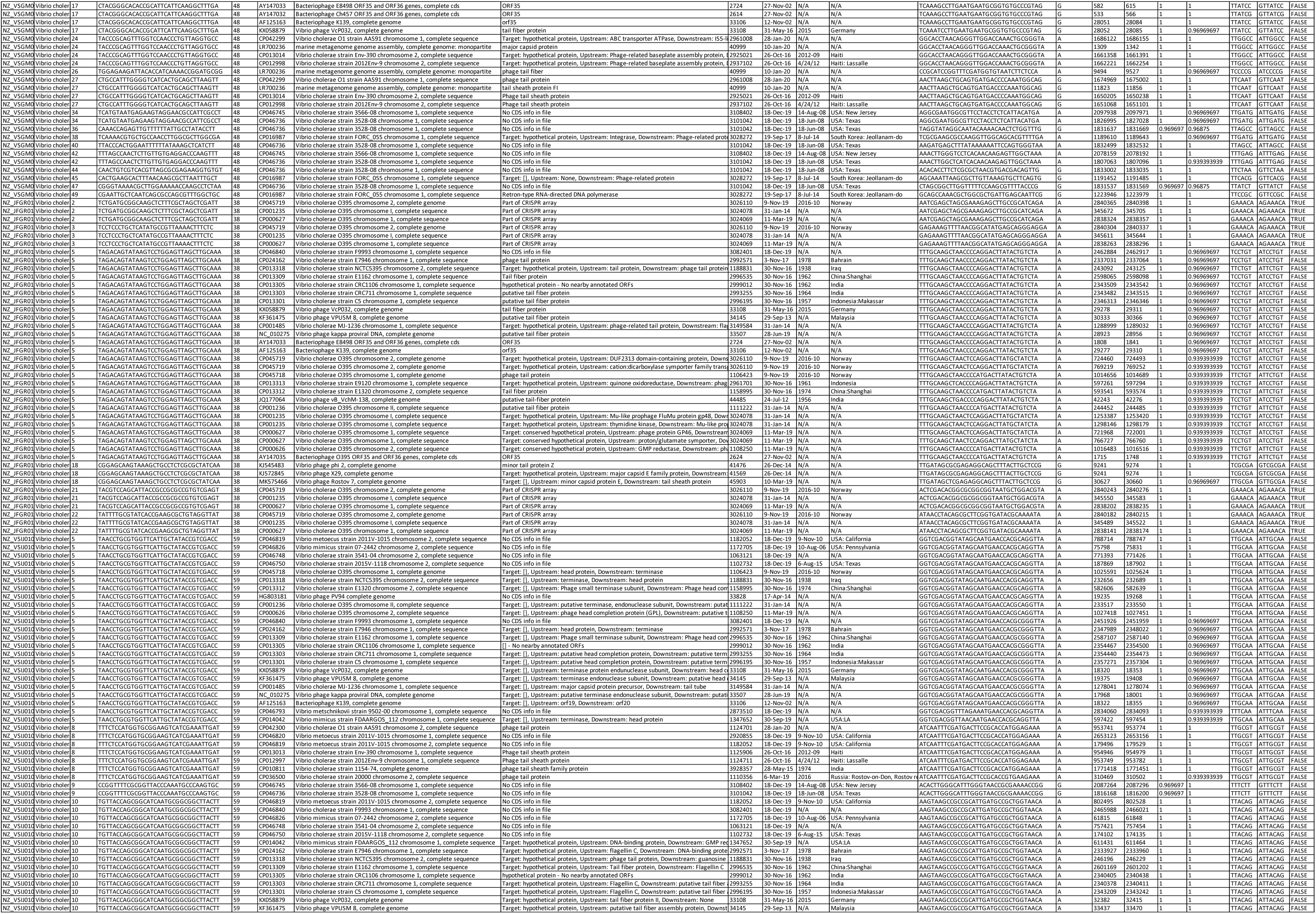
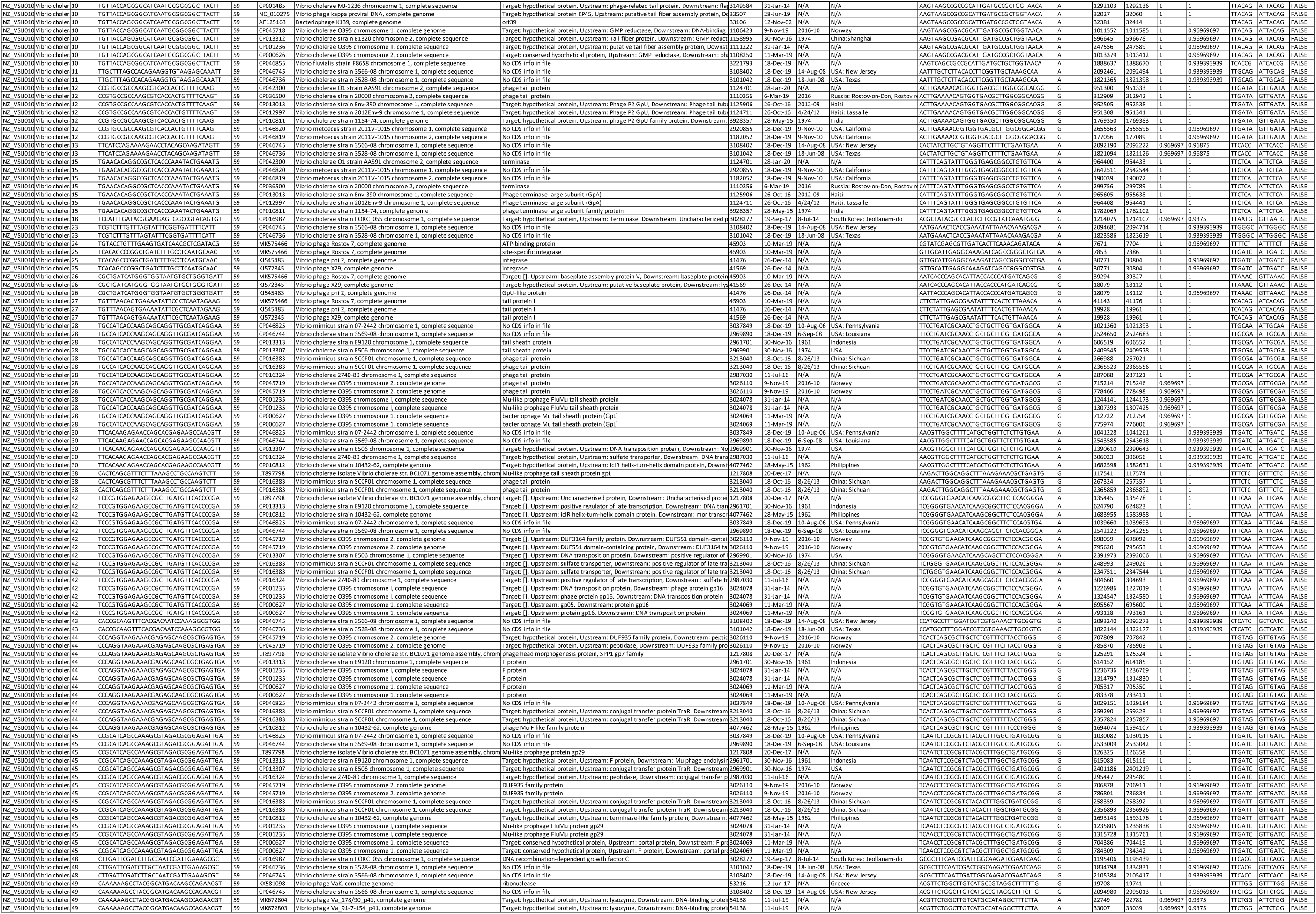
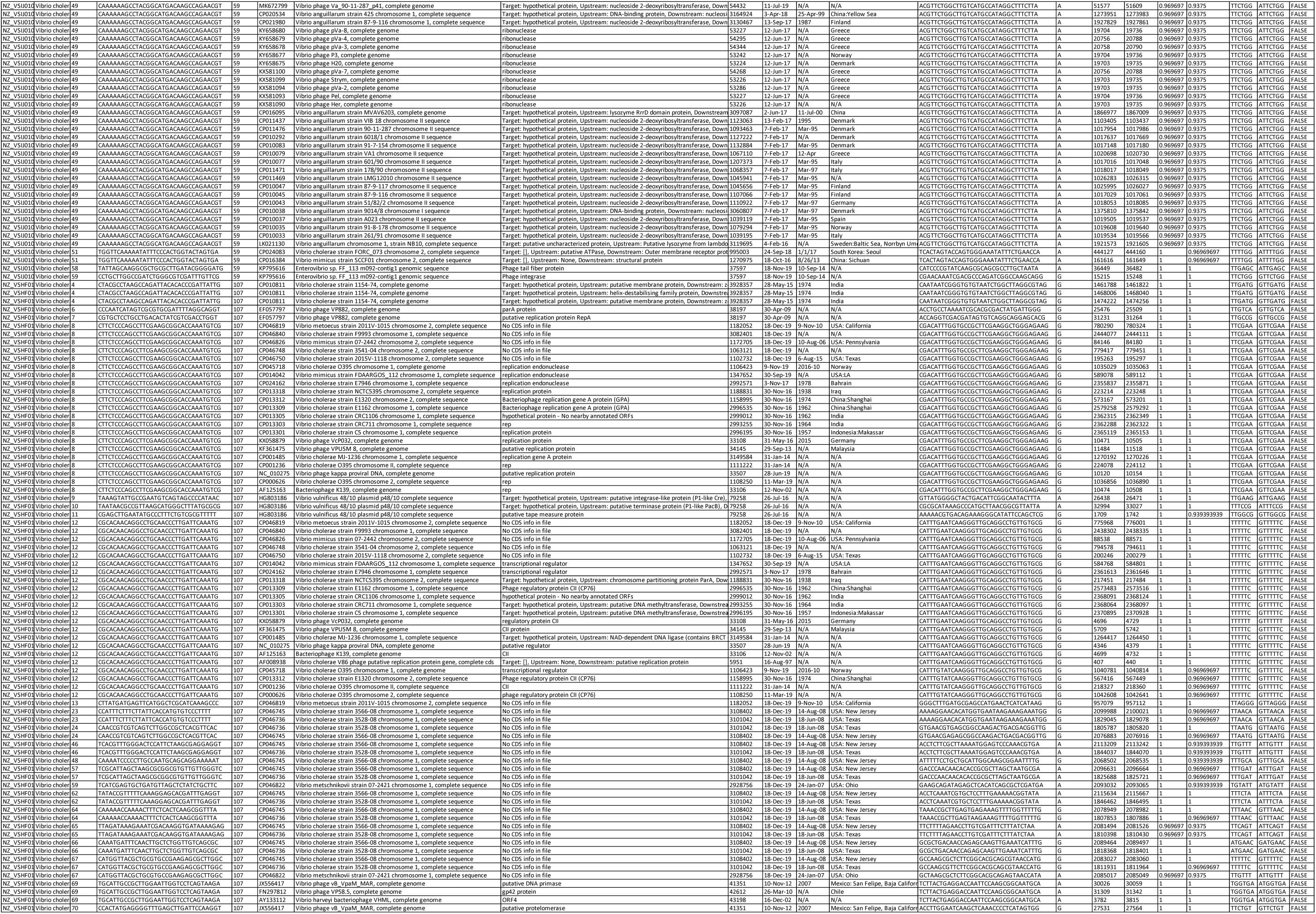
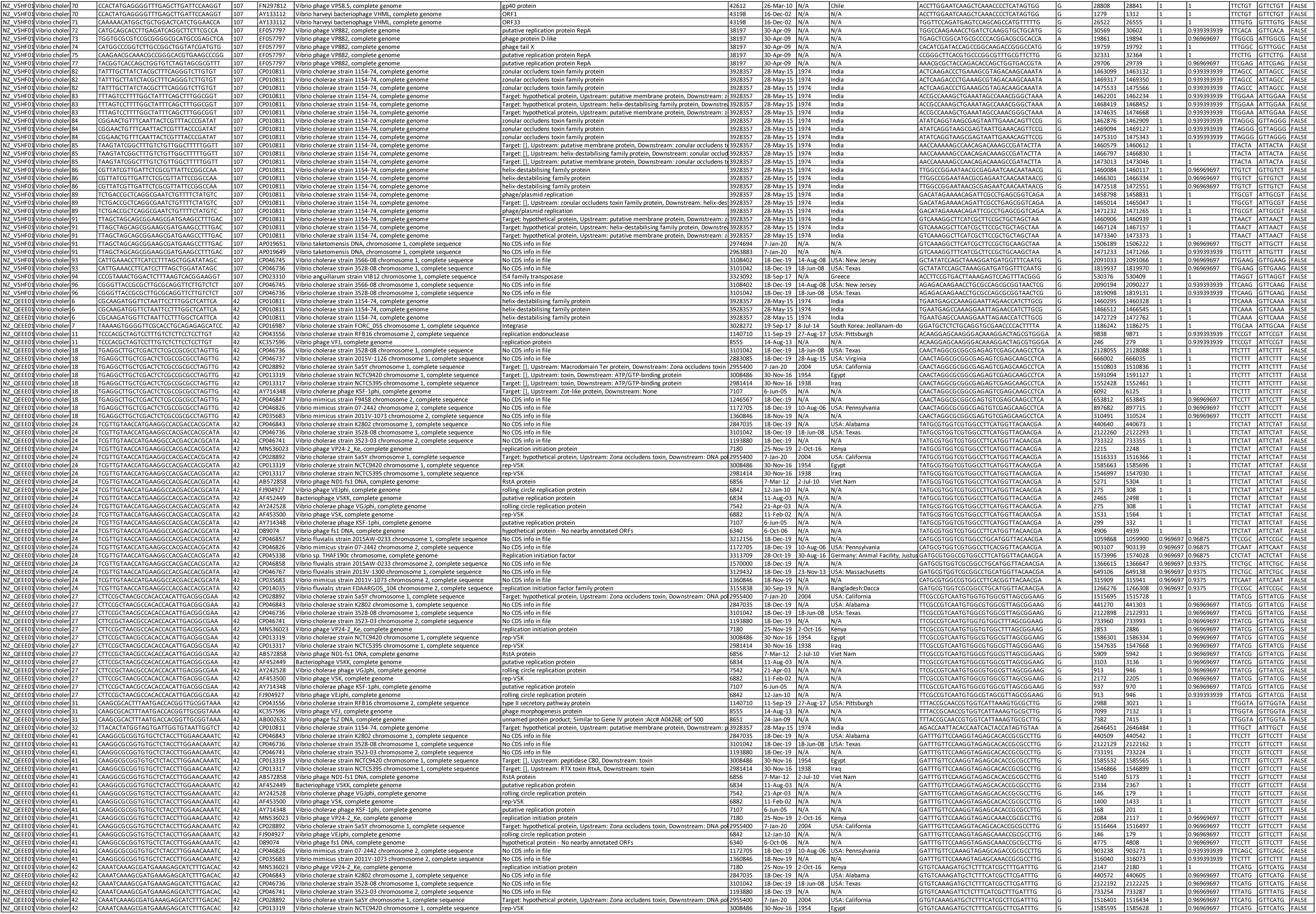
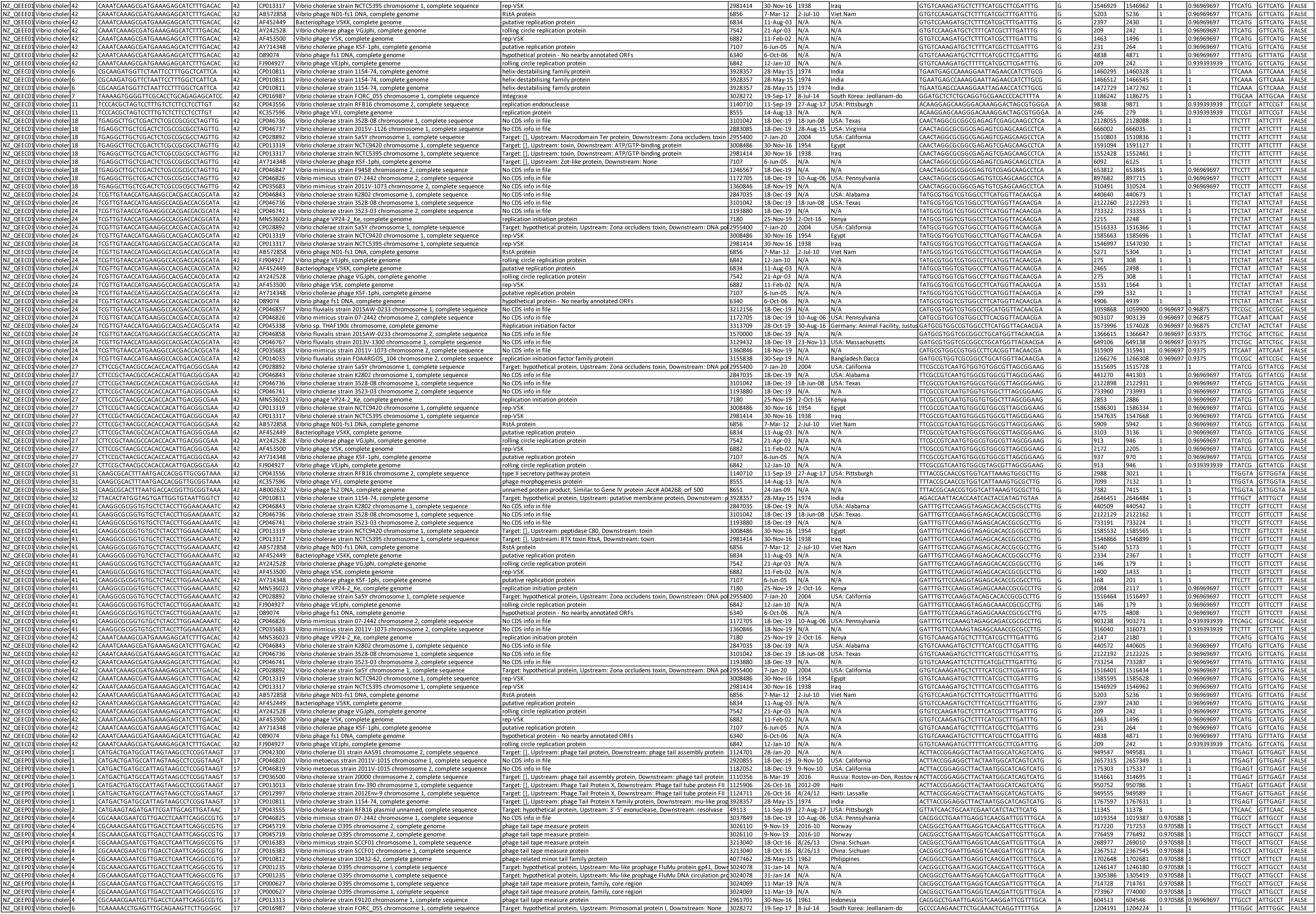
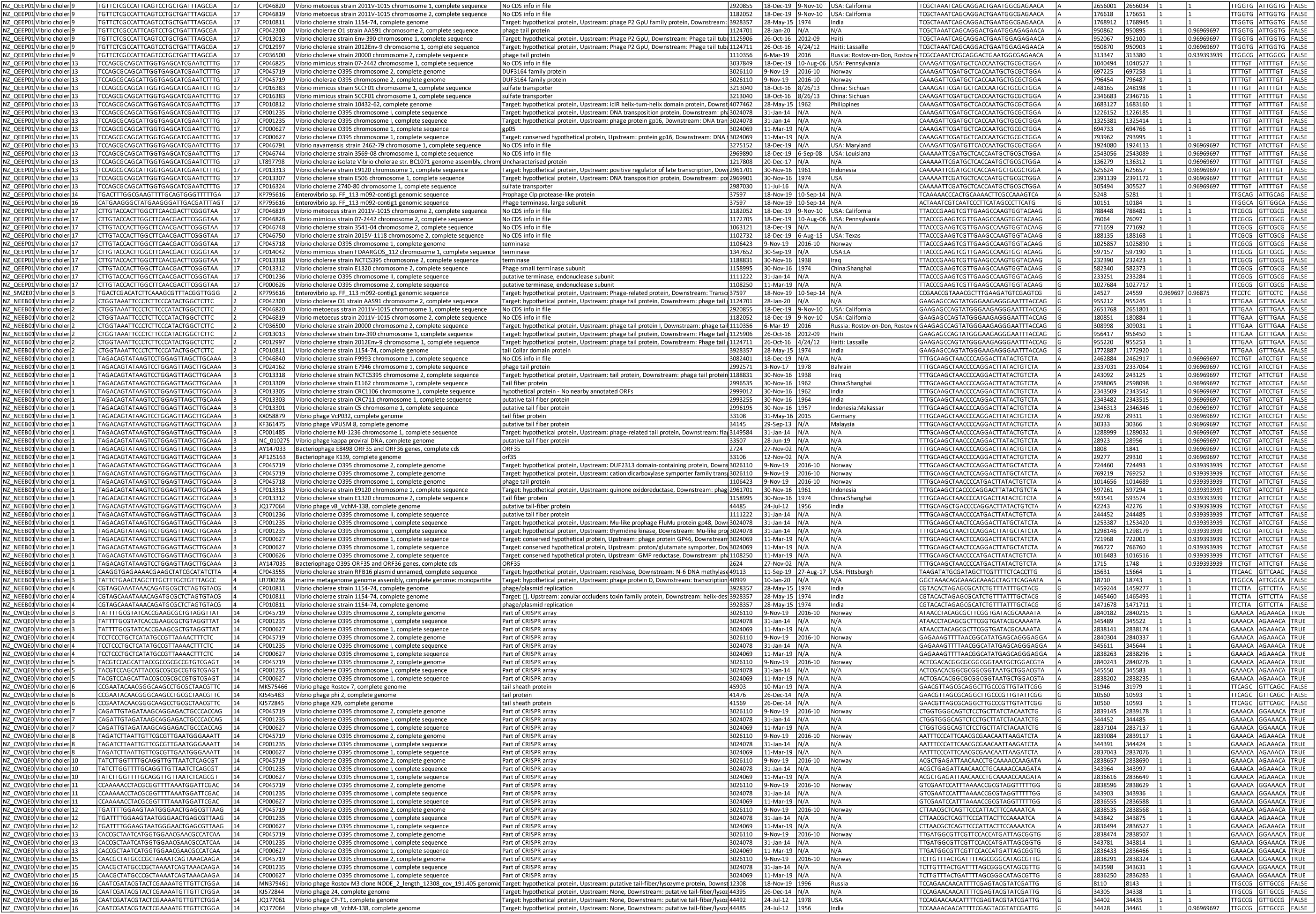
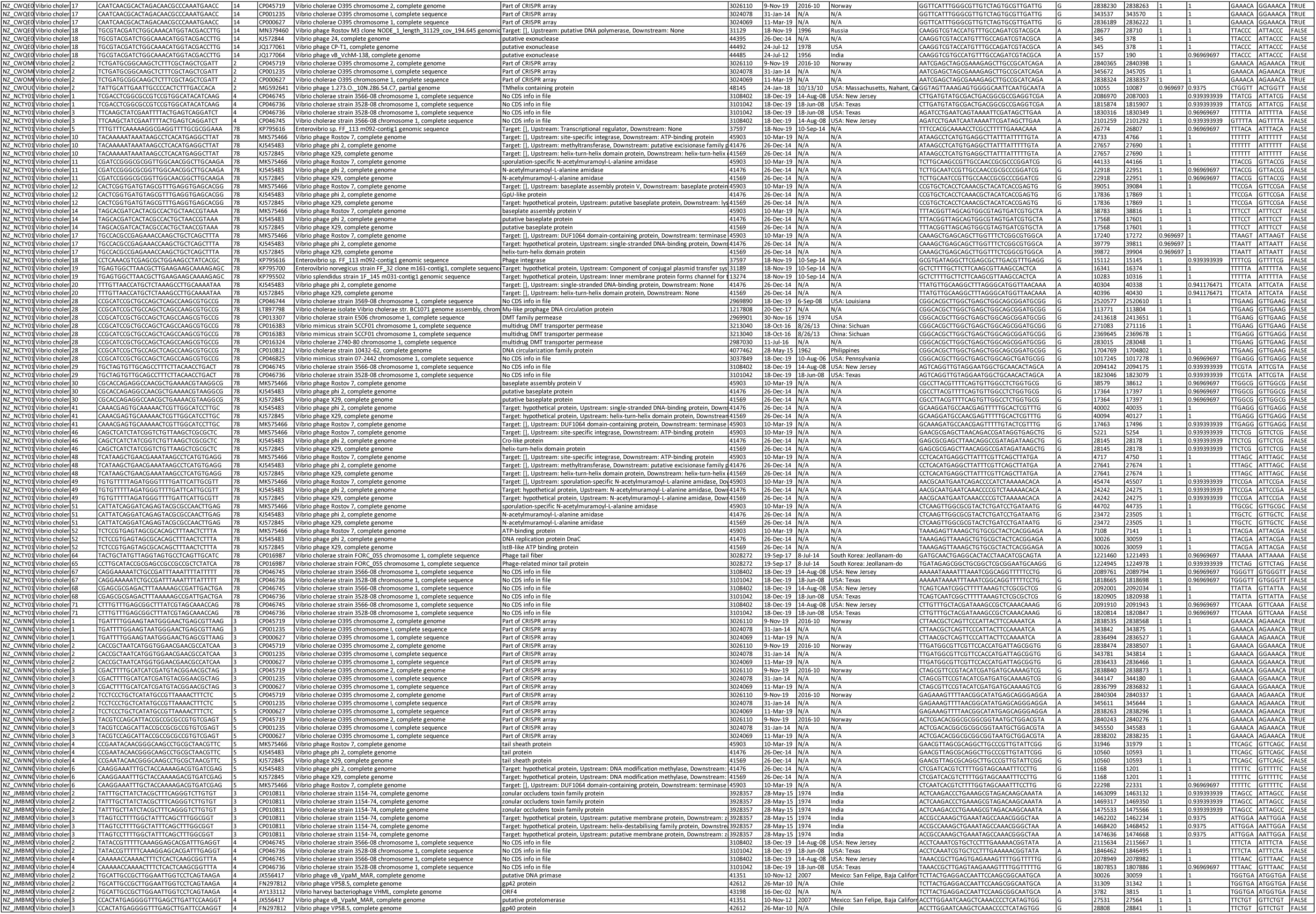
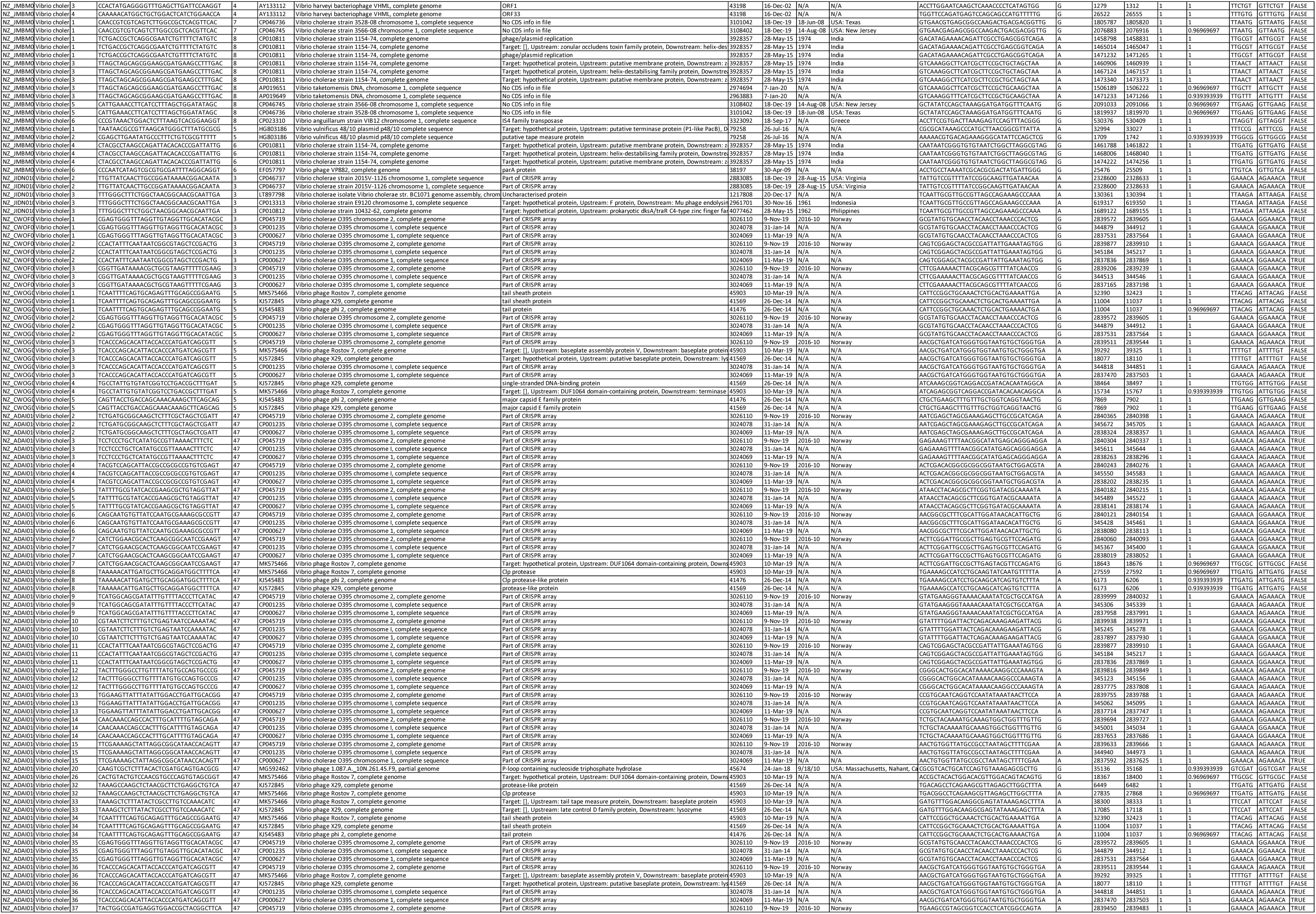
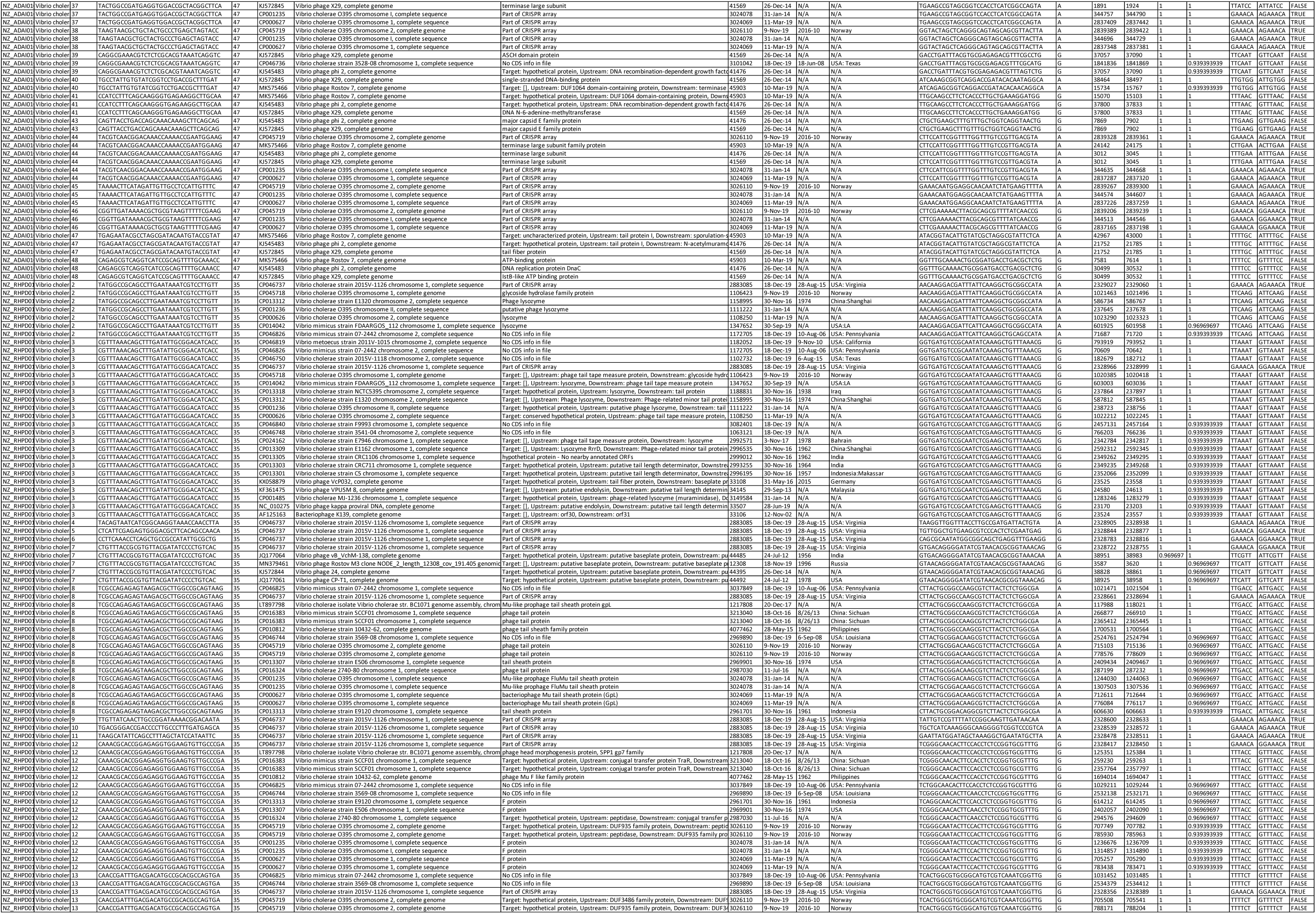
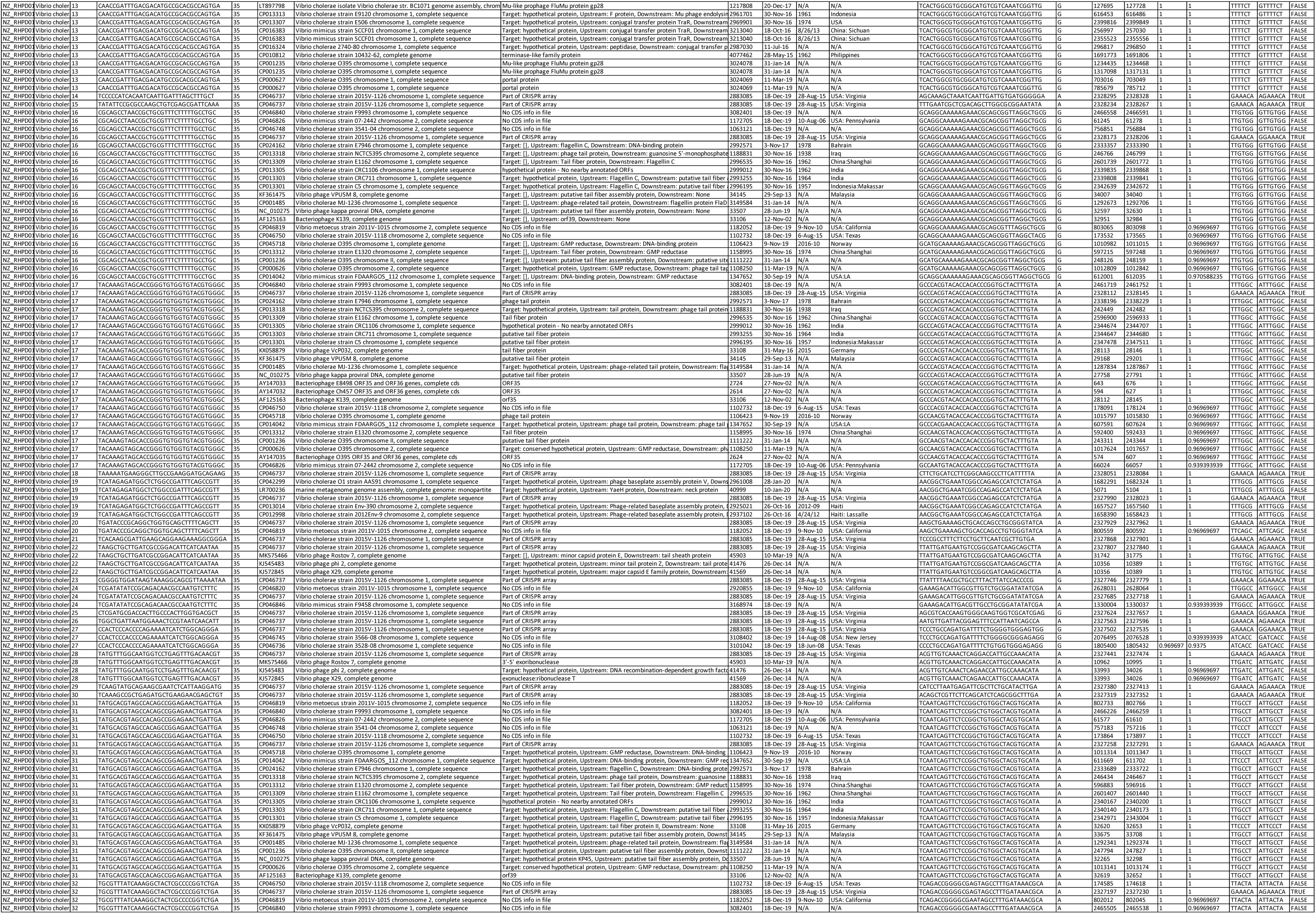
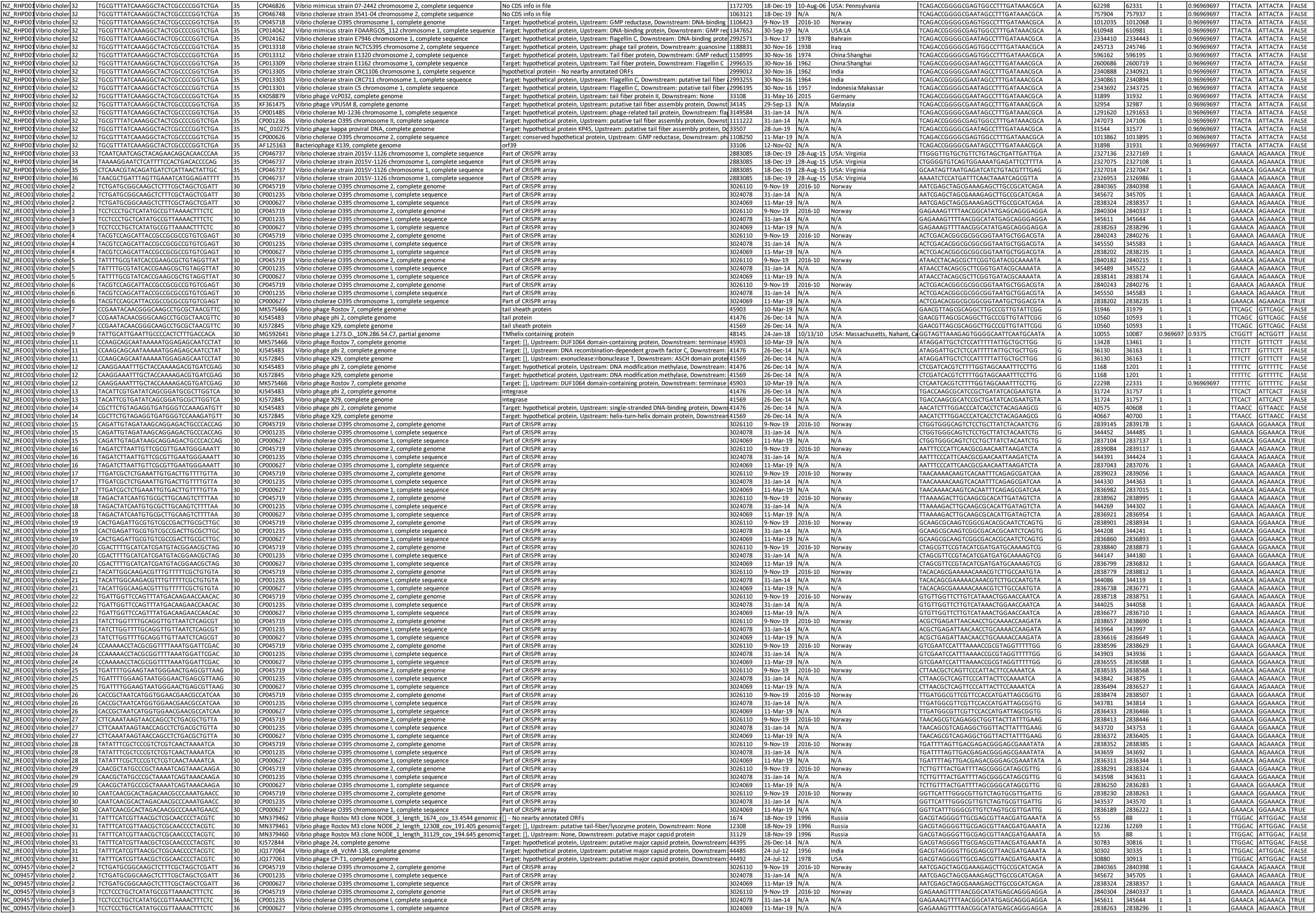
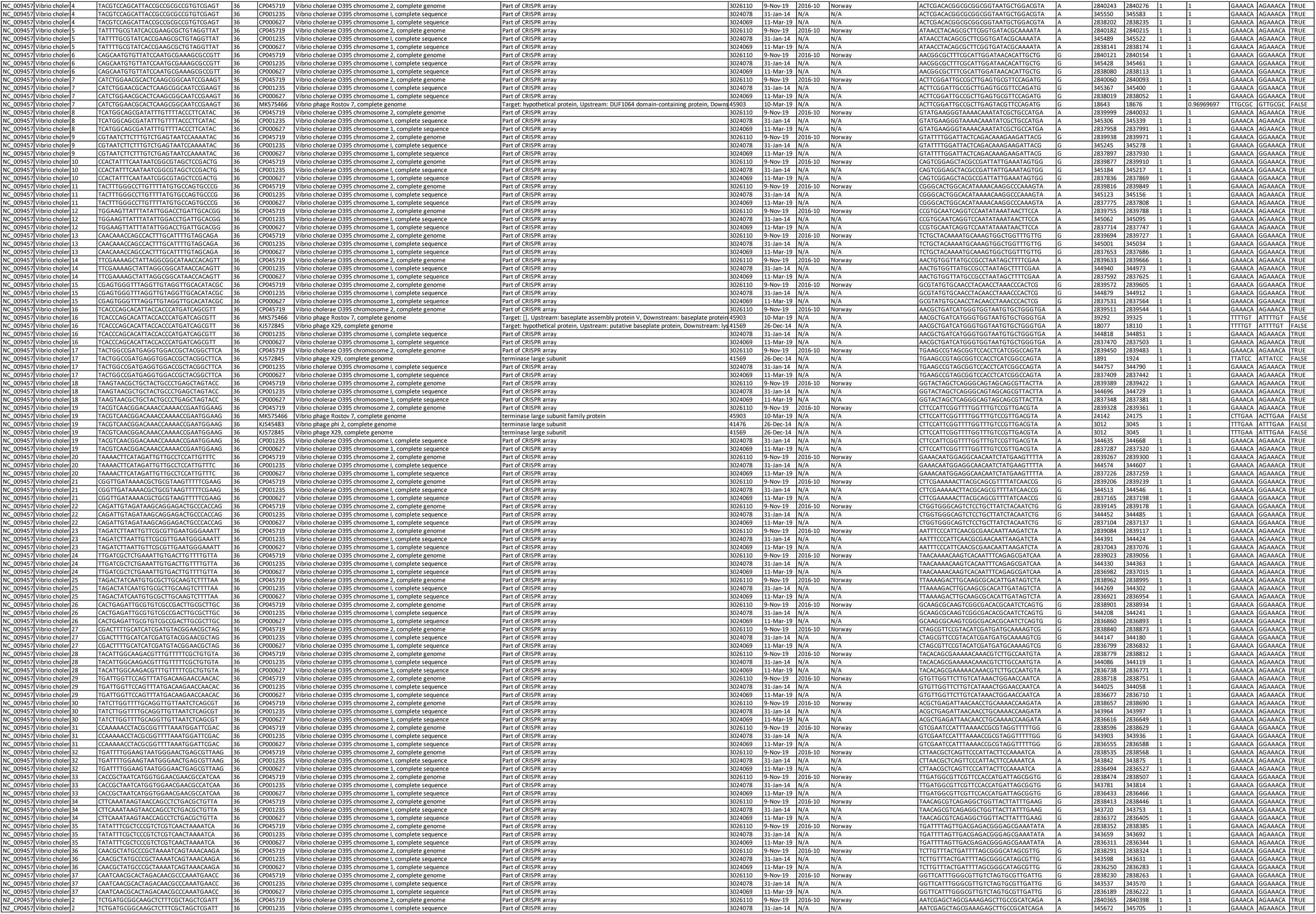
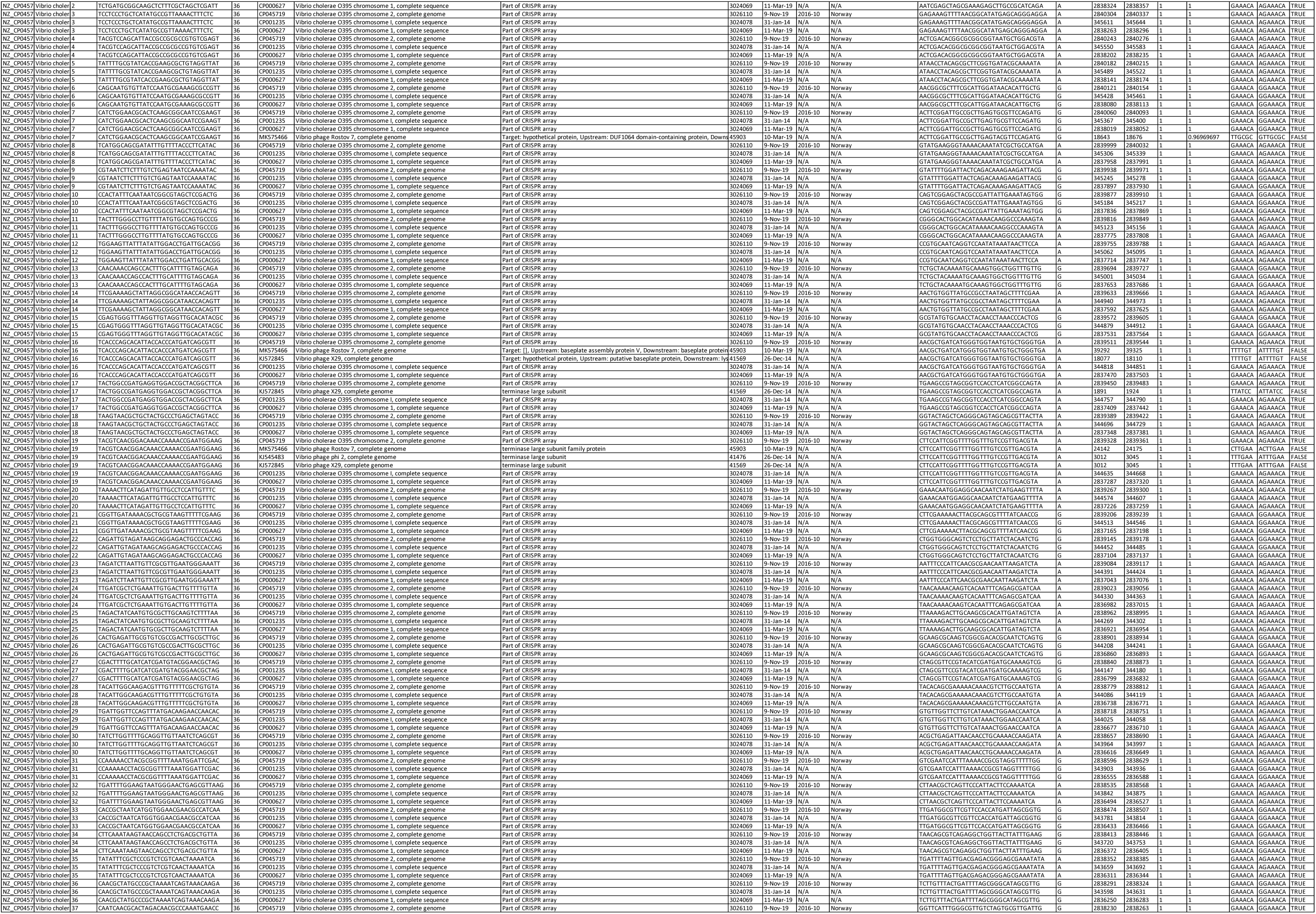
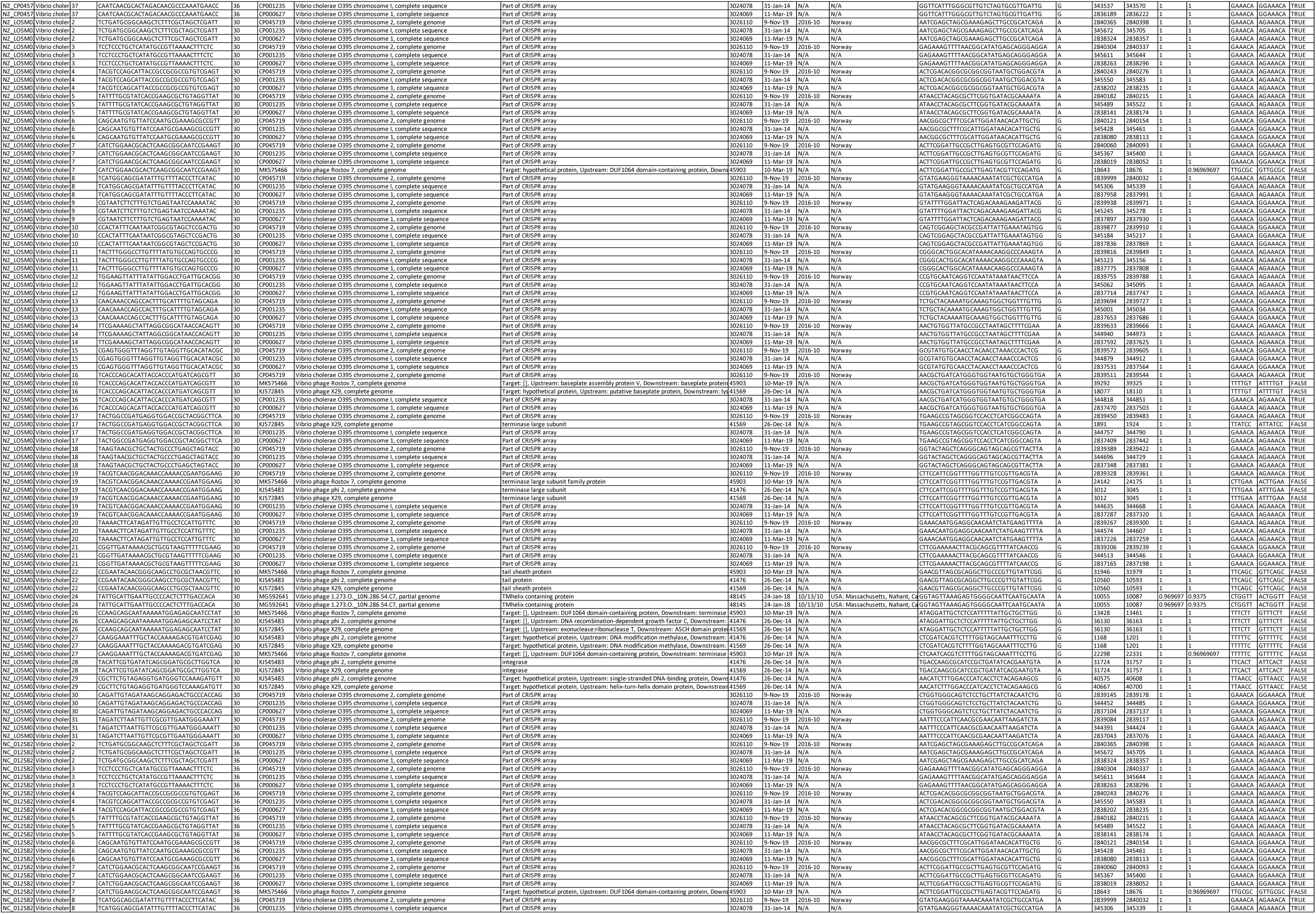
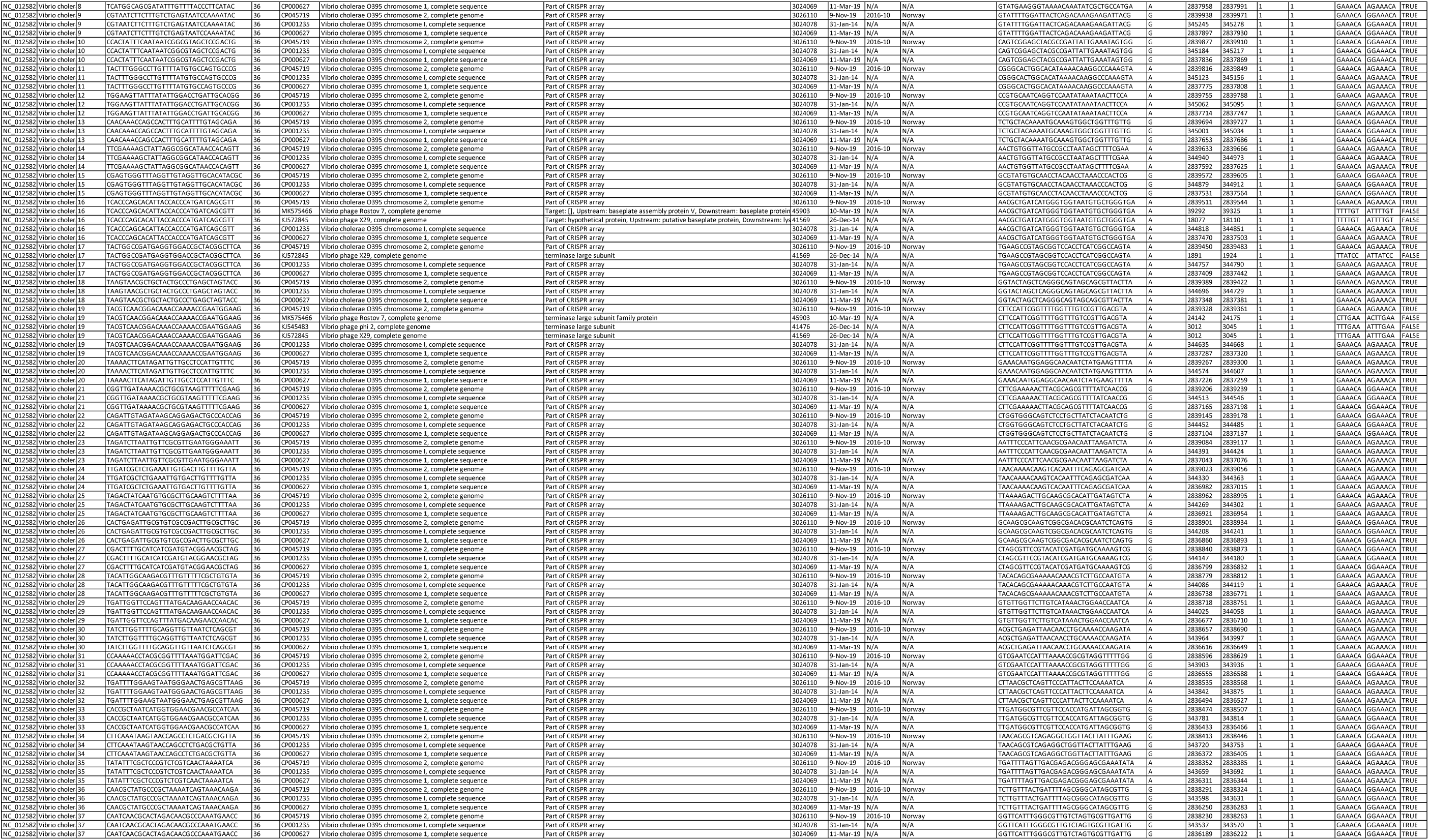
Overview of mined protospacers from identified *V. cholerae* sequences harboring the Type I-E CRISPR/Cas system detected by BLAST to the NCBI non-redundant database.

